# Biomolecular Condensates Can Enhance Homotypic RNA Clustering

**DOI:** 10.1101/2024.06.11.598371

**Authors:** Tharun Selvam Mahendran, Gable M. Wadsworth, Anurag Singh, Ritika Gupta, Priya R. Banerjee

**Affiliations:** Department of Biological Sciences, The State University of New York at Buffalo, Buffalo, NY, 14260, USA; Department of Physics, The State University of New York at Buffalo, Buffalo, NY, 14260, USA

## Abstract

Intracellular aggregation of repeat expanded RNA has been implicated in many neurological disorders. Here, we study the role of biomolecular condensates on irreversible RNA clustering. We find that physiologically relevant, and disease-associated repeat RNAs spontaneously undergo an age-dependent percolation transition inside multi-component protein-nucleic acid condensates to form nanoscale clusters. Homotypic RNA clusters drive the emergence of multiphasic condensate structures, with an RNA-rich solid core surrounded by an RNA-depleted fluid shell. The timescale of the RNA clustering, which accompanies a liquid-to-solid transition of biomolecular condensates, is determined by the sequence features, stability of RNA secondary structure, and repeat length. Importantly, G3BP1, the core scaffold of stress granules, introduces heterotypic buffering to homotypic RNA-RNA interactions and impedes intra-condensate RNA clustering in an ATP-independent manner. Our work suggests that biomolecular condensates can act as sites for RNA aggregation. It also highlights the functional role of RNA-binding proteins in suppressing aberrant RNA phase transitions.

## Introduction

Macromolecular phase separation is a segregative mechanism utilized by living cells to form mesoscale compartments, henceforth referred to as biomolecular condensates^1, 2, 3, 4, 5, 6^. A common feature of many ribonucleoprotein (RNP) condensates, such as the cytoplasmic stress granules, is their dynamic, fluid-like material properties, which enable their facile on-demand formation, dissolution, and rapid macromolecular transport^7^. Experimental and computational approaches probing intra-condensate rheology have revealed that biomolecular condensates are viscoelastic fluids^8, 9, 10, 11^ with material and structural properties that can change over time, termed physical aging^12^. Age-dependent loss in fluidity of condensates can lead to a liquid-to-solid phase transition^12, 13, 14, 15, 16^. Both protein and RNA components can contribute to the physical aging of condensates. In the protein-centric model of condensate aging^17, 18, 19, 20^, phase separation of RNA-binding proteins (RBPs) such as hnRNPA1, FUS, and TDP43 is considered metastable^9, 12, 14, 21, 22, 23, 24^. Hence, condensates formed by these RBPs are susceptible to transition into stable solids, which can either be a glassy material^9^, viscoelastic Kelvin-Voigt solid^12^, or amyloid fiber^23, 25^. Clinically relevant mutations in these RBPs linked to numerous neurodegenerative disorders including amyotrophic lateral sclerosis (ALS)^13, 15, 22, 24^ can accelerate the physical aging of condensates.

Multiple lines of recent evidence also suggest that RNA-driven condensation plays a central role in the formation and regulation of RNP condensates including stress granules^26, 27, 28^ and paraspeckles^29,30^. Furthermore, aberrant intracellular aggregation of GC-rich RNAs with expanded repeats is a hallmark of numerous neurological disorders including Huntington’s disease and ALS^31, 32, 33, 34, 35, 36, 37^. These RNAs form pathological intracellular condensates, which are sensitively dependent on the repeat length and likely involve sequence and structure-specific homotypic RNA-RNA interactions^38, 39, 40^. Homotypic RNA clustering has also been reported to drive mRNA self-assembly in germ granules^41, 42^ and in vitro for RNA homopolymers^26^, indicating condensation of RNA molecules may be ubiquitous in cells. A recent study showed that RNAs have an intrinsic propensity to undergo protein-free phase separation in vitro upon heating with lower critical solution temperatures (LCSTs), driven by desolvation entropy and Mg^2+^- dependent physical cross-linking of RNA molecules^43^. The thermo-responsive phase separation of RNAs can be coupled to an intra-condensate networking transition within the dense phase, referred to as percolation^44, 45, 46^, which engenders long-lived physical cross-linking of RNA chains through nucleobase-specific interactions. Importantly, phase separation and percolation of RNA chains are two distinct transitions, the former being an entropy-driven density transition leading to the formation of phase-separated RNA condensates and the latter being an associative transition mediated by multivalent RNA-RNA interactions. Percolation coupled to phase separation can result in the dynamical arrest of RNAs in the dense phase rendering RNA condensation irreversible. RNA percolation could potentially be exploited in disease conditions to perturb RNP granule dynamics via aberrant RNA clustering, and hence, provides a conceptual framework to model the aggregation landscape of repeat expanded RNAs. These outstanding insights lead to two important questions: (a) do multicomponent RNP condensate microenvironments enhance or suppress RNA percolation, and (b) does intra-condensate RNA percolation contribute to the age-dependent condensate transition from a predominantly liquid to a solid state?

To address these questions, here we employ a designed multi-component condensate system and quantitative microscopy with nanorheology, focusing our attempt to capture key elements of intra-condensate RNA percolation. Our experiments reveal that percolation transitions of physiologically relevant and disease-associated repeat RNAs engender the formation of viscoelastic RNA-rich sub-phases embedded within a fluid-like condensate matrix in an age-dependent manner. The timescale of RNA percolation is tuned by the RNA sequence, secondary structure, and repeat length. Importantly, multivalent RBPs such as G3BP1 can buffer intra-condensate RNA-RNA homotypic interactions, thereby enhancing condensate metastability and delaying the onset of RNA clustering. Overall, our findings suggest that biomolecular condensates can act as sites for RNA clustering and highlight the chaperone-like function of RBPs to buffer the intrinsic capacity of some RNAs to undergo irreversible percolation.

## Results

### RNA aggregation is enhanced in multi-component biomolecular condensates

To study the emergent role of homotypic RNA-RNA interactions within heterotypic protein-nucleic acid condensates, we utilized a model condensate system amenable to quantitative experimental interrogation. This model system consists of an RNA binding motif-inspired multivalent disordered polypeptide [RGG; sequence: (RGRGG)_5_] and a single-stranded nucleic acid [poly-thymine DNA, d(T)_40_]. Phase separation in RGG-d(T)_40_ mixtures is driven by obligate cation-π and electrostatic protein-nucleic acid interactions^8^. Furthermore, the material and physical properties of RGG-nucleic acid condensates are fully tunable via peptide and nucleic acid sequence and length^8, 47^, providing a robust means to tune the condensate microenvironment. Introducing additional components, such as client RNAs featuring specific primary sequence and secondary structures, to this model condensate allows us to systematically probe the effects of compositional complexity and RNA-RNA interaction-driven changes in condensate physical properties (**Fig. 1a**). The rationale for choosing d(T)_40_ over other nucleic acid sequences as a scaffold for our model condensate system is further motivated by a lack of percolation propensity of pyrimidine-based nucleic acids^43^, thereby allowing us to directly probe the percolation behavior of client RNAs. We hypothesize that the introduction of RNAs capable of forming homotypic intra- and inter-molecular contacts within RGG-d(T)_40_ condensates would result in one of two outcomes: the formation of a homogeneous ternary condensate where the RNA fully mixes with the two primary condensate-forming components, RGG and d(T)_40_, or a multiphasic condensate with RNA preferentially partitioning into one of the two co-existing phases. A possible third outcome would be a homogeneous ternary condensate undergoing age-dependent transformation into a multiphasic condensate due to RNA demixing (**Fig. 1a**). In our first set of experiments, we put this idea to the test by utilizing the naturally occurring telomeric repeat-containing RNA^48, 49^ [TERRA, sequence: (UUAGGG)_n_] (**Fig. 1b**), which is known to form G-quadruplex (GQ) structure^50, 51, 52^.

**Figure 1.**
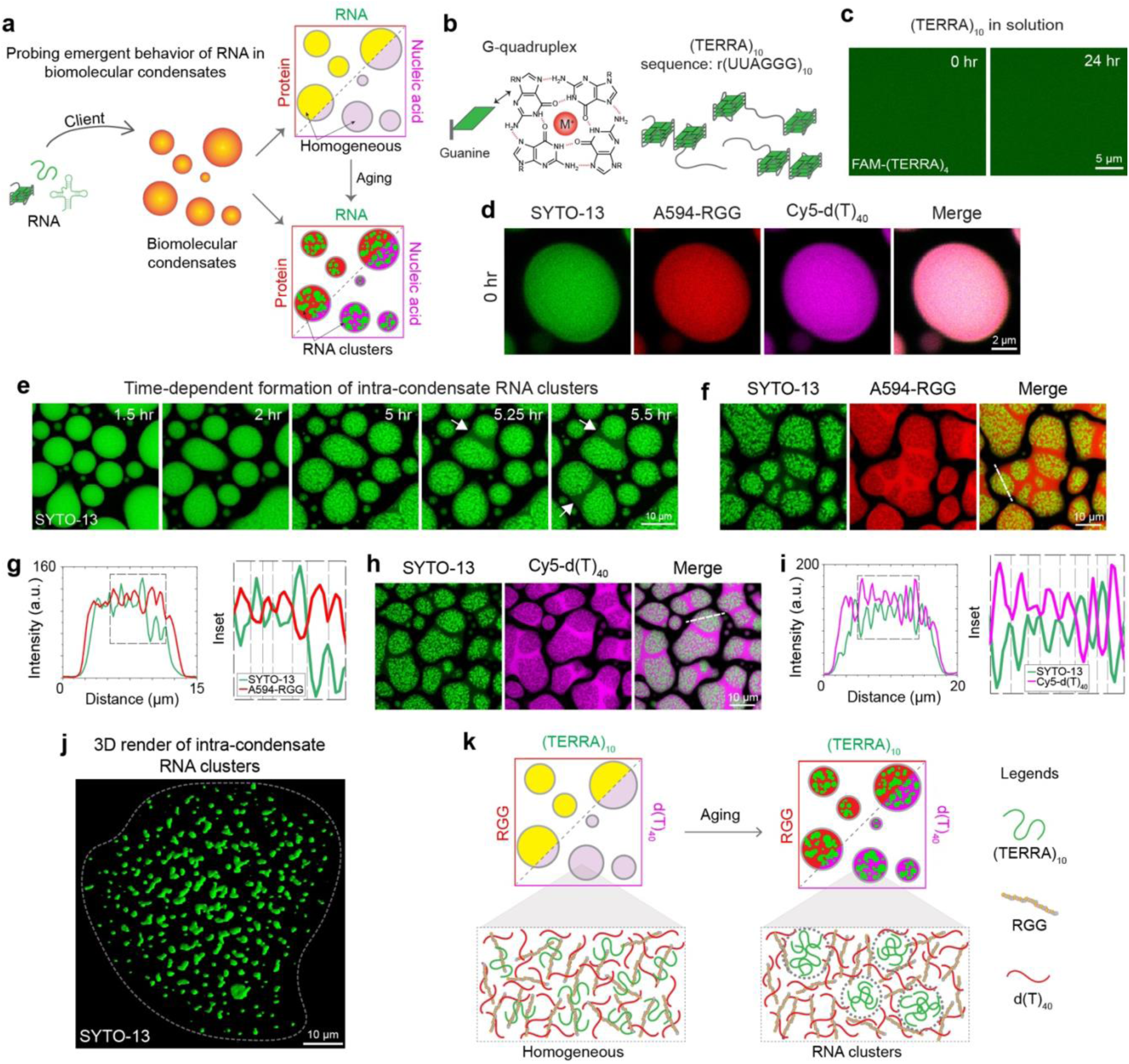
RNA aggregation is enhanced within multi-component biomolecular condensates. **(a)** Depiction of the experimental approach to probing the emergent properties of a client RNA in a heterotypic biomolecular condensate system comprised of RGG and d(T)_40_. **(b)** A top-view schematic of Hoogsteen base-pairing in a G-quartet that forms the structural foundation of (TERRA)_10_. **(c)** In dilute solution, (TERRA)_10_ at 10 mg/ml remains soluble and does not show age-dependent aggregation, as probed by FAM-labeled (TERRA)_4_, over a period of 24 hours at room temperature. **(d)** In multicomponent condensates containing 1.0 mg/ml (TERRA)_10_ (corresponds to 50.7 μM), 5.0 mg/ml RGG, 1.5 mg/ml d(T)_40_ [buffer = 25 mM Tris-HCl (pH 7.5), 25 mM NaCl, 20 mM DTT], all condensate components appear homogeneous within the dense phase at the initial time point (15 minutes after sample preparation) visualized using SYTO-13 (green), Alexa594-RGG (red) and Cy5-d(T)_40_ (magenta). **(e)** (TERRA)_10_ in multicomponent condensates [same composition as (d)] undergoes an age-dependent demixing transition into fractal-like clusters, visualized with SYTO-13. RNA demixing results in a multiphasic architecture of the condensates. Fusion of the shell phase of neighboring condensates is indicated by white arrows (see **Supplementary Video 1**). Localization of (TERRA)_10_ in 18 hours aged condensates [same composition as (d)] is negatively correlated with the localizations of RGG **(f, g)** and d(T)_40_ **(h, i)** as visualized by Alexa594-RGG and Cy5-d(T)_40_. **(j)** Three-dimensional rendering of super-resolution Z-stack images of (TERRA)_10_ clusters from an 18-hour-aged sample [same composition as (d)] visualized with SYTO-13. **(k)** Schematic of age-dependent intra-condensate RNA clustering. In experiments utilizing fluorescently labeled components, the concentration is 250 nM to 500 nM. Each experiment was independently repeated at least three times.

Without multivalent cofactors such as Mg^2+^ ions, (TERRA)_10_ alone remains fully soluble over a period of 24 hours at room temperature (**Fig. 1c**). When introduced as a client to RGG-d(T)_40_ condensates, (TERRA)_10_ preferentially partitions in the dense phase and remains homogeneously distributed at early time points (**Supplementary Fig. 1**) alongside the condensate scaffolding components (**Fig. 1d**). Using a nucleic acid-binding dye (SYTO-13), we next tracked whether the intra-condensate spatial distribution of TERRA changes with condensate age. Remarkably, we find that TERRA undergoes age-dependent demixing inside RGG-d(T)_40_ condensates leading to the formation of RNA clusters (**Fig. 1e**). As a result, a distinct multiphasic architecture emerges that is defined by an RNA-rich inner core surrounded by an RNA-deplete shell. Utilizing fluorescently labeled anti-sense TERRA; we confirmed that the inner SYTO-13-positive sub-phases in aged condensates are indeed formed by TERRA clusters (**Supplementary Fig. 1**). The outer phase of aged condensates displays liquid-like dynamics such as fusion with the outer phase of nearby condensates, which is not observed for the RNA-rich core (**Fig. 1e; Supplementary Video 1**). To elucidate the molecular compositions of these distinct phases, we employ a pairwise imaging approach and identify the localization of each component. The TERRA-enriched clusters within the condensate are significantly depleted of RGG (probed by Alexa594-labeled RGG) and d(T)_40_ [probed by Cy5-labeled d(T)_40_] (**Fig. 1f-i**). Line profiles illustrate the anti-correlation between TERRA and either of these two components (**Fig. 1g, i**). Moreover, the RNA clusters are shown to be quite irregular in morphology (**Fig. 1j**). Replacing d(T)_40_ with r(U)_40_ as a condensate forming component in the ternary system did not alter the age-dependent clustering of TERRA (**Supplementary Fig. 2**). However, we do not observe this time-dependent appearance of microscale RNA clusters in the dense phase of binary condensates composed of TERRA and RGG (**Supplementary Fig. 3**). This observation suggests that the competitive interactions introduced by the nucleic acid component [d(T)_40_] against RGG-(TERRA)_10_ interactions may play a critical role in microscale RNA cluster formation. Upon addition of d(T)_40_ to pre-formed homogeneous RGG-(TERRA)_10_ binary condensates, we observe the emergence of multiphasic condensate architecture with a (TERRA)_10_-rich phase forming the core and the (TERRA)_10_-depleted phase forming the surrounding phase (**Supplementary Fig. 4; Supplementary Video 2**) similar to the ternary system (**Fig. 1e-i**).

To test whether enhanced macromolecular crowding in the dense phase contributes to microscale RNA cluster formation, we titrated the concentration of a molecular crowder, PEG8000, in a solution containing (TERRA)_10_ alone. These samples did not show any signs of RNA aggregation (**Supplementary Fig. 5**). Further, we probed the effect of total RNA concentration on the outcome of RNA cluster formation by titrating the total bulk concentration of (TERRA)_10_ and (TERRA)_4_ in the presence of RGG-d(T)_40_ condensates. We observed RNA clustering with as low as 0.25 mg/ml in either case (**Supplementary Fig. 6**). We also examined whether the increased intra-condensate concentration of RNA can directly lead to cluster formation in the absence of a condensate microenvironment. In the absence of condensates, at 30 mg/ml (TERRA)_4_ concentration, we did not observe any RNA clusters despite the bulk RNA concentration being greater than the lowest dense phase concentration of 25.1 mg/ml at which (TERRA)_4_ clusters are observed in our experiments (**Supplementary Fig. 7**). These observations collectively suggest that age-dependent homotypic RNA clustering (**Fig. 1k**) is an emergent property of the ternary condensates composed of RGG-(TERRA)_10_-d(T)_40_.

### Intra-condensate RNA aggregation is driven by RNA percolation transition

In the absence of proteins, RNA can undergo reversible temperature-dependent phase separation with a secondary percolation transition^43^ (**Fig. 2a**). The percolation transition, which is manifested by homotypic RNA-RNA interactions and sensitively depends on the RNA sequence and secondary structure^43^, dynamically arrests RNA in the dense phase and makes RNA phase separation irreversible. Therefore, RNAs with a strong percolation propensity tend to form irreversible condensates in contrast to RNAs with a weak or no percolation propensity (**Fig. 2a**). We hypothesized that the demixing of (TERRA)_10_ in the dense phase of the multicomponent protein-nucleic acid condensates (**Fig. 1; Supplementary Fig. 2**) stems from a strong percolation propensity of the RNA. To test this idea, we performed temperature-controlled microscopy of (TERRA)_10_ alone in solution. These measurements map the temperature-dependent phase behavior of RNA alone in conditions where confounding effects of time-dependent changes are not observed. We observed that (TERRA)_10_ [1.0 mg/ml RNA in 50 mM HEPES (pH 7.5), 6.25 mM Mg^+2^] remains homogeneous at 20°C but undergoes an irreversible phase transition with a lower cloud point temperature (LCPT) of T = 64.3 ± 1.8°C upon heating. The irreversibility of the (TERRA)_10_ condensates upon cooling to 20°C suggests the formation of a strongly percolated RNA network in the dense phase (**Fig. 2b, c; Supplementary Video 3**). Using a series of Mg^2+^ titrations, we mapped the state diagram of (TERRA)_10_ and observed that the phase separation coupled to percolation behavior of (TERRA)_10_ is extremely sensitive to Mg^2+^ concentration with a very narrow range of Mg^2+^ concentrations (5.75 to 6.5 mM) where thermo-responsive phase separation is experimentally observable (**Fig. 2c**). At Mg^2+^concentrations higher than 6.5 mM, we found that (TERRA)_10_ always exists as percolated clusters even at the lowest temperature tested, 2.0°C (**Fig. 2c**). At temperatures above percolation temperature (T_prc_), (TERRA)_10_ clusters underwent shape relaxation into energetically favorable spherical condensates that persist after cooling, thereby irreversibly trapping the RNA in the condensed state (**Fig. 2d; Supplementary Video 4**). Additionally, temperature-controlled measurements of (TERRA)_8_ and (TERRA)_9_ reveal that the observed phase separation and percolation transitions of the RNA depend on the TERRA repeat number, with higher Mg^2+^ concentrations required for LCST-type transitions of TERRA with fewer number of repeats (**Supplementary Fig. 8**). These results, therefore, reveal a strong percolation propensity of (TERRA)_10_, which is likely to drive its age-dependent aggregation within RGG-d(T)_40_ condensates.

**Figure 2.**
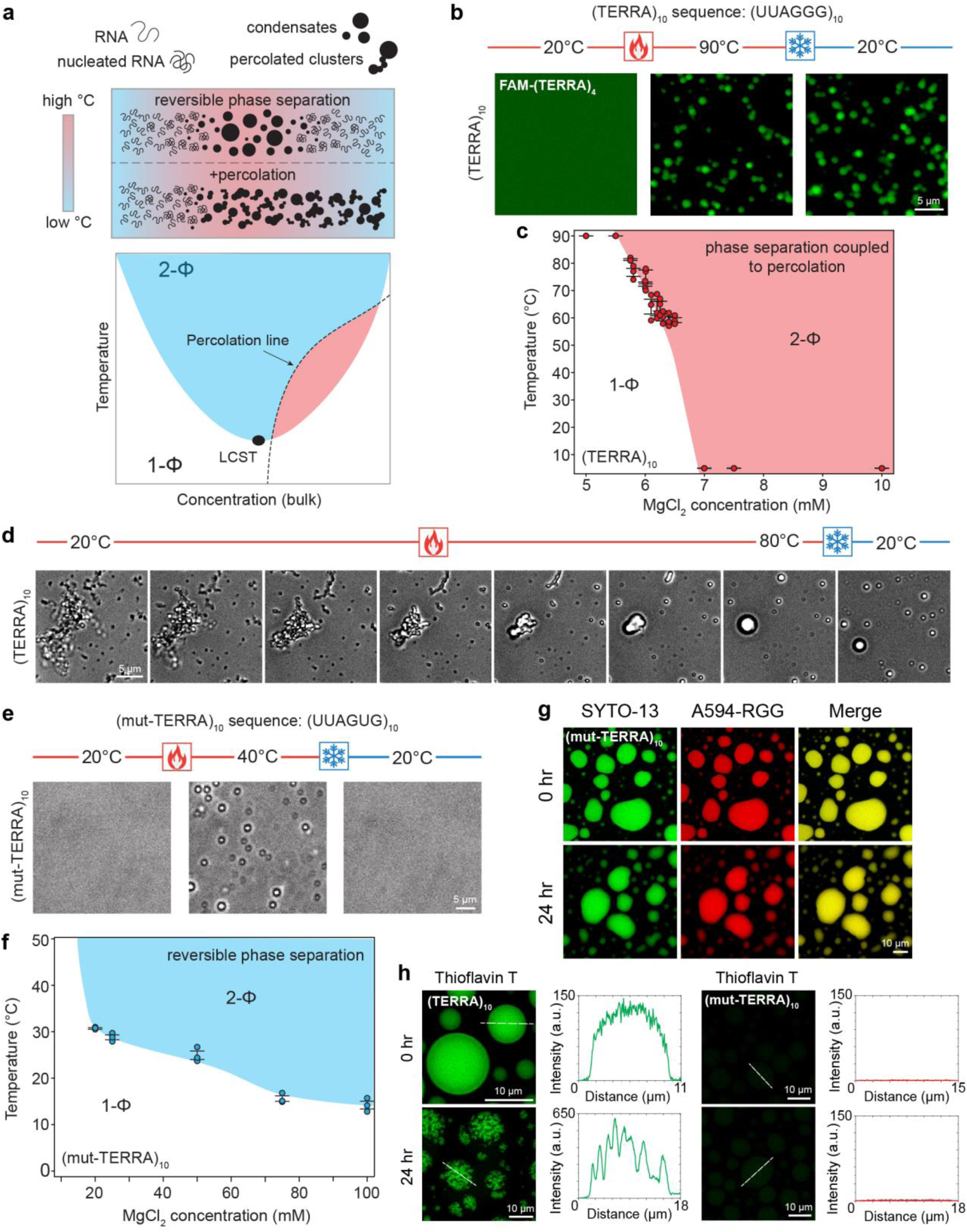
TERRA undergoes phase separation coupled to percolation that can be perturbed by mutations. **(a)** (top) A schematic representing reversible phase separation to form RNA condensates as well as the phase separation coupled to percolation behavior to form irreversible condensates, in response to heating/cooling ramps. (bottom) Depiction of an RNA state diagram, highlighting regions of reversible LCST-type phase separation (blue) and irreversible percolation (red) demarcated by the percolation line (black dashed line). **(b)** (TERRA)_10_ with 6.25 mM Mg^2+^ remains homogeneous in solution at 20°C, as visualized using FAM-(TERRA)_4_. At an elevated temperature, (TERRA)_10_ phase separates into condensates and does not revert to solution upon cooling, indicative of percolated network formation (**Supplementary Video 3**). **(c)** State diagram of (TERRA)_10_ for a set of experiments similar to (b) with titrations of Mg^2+^ concentration. Error bars denote the standard error of the mean (S.E.M.). **(d)** Temperature-controlled microscopy shows that (TERRA)_10_ with 10 mM Mg^2+^ forms percolated clusters at 20°C, which upon heating above the percolation temperature (T_prc_), undergo shape relaxation into spherical condensates that persist when cooled to 20°C (**Supplementary Video 4**). **(e)** Temperature-controlled microscopy of 1 mg/ml (mut-TERRA)_10_ in 50 mM HEPES with 25 mM Mg^2+^ shows reversible RNA phase separation (**Supplementary Video 5**). **(f)** State diagram of (mut-TERRA)_10_ for a set of experiments similar to (e) with titrations of Mg^2+^ concentration. Error bars denote the S.E.M. **(g)** (mut-TERRA)_10_ in RGG-d(T)_40_ condensates does not show intra-condensate RNA percolation. **(h)** Thioflavin T (ThT) staining of (TERRA)_10_ containing RGG-d(T)_40_ condensates shows homogeneous ThT fluorescence at 0 hours and ThT fluorescence within intra-condensate RNA clusters at 24 hours after sample preparation. ThT staining of (mut-TERRA)_10_ containing RGG-d(T)_40_ condensates shows the absence of ThT fluorescence at 0 hours and 24 hours after sample preparation. The concentration of ThT used is 50 μM. The concentration of RNA used for temperature-controlled microscopy measurements is 1 mg/ml [(TERRA)_10_, 50.7 μΜ; (mut-TERRA)_10_, 51.8 μΜ] in 50 mM HEPES (pH 7.5) with the specified Mg^2+^ concentrations. The composition of the ternary condensate system is 1 mg/ml RNA [(TERRA)_10_, 50.7 μΜ; (mut-TERRA)_10_, 51.8 μΜ], 5 mg/ml RGG, and 1.5 mg/ml d(T)_40_ in a buffer containing 25 mM Tris-HCl (pH 7.5), 25 mM NaCl, and 20 mM DTT. In experiments utilizing fluorescently labeled components, the concentration range is 250 nM to 500 nM. Each experiment was independently repeated at least three times.

An important molecular feature of RNA percolation is the contribution of purine-mediated base-pairing and stacking interactions^43^. It is also known that G-rich sequences of RNA can form aggregates in solution^26^. Many of these sequences are also known to form GQ structures with high stability in the presence of monovalent ions^53^. We reasoned that the propensity of TERRA to form GQ structures^50, 52^ may contribute to homotypic RNA clustering via percolation. If true, this can be modulated by TERRA sequence perturbations. To test this idea, we employed a mutant (TERRA)_10_ sequence [(mut-TERRA)_10_] with a G-to-U substitution e.g., (UUAGUG)_10_, which is expected to disrupt the stability of the GQ state^54,55^. We performed temperature-controlled microscopy experiments with (mut-TERRA)_10_ samples and observed that the LCST transition of (mut-TERRA)_10_ requires a substantially higher concentration of Mg^2+^ ions (20 mM) as opposed to 5.75 to 6.5 mM required for (TERRA)_10_ (**Fig. 2b, c** vs. **2e, f**). At 25 mM Mg^2+^, (mut-TERRA)_10_ undergoes reversible phase separation upon heating with an LCPT of 28.8 ± 0.52 °C (**Fig. 2e; Supplementary Video 5**). We note that under the same condition, wild-type (WT) (TERRA)_10_ would exist as percolated clusters at all experimentally accessible temperatures (**Fig. 2c, d**). Percolation of (mut-TERRA)_10_ was not observed at any conditions tested (**Fig. 2f**) demonstrating that the G-to-U substitution attenuates the intermolecular RNA-RNA interactions between TERRA chains. Consistent with the absence of RNA percolation in RNA-only condensates, we find that (mut-TERRA)_10_ does not form age-dependent RNA aggregates in RGG-d(T)_40_ condensates (**Fig. 2g; Supplementary Fig. 9, 10**). This data suggests that intra-condensate RNA percolation is RNA sequence- and structure-specific.

Does the intra-condensate aggregation of TERRA, as reported in **Figure 1e-i**, proceed via a transition from intra-molecular to inter-molecular GQ structures? We tested this idea by utilizing a GQ-selective fluorescent probe Thioflavin T (ThT), which has been previously shown to display fluorescence activation upon binding to GQ structures with high specificity^56, 57, 58^. At an early time-point after the preparation of RGG-d(T)_40_ condensates containing (TERRA)_10_, ThT fluorescence was observed to be uniformly distributed throughout the condensate suggesting that TERRA is likely forming intramolecular GQs within the homogeneous condensate (**Fig. 2h**). Importantly, RGG-d(T)_40_ condensates with (TERRA)_10_ clusters, formed upon aging, displayed ThT positivity, with fluorescence intensities of the clusters being ∼4-fold higher than nascent homogeneous condensates. This observation indicates that the TERRA in these aggregates is also likely to form GQs (**Fig. 2h**). In the case of the (mut-TERRA)_10_, there was no detectable ThT signal under identical imaging conditions at all time points (**Fig. 2h**). This is consistent with the previous reports that G5-to-U substitution in each hexad disrupts the GQ structure of TERRA^54, 55^. Negative control of RGG-d(T)_40_ and RGG-d(T)_40_ condensates containing a non-percolating RNA^43^, poly-uridylic acid (polyU), did not show any measurable ThT signal under identical experimental conditions (**Supplementary Fig. 11**). The lack of the intra-condensate clustering for (mut-TERRA)_10_ could either stem from the loss of GQ structure or the disruption of purine(G)-rich tracts. To probe this further, we designed another mutant sequence, (mut^6GtoU^-TERRA)_10_, wherein alternating G5-to-U substitutions are made across six hexad repeats with the remainder of the repeat sequences left unperturbed (**Supplementary Table 1**). Intriguingly, the (mut^6GtoU^-TERRA)_10_ formed homotypic clusters in RGG-d(T)_40_ condensates in an age-dependent manner, albeit at a slower rate compared to the wild-type (WT) TERRA. Unlike WT (TERRA)_10_, (mut^6GtoU^-TERRA)_10_ containing RGG-d(T)_40_ condensates do not show high ThT fluorescence at homogeneous or in clustered phase (**Supplementary Fig. 12a, b**), suggesting guanine tracts (G-tracts) rather than rGQ structures may constitute the predominant driving force for intra-condensate TERRA percolation into microscale clusters. Finally, intra-condensate (TERRA)_10_ clusters were observed ubiquitously in the presence of a wide variety of monovalent salts (**Supplementary Fig. 13**) that differentially impact the stability of RNA GQ structures, further supporting our proposal that the observed RNA percolation may directly stem from the clustering propensity of purine-rich RNAs^43^.

### Timescale of RNA clustering depends on the TERRA repeat number

The number of guanine tracts (G-tracts) plays important roles in monomeric RNA GQ and multimeric RNA structure formation^59, 60, 61^. We reasoned that time-dependent TERRA aggregation in RGG-d(T)_40_ condensates (**Fig. 1e**) can be tuned by the ability of TERRA to form multi-molecular GQ structures, which is expected to depend on the number of repeat units of TERRA^59, 62^. To test this idea, we titrated the number of TERRA repeat units, [UUAGGG]_n_, where n = 1, 4, 6, and 10, far below the number of repeat units transcribed in cells, 100 to 9000 nucleotides^49, 63^. We observed that all RNAs that could form intramolecular GQs, e.g., n = 4, 6, and 10, showed age-dependent intra-condensate cluster formation with a corresponding timescale correlated inversely with the number of repeats (**Fig. 3a**). (TERRA)_10_ forms microscale aggregates the fastest with a timescale of ∼2 hours, whereas (TERRA)_4_ form clusters at substantially slower rate (∼8 hours) and the clusters are less distinct (**Fig. 3a**). To quantify the relative intra-condensate cluster sizes as a function of time, we employed spatial autocorrelation (SAC) analysis of the confocal fluorescence images of condensates (**Fig. 3a** insets, white-dotted boxes; **Supplementary Fig. 14**). We verified the accuracy of SAC using probe particles of known sizes, which span dimensions above and below the image resolution (see Methods and **Supplementary Fig. 15**). In the absence of clusters larger than the detection limit of the microscope, which is the case for (TERRA)_6_ at 15 minutes after sample preparation, SAC analysis returns the size of spatial fluctuations, 0.28 ± 0.01 μm, which is below the detection limit of the confocal microscope (440 nm; see Methods). However, at the 18-hour timepoint when the RNA clusters in the condensate are detectable, SAC reveals the characteristic size of the RNA clusters being 0.89 ± 0.04 μm (**Fig. 3b**). Employing SAC, we find that the size of RNA clusters increases and the timescale of RNA cluster formation decreases with increasing number of repeat units of TERRA (**Fig. 3c, d**). Simultaneously, we see an increased negative correlation between the intensities of TERRA and the primary components, RGG and d(T)_40_, as a function of condensate age (**Fig. 3e, f; Supplementary Fig. 16, 17**). As clusters become more pronounced with the aging of condensates, the anti-correlation of intensities becomes more apparent (**Fig. 3e, f; Supplementary Fig. 16, 17**). These data suggest that percolated TERRA clusters demix from the RGG and d(T)_40_ in a time-dependent manner. Together with the data presented in the previous section on (mut-TERRA)_10_ failing to form homotypic intra-condensate clusters, these data suggest that repeat number and RNA sequence variations are molecular regulators of age-dependent RNA percolation within biomolecular condensates.

**Figure 3.**
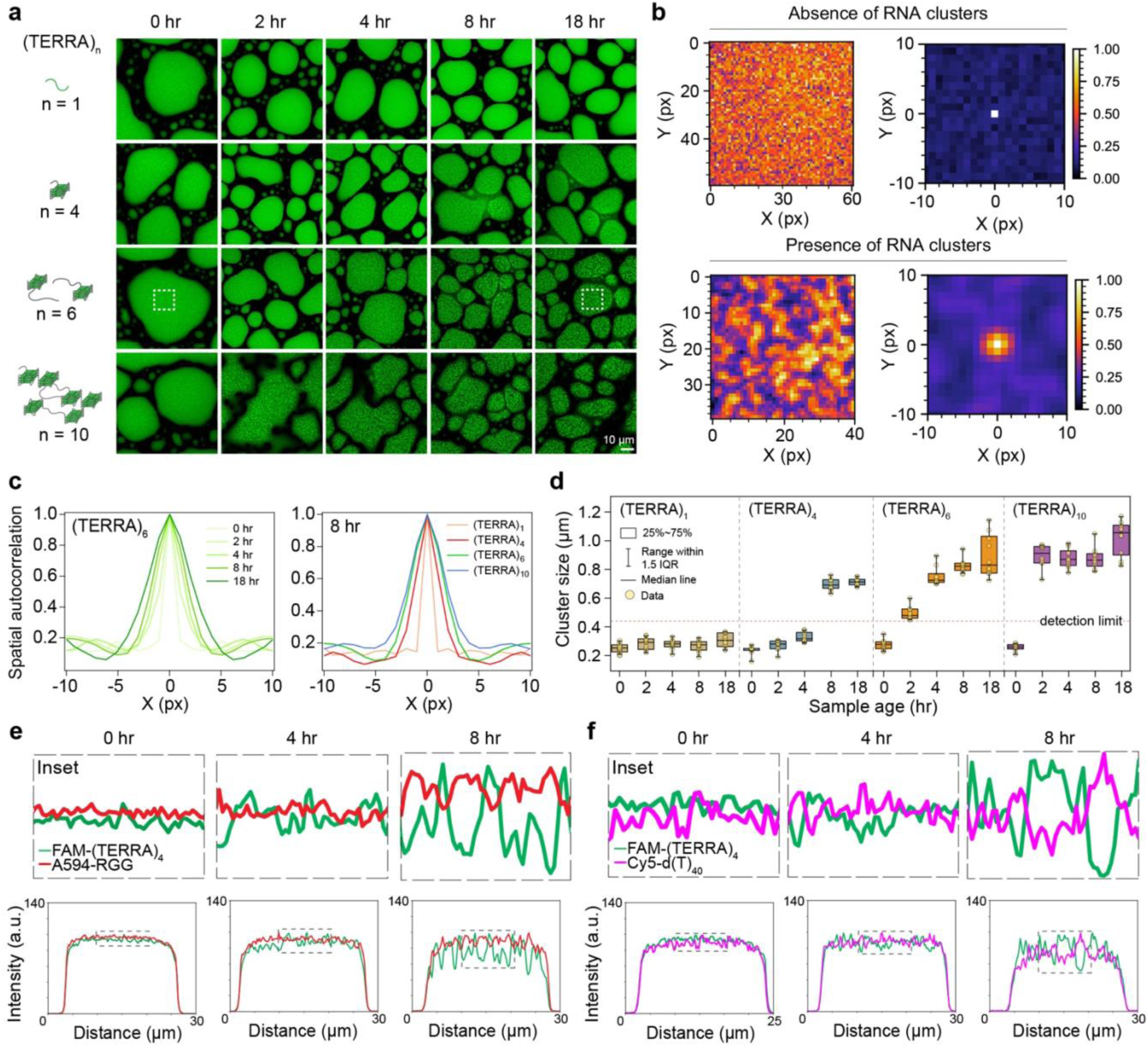
TERRA repeat numbers dictate the timescale of RNA clustering. **(a)** The effect of (TERRA)_n_ repeat number (n) on the timescale of RNA clustering in RGG-d(T)_40_ condensates. TERRA clusters are visualized with SYTO-13 (green). The white box corresponds to analyses done in (b). **(b)** Color map images of (TERRA)_6_ containing RGG-d(T)_40_ condensates as shown in (a) at 15 minutes (above; absence of RNA clustering) and 18 hours (below; presence of RNA clustering) after sample preparation along with corresponding x-y spatial autocorrelation functions from SAC. **(c)** SAC line profiles of (TERRA)_6_ at various time points as shown in (b) (left) and SAC line profiles of (TERRA)_n_ (n= 1, 4, 6, 10) at a sample age of 8 hours (right). **(d)** Cluster sizes derived from SAC of (TERRA)_n_ at increasing sample age. The detection limit of SAC is demarcated (see Methods for further details). Box plot elements are defined within the plot. Pairwise line profile analyses of (TERRA)_4_ images with respect to RGG **(e)** and d(T)_40_ **(f)** as a function of time (for corresponding images, see **Supplementary Fig. 16**). Each line profile shown here is normalized with respect to the maximum intensity value, wherein all values were first offset by the minimum intensity value. The composition of the ternary RGG-d(T)_40_ condensate system is 1 mg/ml RNA [(TERRA)_1_, 521 μΜ; (TERRA)_4_, 127 μΜ; (TERRA)_6_, 84.7 μΜ; (TERRA)_10_, 50.7 μΜ], 5 mg/ml RGG, and 1.5 mg/ml d(T)_40_ in a buffer containing 25 mM Tris-HCl (pH 7.5), 25 mM NaCl, and 20 mM DTT. In experiments utilizing fluorescently labeled components, the concentration range is 250 nM to 500 nM. Each experiment was independently repeated at least three times.

### Repeat expanded RNAs form intra-condensate clusters in a length-dependent manner

Intracellular RNA aggregation has been widely implicated in several diseases primarily in the category of trinucleotide repeat expansion disorders such as Huntington’s disease (CAG repeat), Fragile-X syndrome (CGG repeat), and myotonic dystrophy (CUG repeat)^37, 64, 65, 66, 67, 68, 69^. One characteristic of these disorders is that pathology occurs at a range of repeat lengths beyond what is found under normal circumstances. Further, the threshold number of repeats associated with pathological outcomes varies with the repeat RNA sequence. For example, the number of r(CAG) repeats required for disease onset is ∼15 to 20 repeats less than that required for r(CUG) (**Fig. 4a**). This implies that the cellular machinery can better tolerate r(CUG) repeats than it can with r(CAG) repeats^64, 65, 66, 67, 68^.

**Figure 4.**
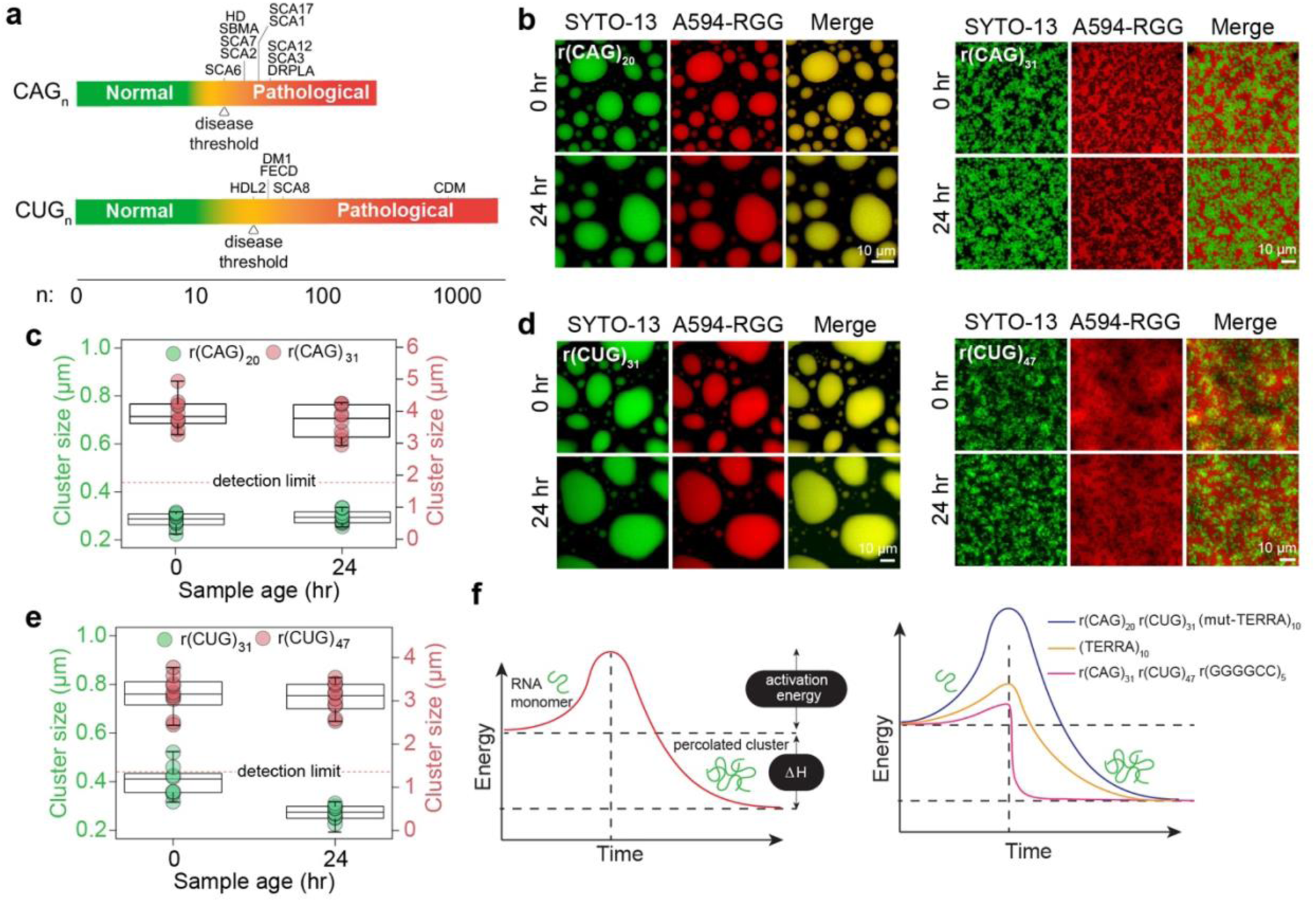
Repeat expanded RNAs form intra-condensate clusters in a length-dependent manner. **(a)** Schematic of disease-associated triplet RNA repeat expansions highlighting the threshold number of RNA repeats (n) that correspond to healthy states versus pathological states. Diseases are listed in an order according to the minimum RNA repeat length required for disease onset. The disease threshold for each GC-rich repeat RNA is marked according to the lowest repeat length linked to disease onset^64, 65, 66^. Disease abbreviations are as follows. SCA6: Spinocerebellar Ataxia (SCA) Type 6, HD: Huntington’s disease, SBMA: Spinal and Bulbar Muscular Atrophy, SCA7: SCA Type 7, SCA2: SCA Type 2, SCA17: SCA Type 17, SCA1: SCA Type 1, SCA12: SCA Type 12, SCA3: SCA Type 3, DRPLA: Dentatorubral-Pallidoluysian Atrophy, HDL2: Huntington’s Disease-Like 2, DM1: Myotonic Dystrophy Type 1, FECD: Fuchs’ Endothelial Corneal Dystrophy, SCA8: SCA Type 8, CDM: Congenital Myotonic Dystrophy. **(b)** r(CAG)_20_ containing RGG-d(T)_40_ condensates remain homogeneous and do not form clusters with time (left) whereas r(CAG)_31_ containing condensates form microscale RNA clusters. The 0-hour image was acquired after 15 minutes of sample preparation (right). **(c)** Cluster sizes derived from SAC corresponding to (b). With respect to the left y-axis, the detection limit of SAC is demarcated (see Methods for further details). The detection limit of the right y-axis is not shown, as the data is significantly above the limit. **(d)** r(CUG)_31_ containing RGG-d(T)_40_ condensates remain homogeneous for 24 hours whereas r(CUG)_47_ containing condensates form microscale RNA clusters. The 0-hour image was acquired after 15 minutes of sample preparation. **(e)** Cluster sizes derived from SAC corresponding to (d). With respect to the left y-axis, the detection limit of SAC is demarcated (see Methods for further details). The detection limit of the right y-axis is not shown, as the data is significantly above the limit. **(f)** A schematic showing the hierarchy of activation energy barrier for an RNA monomer to form percolated RNA clusters. All box plot elements are defined similarly to Fig. 2d. The composition of the ternary RGG-d(T)_40_ condensate system is 1 mg/ml RNA [0.45 mg/ml in the case of r(CUG)_47_; r(CAG)_20_, 51.2 μM; r(CAG)_31_, 33 μM; r(CUG)_31_, 33.8 μM; r(CUG)_47_, 10 μM], 5 mg/ml RGG, and 1.5 mg/ml d(T)_40_ in a buffer containing 25 mM Tris-HCl (pH 7.5), 25 mM NaCl, and 20 mM DTT. In experiments utilizing fluorescently labeled components, the concentration range is 250 nM to 500 nM. Each experiment was independently repeated at least three times.

Motivated by our results of enhanced TERRA clustering as a function of increasing repeat number, we interrogated the clustering propensity of two repeat RNAs, r(CAG)_n_ and r(CUG)_n_, with repeat lengths higher and lower than the pathological limit using our model condensate system. Typically, r(CAG) repeat lengths lower than 21 are found under normal circumstances, however, repeat lengths exceeding this number may be considered to be at the intermediate-high risk scale of being most likely to attain diseases such as spinocerebellar ataxia and Huntington’s disease (**Fig. 4a**). We observed that r(CAG)_20_ stays predominantly homogeneous in the dense phase of RGG-d(T)_40_ condensates at all ages till 24 hours, the end time of our experiments. The mean RNA cluster size is estimated to be 0.28 ± 0.01 μm, which is below the detection limit of the confocal microscope (440 nm; see Methods) (**Fig. 4b, c; Supplementary Fig. 18**). Strikingly, r(CAG)_31_, which corresponds to a pathological number of repeats, spontaneously demixes into fractal-like clusters prior to sample imaging (15 minutes post-preparation) (**Fig. 4b, c; Supplementary Fig. 18**). In these samples, r(CAG)_31_ clusters are extensively percolated with cluster sizes that are nearly 4-fold larger (3.94 ± 0.14 μm) than those formed by TERRA at the same concentration (**Fig. 4c**). Reduced r(CAG)_31_ concentration reduces the intra-condensate cluster size, but the RNA aggregation timescale appears to be independent of r(CAG)_31_ concentration (**Supplementary Fig. 19**).

We next examined intra-condensate percolation of r(CUG)_n_ for n = 31. Similar to r(CAG)_20_, we did not observe intra-condensate r(CUG)_31_ clustering. The estimated cluster size is 0.40 ± 0.02 μm, which is below the detection limit of the confocal microscope (440 nm; see Methods) (**Fig. 4d, e; Supplementary Fig. 18**). In stark contrast, r(CUG)_47_ forms microscale clusters within 15 minutes after sample preparation (**Fig. 4d, e; Supplementary Fig. 18**). We also tested intra-condensate clustering behavior of a hexanucleotide repeat expanded RNA, r(GGGGCC)_n_, that is associated with ALS and frontotemporal dementia (FTD)^59, 70, 71^. We observed spontaneous r(GGGGCC)_5_ clustering in RGG-d(T)_40_ within 15 minutes after sample preparation (**Supplementary Fig. 20**). Importantly, r(GGGGCC)_5_ clusters showed ThT positivity but not r(CAG)_20_ or r(CAG)_31_ clusters (**Supplementary Fig. 21**). This is consistent with the ability of r(GGGGCC)_5_ to form GQ structures^59^. Overall, the distinct differences in the intra-condensate percolation behavior of each of these RNAs and TERRA of various lengths suggest a heuristic framework where a sequence and length-specific activation energy barrier dictates the timescale of intra-condensate RNA clustering (**Fig. 4f**, left). In this picture, r(CAG)_20_, r(CUG)_31_, and (mut-TERRA)_10_ molecules have a substantially higher energy barrier to undergo intra-condensate percolation as compared to (CAG)_31_, r(CUG)_47_, and r(GGGGCC)_5_, whereas (TERRA)_10_ features an intermediate activation energy barrier (**Fig. 4f**, right).

### Intra-condensate RNA clustering accompanies a liquid-to-solid phase transition

What are the material properties of condensates containing percolated RNA clusters? Time-lapse microscopy suggests that the shell phase of the aged condensates containing TERRA clusters can undergo fusion, signifying liquid-like properties (**Fig. 1e, Supplementary Video 1**). However, these condensates are in contact with the glass surface, which can influence their fusion kinetics substantially^72^. To examine the dynamical behavior of (TERRA)_10_ containing RGG-d(T)_40_ condensates quantitatively as a function of age, we employed controlled condensate fusion using optical tweezers^73, 74^. We trapped two condensates using a dual-trap optical tweezer and initiated condensate fusion while recording the fluorescence of FAM-(TERRA)_4_ and Cy5-d(T)_40_ simultaneously. At 20 minutes after the condensate preparation, when both TERRA and d(T)_40_ are uniformly distributed throughout the droplets, they undergo fusion with a fusion relaxation time (τ) of 18.40 ± 0.02 ms/μm. The components were homogeneously mixed after the condensate coalescence was completed (**Fig. 5a; Supplementary Video 6**). However, for condensates aged for 150 minutes, distinct RNA clusters were visible within condensates (**Fig. 5a; Supplementary Video 7**). These aged condensates containing RNA clusters were still able to fuse indicating that the condensate shell is dynamic and behaves as a terminally viscous liquid. Force relaxation analysis showed that fusion of the outer shell of aged condensates occurs more rapidly with a fusion relaxation time of 3.87 ± 0.06 ms/μm (**Fig. 5a**). However, the irregular RNA clusters within the condensate remained demixed at all conditions, indicating that they are in a different material state than RGG-d(T)_40_ rich phase (**Fig. 5a**). We tested this directly by designing a condensate dissolution assay, which would preferentially dissolve the non-percolated components (**Supplementary Fig. 22a**). We observed that addition of 0.5 μl of 5 M NaCl is sufficient to rapidly dissolve condensates formed by RGG-d(T)_40_ (**Supplementary Fig. 22b**). However, NaCl addition to aged RGG-d(T)_40_ condensates containing TERRA clusters showed only partial dissolution where the RGG-d(T)_40_ rich shell phase dissolved immediately but the RNA clusters persisted (**Fig. 5b; Supplementary Videos 8, 9**). Notably, freshly prepared (TERRA)_10_ containing RGG-d(T)_40_ condensates lacking RNA clusters (15 minutes post preparation) were observed to dissolve completely by NaCl treatment (**Supplementary Fig. 22c; Supplementary Video 10)**. We also probed the translational mobility of each component in the condensate as a function of age by fluorescence recovery after photobleaching (FRAP; **Fig. 5c**). We chose (TERRA)_4_ as a representative RNA for these measurements which forms intra-condensate clusters by 8 hours (**Fig. 3a, c**). FRAP analysis reveals that at early time points before the onset of visible RNA clusters (< 4 hours), all three components are relatively dynamic (**Fig. 5c; Supplementary Fig. 23; Supplementary Videos 11, 12, 13**). However, FRAP traces of TERRA, but not RGG and d(T)_40_, show a progressive drop in recovery with condensate age (**Fig. 5c; Supplementary Fig. 23; Supplementary Videos 14, 15, 16**).

**Figure 5.**
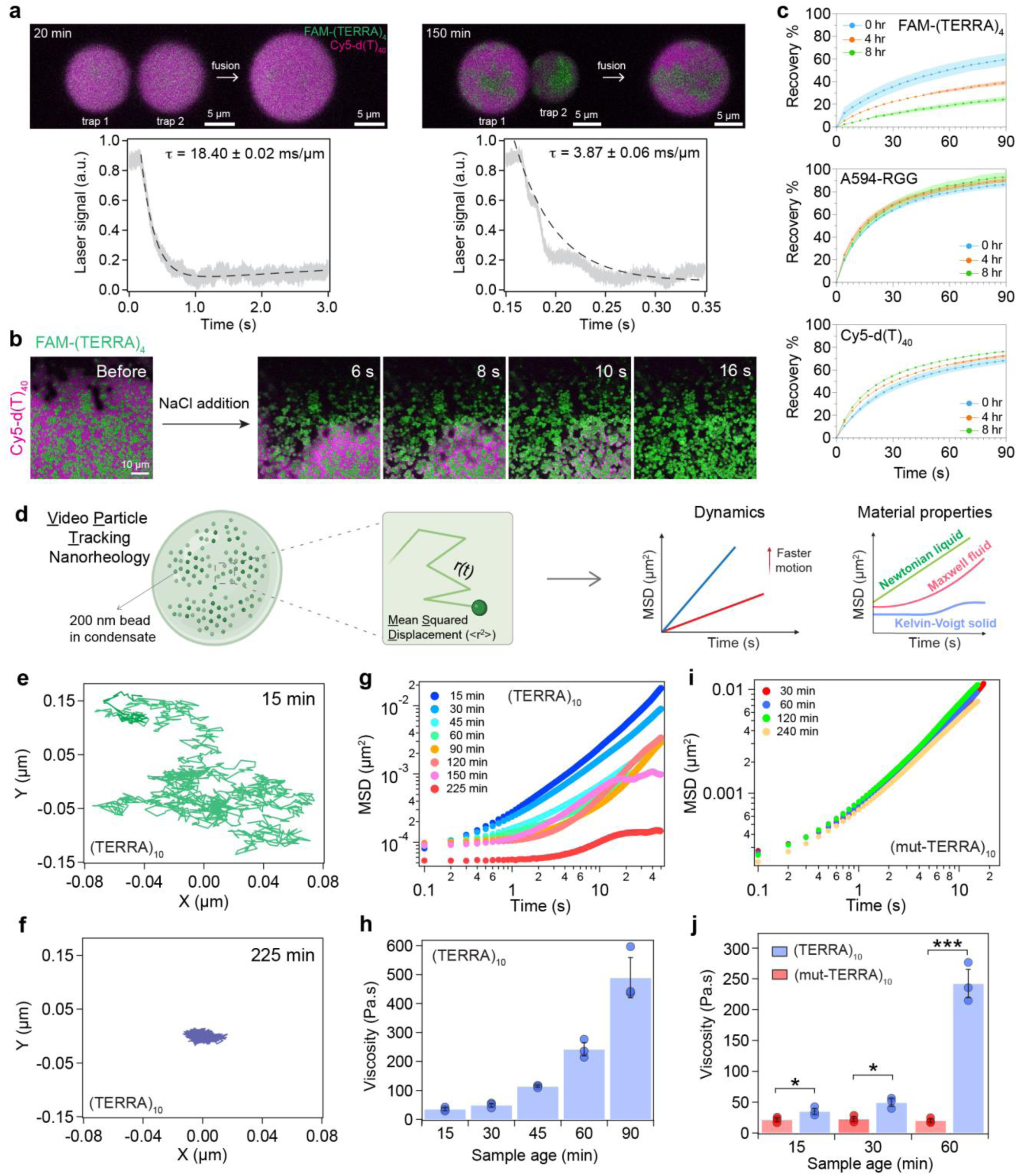
Intra-condensate RNA clustering accompanies a liquid-to-solid transition. **(a)** (top) Optical tweezer-mediated fusion of (TERRA)_10_ containing RGG-d(T)_40_ condensates at two different time points, as indicated, after sample preparation (see **Supplementary Videos 6, 7**). (bottom) Corresponding force relaxation profiles and the estimated fusion relaxation times (error represents ±1 standard deviation). **(b)** The addition of 0.5 μl of 5 M NaCl to (TERRA)_10_ containing RGG-d(T)_40_ condensates at 6 hours after sample preparation shows the dissolution of Cy5-d(T)_40_-rich shell phase (magenta) but not the (TERRA)_10_ clusters (green) (see **Supplementary Fig. 22; Supplementary Video 8**). **(c)** FRAP recovery profiles of FAM-(TERRA)_4_, Alexa 594-RGG, and Cy5-d(T)_40_ in (TERRA)_4_ containing RGG-d(T)_40_ condensate system with age (**Supplementary Fig. 23; Supplementary Videos 11-16**). Shaded regions in each plot signify the S.E.M. **(d)** Schematic of video particle tracking (VPT) nanorheology using beads passively embedded inside condensates, which is used to estimate their mean squared displacements (MSD) to ascertain condensate material properties. Created with BioRender.com. Individual bead trajectories at 15 minutes **(e)** and 225 minutes **(f)** inside (TERRA)_10_ containing RGG-d(T)_40_ condensates. **(g)** MSDs of beads inside (TERRA)_10_ containing RGG-d(T)_40_ condensates as a function of condensate age. **(h)** The corresponding terminal viscosities are reported. Error bars denote the standard deviation. **(i)** MSDs of beads inside (mut-TERRA)_10_ containing RGG-d(T)_40_ condensates as a function of condensate age. **(j)** The corresponding terminal viscosities are reported in comparison to that of (TERRA)_10_ containing RGG-d(T)_40_ condensates as a function of condensate age. Statistical significance was determined by performing a two-sided Student’s *t*-test (* means p<0.05, ** means p<0.01, *** means p<0.001) between viscosities of (TERRA)_10_ and (mut-TERRA)_10_ condensates; the p-values determined at sample ages of 15 minutes, 30 minutes, and 60 minutes are 0.0423, 0.0103, and 0.0003, respectively. Error bars denote the standard deviation. The composition of the ternary condensate system is 1 mg/ml RNA [(TERRA)_10_, 50.7 μΜ; (TERRA)_4_, 127 μΜ], 5 mg/ml RGG, and 1.5 mg/ml d(T)_40_ for the optical tweezer and FRAP experiments; 0.5 mg/ml RNA [(TERRA)_10_, 25.4 μΜ], 2.5 mg/ml RGG, and 0.75 mg/ml d(T)_40_ for the condensate dissolution experiments; and 2 mg/ml RNA [(TERRA)_10_, 101 μΜ; (mut-TERRA)_10_, 104 μΜ], 10 mg/ml RGG, and 3.0 mg/ml d(T)_40_ for the VPT measurements. Buffer composition for all experiments is 25 mM Tris-HCl (pH 7.5), 25 mM NaCl, and 20 mM DTT. In experiments utilizing fluorescently labeled components, the concentration range is 250 nM. All measurements were independently repeated at least three times.

The reduced translational mobility of TERRA could result from the formation of dynamically arrested intra-condensate RNA networks, driving a liquid-to-solid phase transition. To directly probe the material properties of condensates, we employed video particle tracking (VPT) nanorheology using 200 nm fluorescently labeled beads^8^ (**Fig. 5d**). The mean squared displacement (MSD) profiles of the probe particles have distinct characteristics for Newtonian liquids, viscoelastic fluids with terminal viscous behavior, and Kelvin-Voigt solids with terminal elastic behavior^12, 75^ (**Fig. 5d, right**). VPT measurements reveal that freshly prepared RGG-d(T)_40_ condensates containing (TERRA)_10_ display material properties similar to a Maxwell fluid^9^ with a terminal viscosity of 35.2 ± 4.85 Pa.s (**Fig. 5e-h; Supplementary Video 17**). Upon physical aging, the probe particles within condensates are caged as evidenced by a narrower spread of the particle trajectory (**Fig. 5f**). Correspondingly, the ensemble-averaged MSD profiles showed dramatic differences at times beyond the emergence of RNA clusters, e.g., t > 60 minutes (**Fig. 5g**). We find that this is an emergent property of the (TERRA)_10_ containing RGG-d(T)_40_ condensate system since the two-component condensate system composed of RGG and d(T)_40_ does not show time-dependent changes in MSD profiles (**Supplementary Fig. 24**). Conversely, binary condensates of RGG and (TERRA)_10_ are highly viscoelastic and show time-dependent changes in MSD profiles of probe particles, albeit they do not exhibit microscale RNA cluster formation (**Supplementary Fig. 25; Supplementary Fig. 3**). Further measurements of MSD profiles in RGG-d(T)_40_ condensates containing (TERRA)_10_ revealed a dramatic arrest of the probe particles at time points, t = 30 and 45 minutes, at which the microscale RNA clusters were not clearly visible (**Fig. 5g**; **Supplementary Fig. 26; Supplementary Video 18**). The dynamical slowdown of probe particles prior to the emergence of microscale RNA aggregates may indicate the formation of nanoscale pre-percolation clusters of TERRA^76^. Upon further aging, the ensemble-averaged MSDs show a plateauing behavior at longer lag times indicating a terminally elastic response reminiscent of Kelvin-Voigt solids^12^ (**Fig. 5g**; observation time ≥ 150 minutes). This may stem from the confined diffusion of beads due to the onset of percolated RNA clusters (**Fig. 5f**). The estimated terminal viscosity of the condensates at 90 minutes is 489.3 ± 69.3 Pa.s (**Fig. 5h**), which is an order of magnitude higher than condensates at 15 minutes. At 150 minutes after sample preparation, the bead motions were completely arrested, which is a characteristic property of terminally solid material^12^ (**Supplementary Video 19**).

We further performed nano-rheology measurements with (mut-TERRA)_10_ (**Fig. 2e, f**), which lacks the ability to undergo intra-condensate percolation (**Fig. 2g; Supplementary Fig. 9, 10**). We observed that (condensate age ∼15 minutes) RGG-d(T)_40_ condensates containing (mut-TERRA)_10_ behave as a Maxwell fluid with viscosity 21.5 ± 3.1 Pa.s, which is slightly lower than WT (TERRA)_10_ (**Fig. 5i, j; Supplementary Fig. 27**). Importantly, no discernable changes in viscoelastic properties were observed for these condensates over the same period (4 hours; **Fig. 5i, j; Supplementary Fig. 27**) where the WT TERRA containing condensates undergo complete dynamical arrest. These results show that selectively inhibiting the percolation ability of the RNA through sequence perturbations abrogates RNA percolation and in turn, impedes the age-dependent condensate dynamical arrest.

### RNA-binding protein G3BP1 inhibits intra-condensate RNA clustering

In cells, RNP granules are comprised of diverse RNA and protein species with an assortment of sequence-specific and non-specific interactions^77, 78, 79, 80^. Based on our results on age-dependent intra-condensate RNA clustering, we reasoned that introducing additional components in these condensates that can compete with homotypic RNA-RNA interactions may inhibit RNA cluster formation. We tested this idea first by employing an anti-sense oligonucleotide [ASO; sequence: r(CCCUAA)] targeting (TERRA)_10_, introduced either during sample preparation or after the formation of multicomponent condensates (**Fig. 6a**). ASO-treated condensates did not show any signs of RNA clusters even after 96 hours in our fluorescence microscopy experiments (**Fig. 6b, c; Supplementary Fig. 28, 29**). VPT-based nanorheology confirmed the absence of nanoscale clusters (**Fig. 6d; Supplementary Fig. 30**) as no substantial change in condensate viscosity was observed during 24 hours of aging (**Fig. 5f**). The ability of the ASO to prevent RNA clustering is specific to its sequence design as a scrambled ASO variant [sequence: r(CACUAC)] failed to prevent RNA clustering (**Supplementary Fig. 31**). Addition of the ASO after the formation of intra-condensate RNA clusters resulted in the disassembly of RNA clusters, with the rate of disassembly correlating inversely with the sample age possibly linked to the growing dynamical arrest of the clustered RNA-enriched phase (**Fig. 6e**; **Supplementary Fig. 32; Supplementary Videos 20, 21; Fig. 5c**). These results suggest that selectively targeting homotypic RNA-RNA interactions can buffer against intra-condensate RNA aggregation. Could such buffering effects also be imparted by multivalent RBPs with broad specificity for a diverse repertoire of RNA? We tested this idea utilizing G3BP1, which is a core scaffolding RBP of cytoplasmic stress granules^81, 82, 83^. G3BP1 binds to RNAs through a folded RNA recognition motif and disordered Arg/Gly-rich domain (**Fig. 6f**). We found that G3BP1 positively partitions within (TERRA)_10_ containing RGG-d(T)_40_ condensates. G3BP1 concentration was observed to be at least 50-fold higher in the condensate interior as compared to the input concentration (10 μM) (**Supplementary Fig. 33, 34**). In the presence of G3BP1, we observe the emergence of microscale RNA clusters with significant delays. The cluster sizes derived from SAC in the first 24 hours were below the detection limit (0.14 ± 0.01 μm at 15 minutes and 0.14 ± 0.01 μm at 24 hours; the detection limit is 0.2 μm). However, microscale RNA clusters were first observed at 48 hours timepoint (2.52 ± 0.46 μm) (**Fig. 6g, h; Supplementary Fig. 35-37**).

**Figure 6.**
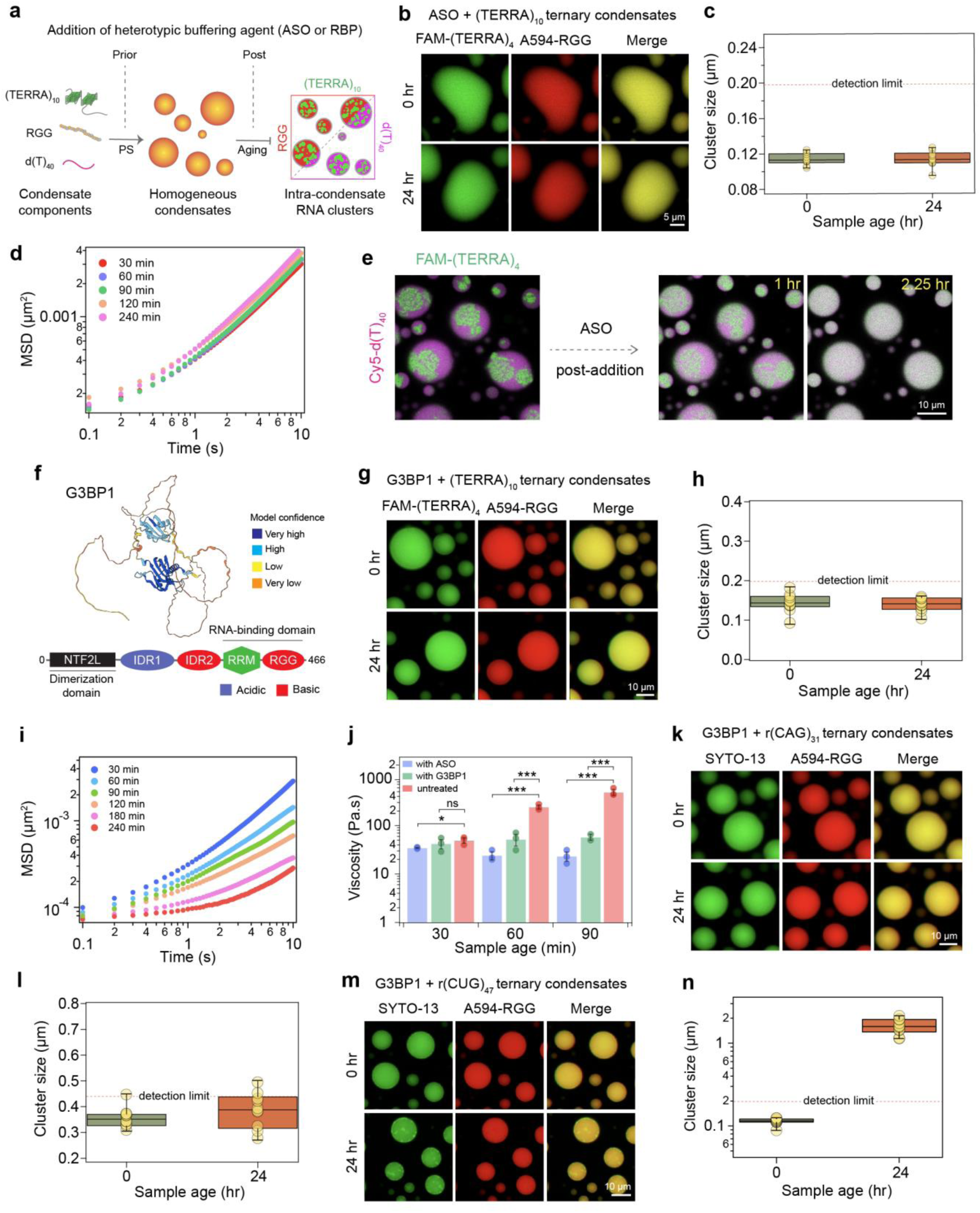
Heterotypic buffering by ASO and G3BP1 can prevent homotypic RNA clustering in biomolecular condensates. **(a)** Schematic depicting the experimental approach to probe whether heterotypic buffering agents such as an ASO or RBP can inhibit intra-condensate RNA clustering. **(b)** (TERRA)_10_ containing RGG-d(T)_40_ condensates treated with TERRA antisense oligonucleotide [ASO; sequence: r(CCCUAA)] does not form age-dependent RNA clusters. **(c)** The corresponding cluster sizes derived from SAC are reported. The detection limit of SAC is demarcated (see Methods for further details). **(d)** Age-dependent MSDs of 200 nm beads inside (TERRA)_10_ containing RGG-d(T)_40_ condensates treated with ASO (also see **Supplementary Fig. 30**). **(e)** Addition of ASO [sequence: r(CCCUAA)] to (TERRA)_10_ containing RGG-d(T)_40_ condensates after the formation of RNA clusters (5 hours since sample preparation). These observations correspond to **Supplementary Video 20**. The condensate composition is similar as in 6b except the concentrations of all condensate components and ASO were halved. Also, see **Supplementary Fig. 32** and **Supplementary Video 21**. **(f)** Alphafold2^85^ predicted structure of G3BP1 (identifier: AF-Q13283-F1) with color coding corresponding to model confidence. Domain architecture of G3BP1 (NTF2L, nuclear transport factor 2-like domain; IDR, intrinsically disordered region; RRM, RNA recognition motif; RGG, Arg/Gly-rich domain). **(g)** (TERRA)_10_ containing RGG-d(T)_40_ condensates with G3BP1 do not show age-dependent morphological changes. **(h)** The corresponding cluster sizes derived from SAC indicate a lack of microscale RNA clustering. The detection limit of SAC is demarcated (see Methods for further details). **(i)** Age-dependent MSDs of 200 nm beads inside (TERRA)_10_ containing RGG-d(T)_40_ condensates with G3BP1 (also see **Supplementary Fig. 38**). **(j)** Comparison of terminal viscosities of (TERRA)_10_ containing RGG-d(T)_40_ condensates with or without the ASO or G3BP1 [“untreated” data taken from Fig. 5h]. Statistical significance was determined by performing a two-sided Student’s *t*-test (* means p<0.05, ** means p<0.01, *** means p<0.001, ns means ‘not significant’) between viscosities of untreated condensates versus condensates containing either the ASO or G3BP1. The p-values, between ‘untreated’ and ‘with ASO’ condition at 30 minutes, 60 minutes, and 90 minutes sample age, are 0.041, 0.0003, and 0.0009, respectively. The p-values between ‘untreated’ and ‘with G3BP1’ conditions at 30 minutes, 60 minutes, and 90 minutes sample age are 0.4766, 0.0009, and 0.0005, respectively. Error bars denote the standard deviation. **(k)** r(CAG)_31_ containing RGG-d(T)_40_ condensates with G3BP1 do not show age-dependent morphological changes. **(l)** The corresponding cluster size analysis indicates an absence of RNA clustering. The detection limit of SAC is demarcated (see Methods for further details). **(m)** r(CUG)_47_ containing RGG-d(T)_40_ condensates with G3BP1 do not show visible RNA clusters at 15 minutes after sample preparation but show some RNA clusters at an age of 24 hours. **(n)** The corresponding cluster size analysis is reported. The detection limit of SAC is demarcated (see Methods for further details). All box plot elements are defined similarly to Fig. 2d. The concentrations of the ASO and G3BP1 are 1 mg/ml and 10 μM, respectively. The composition of the condensate system used for imaging is 1 mg/ml RNA [0.45 mg/ml in the case of r(CUG)_47_; (TERRA)_10_, 101 μΜ, r(CAG)_31_, 33 μM; r(CUG)_47_, 10 μM], 5 mg/ml RGG, and 1.5 mg/ml d(T)_40_. For the nanorheology measurements, the relative proportion of the condensate components was kept the same, but the overall concentration of each component was doubled to achieve a higher volume fraction of the dense phase. Buffer composition for all experiments is 25 mM Tris-HCl (pH 7.5), 25 mM NaCl, and 20 mM DTT. In experiments utilizing fluorescently labeled components, the concentration range is 250 nM to 500 nM. Each experiment was independently repeated at least three times, except for the sample with G3BP1 in (h) that aged for 90 minutes, which was independently repeated two times.

Earlier, our nano-rheology experiments on (TERRA)_10_ containing RGG-d(T)_40_ condensates (**Fig. 5g, h**) revealed signs of dynamical arrest and a substantial increase in viscosity even before microscale RNA clusters were visible. To quantify the effects of G3BP1 on the rheology of these condensates, we probed the time-dependent changes in condensate material properties using VPT. MSD profiles and viscosity measurements reveal that despite the absence of microscale RNA clusters, G3BP1 containing condensates undergo a progressive dynamical slowdown (**Fig. 6i; Supplementary Fig. 38**). There is a concomitant increase in viscosity from 42.3 ± 9.0 Pa.s for condensates at 30 minutes to 154 ± 22.3 Pa.s within 3 hours of condensate preparation (**Supplementary Fig. 38**). However, condensate aging is substantially slower in the presence of G3BP1 (**Fig. 6j**). Therefore, although microscale RNA clusters are disfavored in the presence of G3BP1, our rheology measurements suggest that there may be nanoscale RNA clustering that contributes to increased viscoelasticity of aged condensates. Conversely, G3BP1 can increase the activation energy barrier of intra-condensate (TERRA)_10_ clustering. However, the addition of G3BP1 to pre-formed intra-condensate RNA clusters did not lead to their disassembly (**Supplementary Fig. 39**). RNA concentration measurements in condensates show a higher partitioning of RNA in G3BP1-treated condensates but not in ASO-treated condensates, relative to untreated conditions (**Supplementary Fig. 40**). Interestingly, a substantial difference in ThT fluorescence is observed, with ASO-treated condensates showing a lack of ThT fluorescence and G3BP1 condensates showing prominent ThT fluorescence similar to untreated ternary condensates (**Supplementary Fig. 41**; see also **Fig. 2h**).

We further tested the generalizability of the observed buffering effect of G3BP1 with GC-rich repeat RNAs, r(CAG)_31_ and r(CUG)_47_, that spontaneously form microscale clusters in RGG-d(T)_40_ condensates (**Fig. 4**). In the presence of G3BP1, both RNAs formed homogeneous condensates (**Fig. 6k-n; Supplementary Fig. 36**). Furthermore, in the case of (CAG)_31_, no microscale clusters were observed even at 24 hours age where the cluster size = 0.39 ± 0.02 μm, close to the detection limit (440 nm) (**Fig. 6k, l; Supplementary Fig. 36**). In the case of r(CUG)_47_ we observed the formation of a few RNA foci at 24 hours (**Fig. 6m, n; Supplementary Fig. 36**). However, the total fraction of RNA, determined by intensity-based analysis, in these clusters were only ∼6.65 ± 2.31% (**Supplementary Fig. 42**), whereas the same RNA was present almost exclusively (∼100%) in the intra-condensate clusters in the absence of G3BP1 (**Fig. 4d**). Together, these results signify that G3BP1 can frustrate^84^ RNA-RNA homotypic interactions in these condensates thereby disfavoring RNA clustering and preserving intra-condensate solubility of repeats RNAs.

## Conclusions

Recently, it has been shown that biomolecular condensates can act as sites for pathological protein aggregation^21, 22, 23, 24, 25, 86^. We now demonstrate that irreversible clustering of repeat expanded RNA molecules, a process widely implicated in many neurological disorders^37, 64, 65, 66, 67, 68, 69^, can be nucleated in multi-component protein-nucleic acid condensates. The underlying mechanism is sequence-specific percolation transitions of RNA chains driven by homotypic RNA-RNA interactions. RNA percolation engenders an age-dependent liquid-to-solid phase transition of condensates. Multivalent co-factors, such as ASO and RBPs, that compete with homotypic RNA-RNA interactions can increase the activation energy barrier of RNA clustering, thereby acting as inhibitors of this process.

Two important implications, as outlined in **Fig. 7**, stem from our experiments reported in this study. Firstly, dynamic frustration of homotypic interactions between RNA chains can increase the range of metastability of biomolecular condensates that are poised to undergo RNA percolation-driven physical aging. In this model, a multi-component biomolecular system is frustrated if the probability of minimizing its global free energy through coordinated optimization of all possible interaction modalities of constituent macromolecules is kinetically sluggish due to overlapping inter-molecular interactions^84, 87, 88, 89, 90^. For RNA containing RGG-d(T)_40_ condensates, the RNA [e.g., (TERRA)_10_ or r(CAG)_31_ or r(CUG)_47_] is initially kept frustrated through heterotypic interactions with the primary condensate components, despite having a strong propensity to self-associate as evidenced by its hysteretic phase behavior and irreversible clustering^43^ (**Fig. 2**). However, with time, RNA-RNA homotypic interactions dominate, leading to age-dependent RNA clusters that demix from the fluid phase formed by RGG and d(T)_40_. The onset of RNA clustering is determined by the driving force for RNA percolation, which is modulated by repeat RNA length and/or sequence perturbations that weaken base pairing and stacking interactions. The condensate microenvironment as well as binding to the RGG polypeptide, could play an active role in this process by potentially lowering the energy barrier for unfolding or expansion of folded RNA chains stabilized by intra-molecular interactions^91, 92, 93, 94, 95^, thereby shifting the dynamical equilibrium from dominant intra-molecular interactions to inter-molecular RNA-RNA interactions. Further, the observed differences in the timescale of clustering between (TERRA)_10_ versus r(CAG)_31_ or r(CUG)_47_ might stem from the distinction in base pairing and the base-stacking propensity of these RNAs. The formation of a G-quadruplex structure for (TERRA)_10_ as opposed to a hairpin structure in the triplet repeat RNAs^96^ could impact the timescale of RNA clustering due to a combination of factors, including the kinetic barrier of unfolding, number of available sites and modes for inter-molecular RNA-RNA interactions, and the conformational entropy of the RNA chains^40, 97, 98^. Reducing the homotypic RNA-RNA interactions, and hence, the percolation propensity shifts the balance towards heterotypic interactions between RNA and condensate components over RNA self-assembly. Consequently, the cross-play of interactions between all components, as opposed to the dominance of a singular intermolecular, homotypic interaction node, may be essential for enhancing the metastability of multi-component biomolecular condensates.

**Figure 7.**
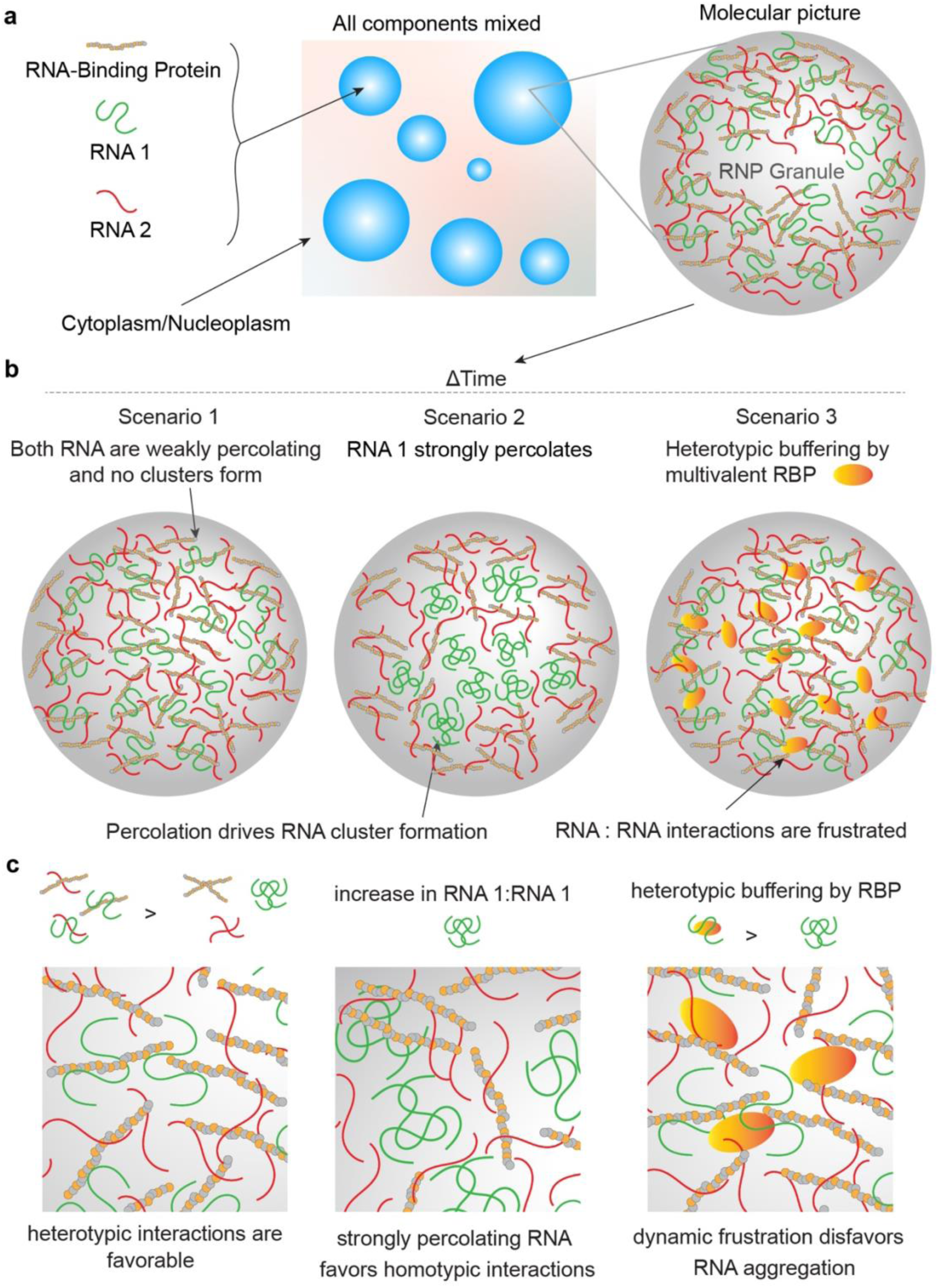
A schematic showing the proposed model of intra-condensate RNA percolation and heterotypic buffering. **(a)** A model of multi-component condensates formed by two RNAs with strong (as shown in green) and weak (as shown in red) percolation propensity, respectively, and an RBP. **(b)** Three possible scenarios of RNA percolation-driven condensate aging or a lack thereof in the presence of a multivalent RBP. **(c)** Zoomed-in views of the panels shown above.

Second, in the cases of strongly percolating RNA molecules [e.g., (TERRA)_10_ or r(CAG)_31_ or r(CUG)_47_], G3BP1 reduces the propensity of RNA clustering suggesting that the solubility of RNAs in condensates is enhanced. Based on these observations, we propose that RNA binding proteins in complex biomolecular condensates may employ heterotypic RNA-protein interactions^99^ as a regulatory mechanism to prevent RNAs from aberrant homotypic self-assembly. This phenomenon, termed heterotypic buffering^99^, was previously proposed as a mechanism to enhance the solubility of aggregation-prone proteins, which possess a strong preference for homotypic interactions and amyloid fiber formation linked to diseases. Our study extends this thermodynamic framework to rationalize the biological roles of RBPs in regulating intra-condensate RNA aggregation. During cellular stress, polysomes are disassembled and polyA-tailed mRNAs are sequestered in stress granules, thereby globally inhibiting translation^100, 101^. Intra-stress granule RNA percolation can compromise the disassembly of stress granules upon removal of stress, leading to the accumulation of irreversible granules that are cytotoxic^22, 102^. Our results indicate that multivalent RNA binding proteins can effectively provide the first line of defense against irreversible RNA clustering before ATP-dependent RNA helicases can actively engage in remodeling these assemblies^78^.

In summary, we report that percolation-driven irreversible RNA clustering can be enhanced in biomolecular condensates, leading to their liquid-to-solid phase transitions. This can be buffered by multivalent RBPs supporting liquid-phase condensate homeostasis. The insights gained from our study provide a complementary perspective on the role of RBPs in regulating aberrant RNA self-assembly in living cells.

## Data availability

All data are available in the manuscript or the supplementary materials.

## Code availability

Codes for nanorheology, SAC, RNA state diagram, co-localization analyses, condensate fusion assay, and complex shear moduli estimation are available on GitHub (see https://github.com/BanerjeeLab-repertoire/Biomolecular-Condensates-Can-Enhance-Pathological-RNA-Clustering).

## Acknowledgments

This work was supported by the US National Institutes of Health through grants R35 GM138186 (P.R.B) and the St. Jude Children’s Research Collaborative on the Biology and Biophysics of RNP Granules (P.R.B.). We gratefully acknowledge Dr. Paul Taylor’s lab at St. Jude Children’s Research Hospital for providing purified G3BP1 protein. We also gratefully acknowledge Dr. Ibraheem Alshareedah, currently at Boston Children’s Hospital, Harvard Medical School for providing the initial SAC analysis codes. The authors deeply appreciate critical feedback from Drs. Rohit Pappu and Tanja Mittag and the group members of Banerjee laboratory.

## Author contributions

Conceptualization: P.R.B., and T.M.; Methodology: P.R.B., T.M., A.S., and G.M.W.; Investigation: P.R.B., T.M., A.S., and G.M.W.; Resources: P.R.B.; Writing – original draft and revisions: P.R.B., G.M.W., and T.M.; Writing – reviewing and editing: all authors except R.G.; Funding acquisition: P.R.B. R.G. only participated at the development phase of this study.

