## Supplementary Information for "Biomolecular Condensates Can Enhance Homotypic RNA Clustering"

### Materials and Methods

#### RNA reconstitution and sample preparation for temperature-controlled microscopy

The RNAs used in the present study were purchased from Integrated DNA Technologies (IDT; refer to **Supplementary Tables 1** and **2** for RNA sequences used in this study). The RNAs were reconstituted in RNase-free water followed by centrifugation at 23000xg for 2 minutes to remove any solid particles. The supernatant was extracted with a final concentration of at least 5 mg/ml, which was subsequently stored at −20°C. The resulting stock solutions were divided into multiple aliquots before storage. We also confirmed the absence of microscale aggregates in all RNA stock solutions through light microscopy. Preparation of RNA samples for temperature-controlled microscopy was performed following our previous work[1] and is briefly outlined below. Glass slides made of borosilicate (25.4 mm × 76.2 mm) were resized to approximately 20 mm × 25.4 mm using a carbide blade glass scribe from Thorlabs. The slides and coverslips (18 mm × 18 mm × 0.17 mm) used in the experiments were coated with Tween-20 [20% (v/v) solution in MilliQ water] for 30 minutes. Subsequently, they were rinsed with MilliQ water, followed by drying with compressed air and heating at 40 °C for 16 hours. Next, we created sample chambers with a sample volume of ~5 µl using double-sided tape. RNA samples with desired concentrations were prepared by diluting the stock solutions in a buffer containing 50 mM HEPES–KOH (pH 7.5) and variable $MgCl_2$, as indicated in the text and figures. After filling the channels with RNA solution, mineral oil was applied to seal both sides of the channel to prevent evaporation.

#### Temperature controlled microscopy

The RNA samples were imaged using a Zeiss Primovert microscope equipped with a 40× air objective immediately after preparation. The microscope was equipped with a Blackfly S USB3 CMOS camera (Teledyne FLIR) and a temperature-controlled stage (Instec) with an accessible temperature range of 2 to 90°C. The temperature-controlled microscopy instrumentation and measurements of RNA samples were conducted following a step-by-step protocol that we previously reported in detail[1]. Recording of the temperature at which RNA condensation occurred upon heating ($T_{phase}$) during data acquisition was done manually. The reported $T_{phase}$ values were determined through a minimum of three technical replicates. RNA phase diagrams were generated using the Matplotlib library (https://matplotlib.org/) in Python (https://www.python.org/).

#### Polypeptide and single-stranded nucleic acid stock preparation

RGG peptide (sequence: $[RGRGG]_5C$) was synthesized by GenScript USA Inc. (NJ, USA, >90% purity). A cysteine residue was incorporated at the C-terminal of the peptide for site-specific fluorescence labeling using cysteine-maleimide chemistry[2]. The peptide was reconstituted in RNase-free water (Santa Cruz Biotechnology) supplemented with 50 mM dithiothreitol (DTT) (ThermoFisher Scientific) to prevent cysteine oxidation during storage.

Homopolymeric single-stranded DNA, $d(T)_{40}$, and homopolymeric single-stranded RNA, $r(U)_{40}$, were purchased from Integrated DNA Technologies (IDT). Polyuridylic acid, poly(rU), was purchased from Sigma-Aldrich (cat # P9528). The nucleic acids were reconstituted in RNase-free water. The reconstituted solutions were centrifuged at 23000xg for 2 minutes to remove any solid particles and the supernatant was removed. Nucleic acid concentration was subsequently measured using a NanoDrop 1C™ spectrophotometer. The resulting stock solutions of both polypeptide and nucleic acid were divided into multiple aliquots and stored at −20°C. We further used microscopic examination to confirm the absence of aggregates in all polypeptide/nucleic acid stock solutions.

#### Multicomponent condensate preparation and imaging

Multicomponent peptide-nucleic acid condensates were prepared by mixing $[RGRGG]_5$ (hereafter, RGG) polypeptide, $d(T)_{40}$ ssDNA, and RNA in a buffer containing 25 mM Tris-HCl (pH 7.5), 25 mM NaCl, and 20 mM DTT unless otherwise noted. The polypeptide was added as a final component to the mixture of ssDNA and RNA to induce phase separation. For fluorescence imaging of condensates, trace amounts (250 nM to 500 nM) of Cy5-labeled $d(T)_{40}$ [IDT; Cy5- $d(T)_{40}$], SYTO-13 (Invitrogen), which is a nucleic acid-specific fluorophore, FAM-labeled RNA, and Alexa594-labeled RGG (A594-RGG) were used as indicated in appropriate sections of the text and figures. Polypeptide labeling with Alexa594 (Invitrogen) was achieved through cysteine-maleimide chemistry following the manufacturer-provided instructions. Unless stated otherwise, the total polypeptide concentration is 5.0 mg/ml, RNA concentration is 1 mg/ml, and the $d(T)_{40}$ concentration is 1.5 mg/ml in all condensate samples, corresponding to a polypeptide-polynucleotide mass ratio of 0.5. This mass ratio of polypeptide to nucleic acid was previously shown to be optimal for phase separation[3]. In the case of nanorheology measurements, the same mass ratio was used but the overall concentrations were doubled to achieve larger condensates for video particle tracking measurements with higher throughput. Across experiments reported in this study, the identity of the RNA in the system was varied but the RNA concentrations remained constant (unless specified otherwise), and all buffer conditions were kept identical for comparison between experiments (unless specified otherwise).

The preparation of the quaternary condensate systems with G3BP1 followed a similar procedure with the following order of addition: 10 µM G3BP1 was added along with other components [$d(T)_{40}$ and RNA] before adding RGG peptide. A G3BP1 stock of 170 µM in 150 mM HEPES (pH 7.5) and 150 mM KCl was diluted to 10 µM during sample preparation. As a negative control, we used a 3-component condensate sample containing RGG, $d(T)_{40}$, and RNA along with a 17-folded diluted G3BP1 buffer [50 mM HEPES (pH 7.5) and 150 mM NaCl without G3BP1 protein]. This control sample underwent RNA clustering at a similar timescale in comparison to a condensate sample only containing the native buffer, 25 mM Tris-HCl (pH 7.5), 25 mM NaCl, and 20 mM DTT, ensuring that G3BP1-induced effects did not arise due to mild variations in sample buffer conditions (**Supplementary Fig. 36**). Fluorescence imaging of multicomponent condensate samples was done either using a Zeiss LSM710 laser scanning confocal microscope (Plan-Apochromat 63×/1.4 oil DIC M27) or a Q2 laser scanning confocal microscope (ISS Inc., 63× objective). Super-resolution imaging of intra-condensate RNA clusters was performed using a Leica TCS SP8 confocal microscope equipped with the Leica Lightning super-resolution acquisition and deconvolution package. 3D rendering of the acquired super-resolution images was done using Fiji[4] (version 1.54f).

### Estimation of RNA and protein concentration in condensates

For quantification of RNA concentration in $(TERRA)_4$ containing RGG-$d(T)_{40}$ condensates with or without G3BP1 or ASO, FAM-$(TERRA)_4$ fluorescence intensity as a function of concentration was used as a readout, similar to that reported in a previous study[5]. Briefly, fluorescence readings of FAM-$(TERRA)_4$ prepared in a buffer composed of 25 mM Tris-HCl (pH 7.5), 25 mM NaCl, and 20 mM DTT, were collected using the Q2 laser scanning confocal microscope (ISS Inc., 63× objective) equipped with single-photon avalanche diodes (SPADs). These measurements yielded a concentration versus fluorescence intensity calibration curve for FAM-$(TERRA)_4$. Next, FAM-$(TERRA)_4$ fluorescence readings from ternary RGG-$(TERRA)_4$-$d(T)_{40}$ condensates were interpolated to estimate the intra-condensate concentration of FAM-labeled $(TERRA)_4$. Subsequently, the interpolated measurements were transformed by a factor that accounts for the ratio of the bulk total RNA concentration to the bulk labeled RNA concentration used in our experiments, yielding the intra-condensate total RNA concentration. All measurements were taken while maintaining consistent image acquisition and processing settings. Estimating the concentration of G3BP1 in a quaternary condensate system composed of $(TERRA)_4$, RGG, $d(T)_{40}$, and G3BP1 involves the use of a similar strategy except the use of Alexa488-labeled G3BP1. The data used for concentration estimations were gathered from 30 condensates across 3 independently prepared samples.

### Fluorescence recovery after photobleaching (FRAP)

Condensate samples consisting of 5 mg/ml RGG, 1.5 mg/ml $d(T)_{40}$, and 1 mg/ml $(TERRA)_{10}$ (as described in the '*Multicomponent condensate preparation and imaging*' section above) in a Tween-20 coated imaging chamber was placed in a confocal microscope stage (Lumicks C-trap) and imaged with a 63× oil objective. Imaging for reference and bleaching steps of FAM-$(TERRA)_4$, A594-RGG, and Cy5-$d(T)_{40}$ were accomplished using the respective excitation laser lines. The regions of interest (ROIs) for reference, bleaching, and continuous imaging were selected using the Bluelake software (Lumicks, https://lumicks.github.io/bluelake-api/2.4.0/index.html). Samples were bleached for 10-20 frames (selected based on the optimal bleach depth) with a dwell time of 150 ms per pixel. Recovery was observed for 92 s after bleaching. The FRAP images were analyzed using Fiji[4] (version 1.54f) to quantify the fluorescence recovery profiles. The intensities were normalized and corrected as described earlier[6]. In brief, the normalization takes into account the correction for background intensity after bleaching and subsequent photofading using the following expression, where $I_{correct}(t)$ is the intensity of the bleached ROI at time $t$ corrected for photofading and $I_{correct}(0)$ is the value of intensity before bleaching starts (reference ROI to quantify the bleaching depth):

$$I_{normalized}(t) = \frac{I_{correct}(t) - \min(I_{correct})}{I_{correct}(0) - \min(I_{correct})} \qquad (1)$$

At least 6 recovery profiles from three independent experiments were collected and averaged. Intensity error bars were estimated using the standard error at each time point recorded.

#### Thioflavin T (ThT) assay

To probe the presence or absence of G-quadruplex structures within multicomponent condensates containing RNAs, 50 µM ThT was used[7, 8, 9]. ThT was premixed with the experimental buffer before adding condensate-forming components. Simultaneous fluorescence imaging of the condensates was performed by using ThT (excitation with a 488 nm laser) and Cy5-d(T)$_{40}$ signals. Fluorescence imaging was done using a Q2 laser scanning confocal microscope (ISS Inc., 63× objective).

#### 2D spatial autocorrelation (SAC) analysis of confocal images

SAC analysis was performed on the confocal fluorescence images to estimate the size of the RNA clusters (or a lack thereof) formed inside condensates. This size of the RNA clusters was captured by fluorescence imaging at multiple time points. Random regions close to the middle of condensates were chosen in a fluorescence image and used as input to the analysis. The selected regions were converted into a color-mapped image. Fourier transform was performed using the Weiner-Khinchin theorem[10] to obtain the 2D spatial autocorrelation functions. The spatial autocorrelation profiles in the X-axis were further approximated with a Gaussian fit to estimate the cluster sizes (see **Supplementary Fig. 14**). Image data used for SAC was gathered from 10 condensates across 3 independently prepared samples. We verified the size estimation capacity of SAC analysis using fluorescent probe particles of known sizes. The probe particles used are 1 µm, 200 nm, and 100 nm yellow-green carboxylate-modified polystyrene beads (FluoSpheres™, Invitrogen) reconstituted at a low concentration (0.0005 % solids) in a sample containing condensates in order to mimic the solution conditions of SAC analysis performed for RNA cluster size evolution [sample composition: 5 mg/ml RGG and 5 mg/ml d(T)$_{40}$ in buffer containing 25 mM Tris-HCl (pH 7.5), 25 mM NaCl, 20 mM DTT]. We find that SAC is capable of capturing relative size differences reliably when the particle sizes are above the resolution (r) of the confocal microscope (r = 198 nm for ISS Q2 laser scanning confocal microscope), whereas, for particles below this limit, SAC provides the size of spatial fluctuations converging to the diffraction limit i.e. image resolution (see **Supplementary Fig. 15**). Hence, the detection limit of SAC is based on the resolution of the images acquired by confocal fluorescence microscopy. The pixel size of the microscope is set by the user to match the spatial frequency determined by the Nyquist bandwidth of the microscope as closely as possible. This corresponds to 46 nm in xy and 119 nm in z, which can be calculated using the formulas for a line-scanning confocal microscope[11], where '$Pixel_{xy}$' is the xy pixel size, '$Pixel_z$' is the z pixel size, '$\lambda_{ex}$' is excitation wavelength, '$n$' is the refractive index, '$\theta$' is the half-angle of the objective's aperture:

$$Pixel_{xy} = \frac{\lambda_{ex}}{8n \sin\theta} \qquad (2)$$

$$Pixel_z = \frac{\lambda_{ex}}{4n\,(1 - \cos\theta)} \qquad (3)$$

To delineate the detection limit of SAC for the image datasets acquired using the ISS Q2 laser scanning confocal microscope, we empirically measure the microscope's point spread function (PSF) using 100 nm beads, which is expected to be above the theoretical resolution following the Rayleigh criterion[12], where '$r$' is the resolution, ' $\lambda_{ex}$' is the excitation wavelength, and '$NA$' is the numerical aperture of the objective:

$$r = \frac{0.51\,\lambda_{ex}}{NA} \qquad (4)$$

We demonstrate that SAC converges to the theoretical limit of r = 191.45 nm, i.e., 100 nm beads yield 197.5 ± 11.9 nm (**Supplementary Fig. 15a**). Thus, we set the detection limit to the experimentally determined value (198 nm).

To delineate the detection limit of SAC for the image datasets acquired using the Zeiss LSM710 laser scanning confocal microscope, we refer to the Nyquist sampling theorem, which states that the sampling frequency must be at least two times the object resolution to achieve optimal sampling, and we, therefore, choose two times the pixel size to be the detection limit (440 nm).

### Video Particle Tracking (VPT) nanorheology

The condensate samples for VPT measurements were prepared using yellow-green carboxylate-modified polystyrene beads (200 nm in diameter, FluoSpheres™, Invitrogen) in the sample buffer at a low concentration (0.0005 % solids). Fluorescence microscopy was performed using a Zeiss Primovert inverted microscope equipped with a 100× oil immersion objective and a Blackfly S USB3 CMOS camera (Teledyne FLIR). Movies capturing the diffusion of beads within the condensate were recorded with a frame time of 100 ms over a duration of approximately 2 to 5 minutes capturing a total of 1000-1500 frames. Movies with polystyrene beads diffusing inside the condensates were processed using Fiji[4] (version 1.54f). Particles were tracked using the Trackmate[13] software plugin in Fiji. Appropriate intensity filters were used during tracking to exclude the particle aggregates from being tracked.

The detailed protocol for estimating the mean square displacements (MSDs) from the particle trajectories is described in detail in our previous work[3]. To summarize, the extracted trajectories were corrected for drifting by subtracting the center of mass trajectory, which was calculated using the velocities of particles in the following way:

$$\boldsymbol{X_{COM}}(k) = \boldsymbol{X_0} + \sum_{j=0}^{k} \frac{1}{N_j} \sum_{i=1}^{N} \boldsymbol{v_{i,j}} \qquad (5)$$

Where $k$ is the frame number at which $\boldsymbol{X_{COM}}$ is calculated, $j$ is the individual frame, and $N_j$ is the number of particles in frame $j$. $\boldsymbol{X_0}$ is the center of the mass vector of the first frame. After correcting the trajectories for drifting, the mean squared displacement (MSD) was calculated as

$$MSD(\tau) = \langle \boldsymbol{R}(t+\tau) - \boldsymbol{R}(t) \rangle_{t,N} \qquad (6)$$

Where $\tau$ is the lag time and $\boldsymbol{R}$ is the position vector of the particle. The MSD was computed from the trajectories across various lag times $\tau$ using custom Python scripts, which are publicly available on GitHub (https://github.com/BanerjeeLab-repertoire/Biomolecular-Condensates-Can-Enhance-Pathological-RNA-Clustering). The ensemble average MSD was then determined based on data from approximately 20–100 individual beads. The MSD was then fitted[3, 14] using:

$$MSD(\tau) = 4D\tau^{\alpha} + N \qquad (7)$$

to obtain the diffusion coefficient $D$ of the beads. For all the systems, we ensured that the value of the diffusivity exponent $\alpha$ is equal to 1, which signifies terminal viscous behavior, by choosing an appropriately long frame time. The diffusion coefficient is then converted to viscosity using the Stokes-Einstein equation[3, 14]:

$$\eta = \frac{k_B T}{6\pi D R} \quad (8)$$

where $\kappa_B$ is the Boltzmann constant, $R$ is the radius of the particles, and $T$ is the temperature of the sample.

Two-sided Student's *t*-tests were performed using the data analysis toolbox of Microsoft 365 Excel to ascertain statistical significance by determining *P* values for VPT nanorheology-based viscosity measurements.

For each sample, 20-200 trajectories were gathered from 3 to 5 condensates in 3 independently prepared samples, unless stated otherwise. To capture condensate aging dynamics, VPT was performed on the same condensates at different time points during aging.

### Estimation of complex shear moduli from the ensemble-averaged MSD measurements

The mean square displacement of the probe particles embedded inside condensates, measured using VPT nanorheology, can be analyzed using custom Python scripts to estimate the condensate viscoelastic properties. A detailed description of the procedure to estimate the dynamical moduli from the measured MSDs can be found elsewhere[15]. Briefly, the compliance $j(t)$ was estimated from the MSDs $< \Delta r^2(\tau) >$ using,

$$j(t) = \frac{3\pi a}{n_d k_B T} < \Delta r^2(\tau) > \quad (9)$$

Where a is the radius of the probe particle, $n_d$=2 is the dimension of the probe particle trajectories from which MSDs were estimated, $k_B$ is the Boltzmann constant, and $T$ is the absolute temperature at which nanorheology experiments were conducted.

The complex shear modulus and the compliance are related through a convolution integral, which allows the estimation of the complex shear modulus using a Fourier transform,

$$\int_0^\tau G(t) j(\tau - t) dt = \tau \quad (10)$$

$$G^*(\omega) = \frac{1}{i\omega \hat{j}(\omega)}. \quad (11)$$

To estimate the complex shear modulus, we used the method proposed by Evans et al[16]. This method uses discrete experimental data points $(t_i, j_i)$ of the calculated compliance from Eq. (9) to estimate the frequency-space shear modulus $G^*(\omega)$ using the following relation.

$$\frac{i\omega}{G^*(\omega)} = i\omega J(0) + \frac{\left(1 - e^{-i\omega t_1}\right)\left(J_1 - J(0)\right)}{t_1} + \frac{e^{-i\omega t_N}}{\eta} + \sum_{k=2}^{N} \left(\frac{J_k - J_{k-1}}{t_k - t_{k-1}}\right)\left(e^{-i\omega t_{k-1}} - e^{-i\omega t_k}\right) \quad (12)$$

For the calculations, we used a cubic spline to oversample the MSD data from our VPT measurements. Other parameters needed in Eq. (12) to compute the shear modulus can be obtained as follows. $j(0)$ can be estimated by extrapolating the experimental data for compliance to $t = 0$ by a linear fit of the first four data points and $\eta$ by a linear fit of the last ten points using the equation $\eta = 1/j(t)$.

### Optical tweezer-based fusion assay

Controlled fusion of condensates was employed to explore changes in the material properties[6, 17] of multicomponent condensates in the presence or absence of RNA clustering. Samples placed in a Tween-20 coated imaging chamber were subjected to controlled fusion using a correlated dual-trap optical tweezer system and a laser scanning confocal fluorescence microscope (Lumicks C-trap). In these experiments, two droplets were trapped using a 1064 nm laser at the minimum power (1-5% overall power) to minimize the perturbation of condensates by the trapping laser. Droplet trapping relied on the refractive index disparity between the condensate dense phase and the dilute phase. Following trapping, one droplet was brought into contact with the other stationary droplet at a constant velocity of 100 nm/s. The trap maintained this velocity until fusion completion, allowing the final droplet to relax to a spherical shape. Data was analyzed using custom Python scripts to estimate the droplet relaxation timescale.

#### Condensate dissolution assay

A freshly prepared condensate sample of 5 µl volume was placed in a chambered slide and subsequently covered with immiscible mineral oil to prevent sample evaporation (**Supplementary Fig. 23a**). The condensates were allowed to age for specified periods as indicated in the text and figure captions. For condensate dissolution, 0.5 µl volume of 5 M NaCl solution was carefully pipetted onto the oil-embedded condensate sample, making a final concentration of NaCl to 455 mM. The sample was subjected to time-lapse fluorescence imaging with 1 frame taken every two seconds. To account for any disturbance in Z-focus created by NaCl addition, the objective focus was manually adjusted as quickly as possible to regain the initial Z-axis position.

#### G3BP1 purification

GST-tagged G3BP1 protein was expressed and purified from *E. coli* BL21 (DE3) cells in Dr. Paul Taylor's lab at St. Jude Children's Research Hospital as reported before[18]. The GST tag was removed via TEV cleavage. The purity of recombinant G3BP1 protein was analyzed by SDS-PAGE and stored at -80 °C. Fluorescence labeling of G3BP1 with Alexa 488 was achieved through amine-ester chemistry following the manufacturer-provided instructions (Invitrogen).

#### In vitro transcription

r(CUG)$_{47}$ was prepared using in vitro transcription (IVT) as described previously[19]. In brief, the plasmid for expression was pBluescript-CTG-47 (a gift from Ron Vale; Addgene plasmid # 99150). This plasmid was digested using EcoRI (Promega cat # R6011) prior to performing IVT. 1.5 µg DNA was mixed with 5U of EcoRI enzyme and the provided buffer from Promega and incubated for 15 minutes in a water bath at 37°C. The reaction was inactivated by heating at 65°C for 20 minutes. The resulting DNA was then used to perform IVT using a HiScribe T7 kit from NEB (cat # E2040S). Reaction buffer, 10 mM NTPs, 2 µg template DNA, 5 mM DTT, and T7 RNA polymerase master mix were combined and mixed thoroughly by pulse-spin in a microcentrifuge. The reaction was incubated overnight at 37°C. Following IVT the sample was purified using an RNA cleanup kit (Zymo cat # R1015). 150 µL of RNA lysis buffer was added to 50 µL of IVT product. 200 µL of ethanol was added to the mix and transferred to a Zymo-Spin IIICG column in a collection tube and spun at 16000xg for 30s. The sample was collected and treated with DNase1 by washing the column with 400 µL of RNA wash buffer and spun at 16000xg for 30s discarding the flow-through. 5 µL of DNase 1 was added to 75 µL of DNA digestion buffer and added to the matrix of the column. The column was incubated at room temperature for 15 minutes. RNA was eluted from the column and the sample was kept at -20°C until use.

#### Estimation of RNA population in intra-condensate clusters

To determine the percentage of RNA in the intra-condensate clusters, intensity-based thresholding [Fiji[4] (version 1.54f)] was used to determine the area of the clusters ($A_c$) and the area of the cluster-devoid regions in the condensate ($A_{nc}$). The corresponding area of the clusters and the homogeneous phase were multiplied by the average grey scale intensities of the regions ($I_c$ and $I_{nc}$) to get the contribution of the clusters ($A_c \times I_c$) and homogeneous phases ($A_{nc} \times I_{nc}$) to the total fluorescence intensity of the condensate. Finally, the percentage of clusters inside the condensates was estimated from the fraction [$A_c \times I_c / (A_c \times I_c + A_{nc} \times I_{nc})$].

### Surface treatment for microscopy

For all microscopy experiments reported in this study, Tween-20 coated coverslips were used to prevent droplets from excessively wetting the glass surface. Coverslips were first incubated with 2% Hellmanex solution for two hours and then rinsed 6 times with MilliQ water. Next, coverslips were immersed in a 20% (v/v) solution of Tween-20 for 30 minutes. Subsequently, the coated coverslips were rinsed 6 times with MilliQ water and dried using compressed air. Finally, the coverslips were further dried in an oven kept at 40 °C overnight and later stored at room temperature for use.

### Software

Fiji[4] (version 1.54f) was used for image processing. Python scripts were used for graphing temperature-controlled microscopy data, co-localization analyses, VPT nanorheology-based MSD measurements, optical tweezer-based fusion assay, complex shear moduli estimation, and SAC cluster size analyses. These are available on GitHub (https://github.com/BanerjeeLab-repertoire/Biomolecular-Condensates-Can-Enhance-Pathological-RNA-Clustering). In addition, GraphPad Prism 10 (v10.4.1) and IGOR Pro (v6.3 beta 1) were used for graphing and/or data analyses. Temperature-controlled microscopy data acquisition was performed using DAQ software (DAQami, available at https://www.measurementsystems.co.uk/measurement-computing/software-mcc/daqamitrade) and Micromanager (https://micro-manager.org/). ZEN (blue, v2.3) was used for image recording using a Zeiss Primovert microscope. Bluelake (v1.6.11) was used for FRAP image recording and optical tweezer-based fusion assay experiments performed using a Lumicks C-Trap microscope. UCSF ChimeraX[20] (v1.7) was used for visualizing protein structures and preparing the corresponding representative images. For acquiring fluorescence images of condensates, Vistavision (v4.2; ISS Q2 laser scanning confocal microscope) and ZEN (SP5 2012 Black) as part of the Zeiss LSM710 laser scanning confocal microscope were used. The data analysis toolbox of Microsoft 365 Excel was used for determining statistical significance. Adobe Illustrator CC (2024) was used for the assembling figures.

### Supplementary Figures

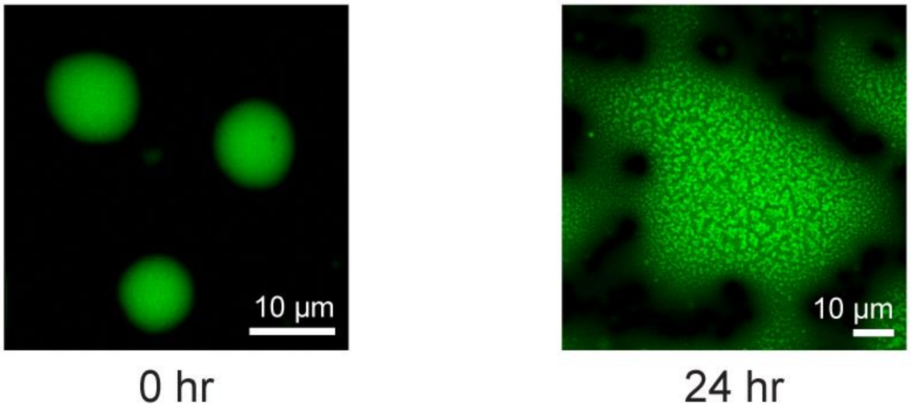


**Supplementary Figure 1.** $(TERRA)_{10}$ partitions homogeneously to RGG-d$(T)_{40}$ condensates at an initial time point (15 minutes after sample preparation; left) but forms intra-condensate clusters in a time-dependent manner (24 hours after sample preparation, right). $(TERRA)_{10}$ was visualized by a FAM-labeled antisense oligonucleotide [sequence: r(CCCUAA)]. The composition of the condensate system used here is 1 mg/ml $(TERRA)_{10}$ (corresponds to 50.7 µM), 5 mg/ml RGG, and 1.5 mg/ml d$(T)_{40}$ in a buffer containing 25 mM Tris-HCl (pH 7.5), 25 mM NaCl, and 20 mM DTT. The concentration of the labeled component is 250 nM. This experiment was independently repeated three times.

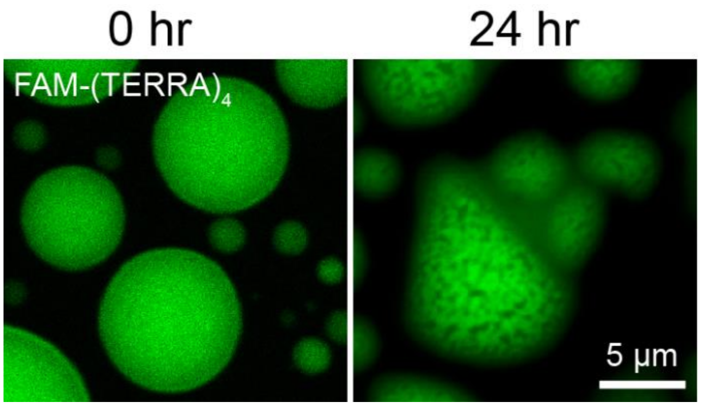


**Supplementary Figure 2.** Formation of RNA clusters in $(TERRA)_{10}$ containing RGG–$r(U)_{40}$ condensates in a time-dependent manner. $(TERRA)_{10}$ was visualized by FAM-labeled $(TERRA)_4$. The composition of the condensate system used here is 1 mg/ml $(TERRA)_{10}$ (corresponds to 50.7 µM), 5 mg/ml RGG, and 1.5 mg/ml $r(U)_{40}$ in a buffer containing 25 mM Tris-HCl (pH 7.5), 25 mM NaCl, and 20 mM DTT. The concentration of the labeled component is 250 nM. This experiment was independently repeated three times.

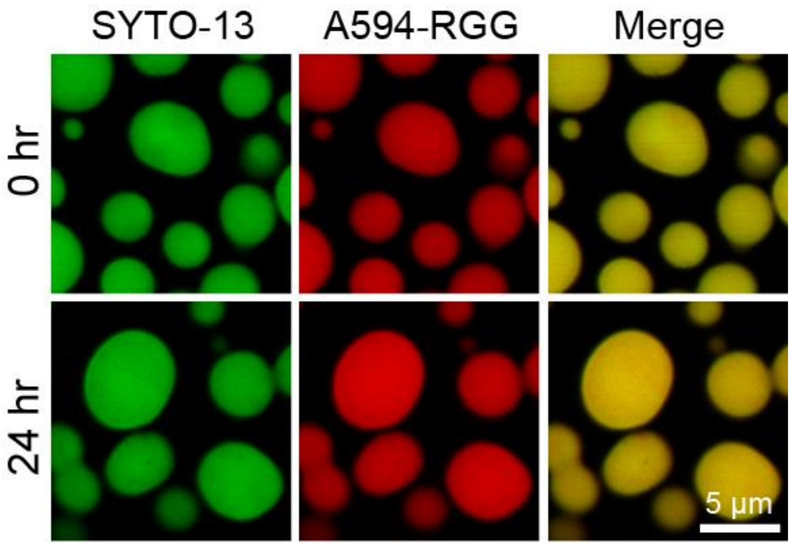


**Supplementary Figure 3.** Fluorescence images of binary condensates consisting of (TERRA)$_{10}$ and RGG. These condensates do not show time-dependent RNA cluster formation. RNA in condensates was visualized by SYTO-13 and the peptide was visualized by Alexa594-labeled RGG. The composition of the condensate system used here is 2.5 mg/ml (TERRA)$_{10}$ (corresponds to 127 µM) and 5 mg/ml RGG in a buffer containing 25 mM Tris-HCl (pH 7.5), 25 mM NaCl, and 20 mM DTT. The concentration range of the labeled components is 250 nM to 500 nM. This experiment was independently repeated three times.

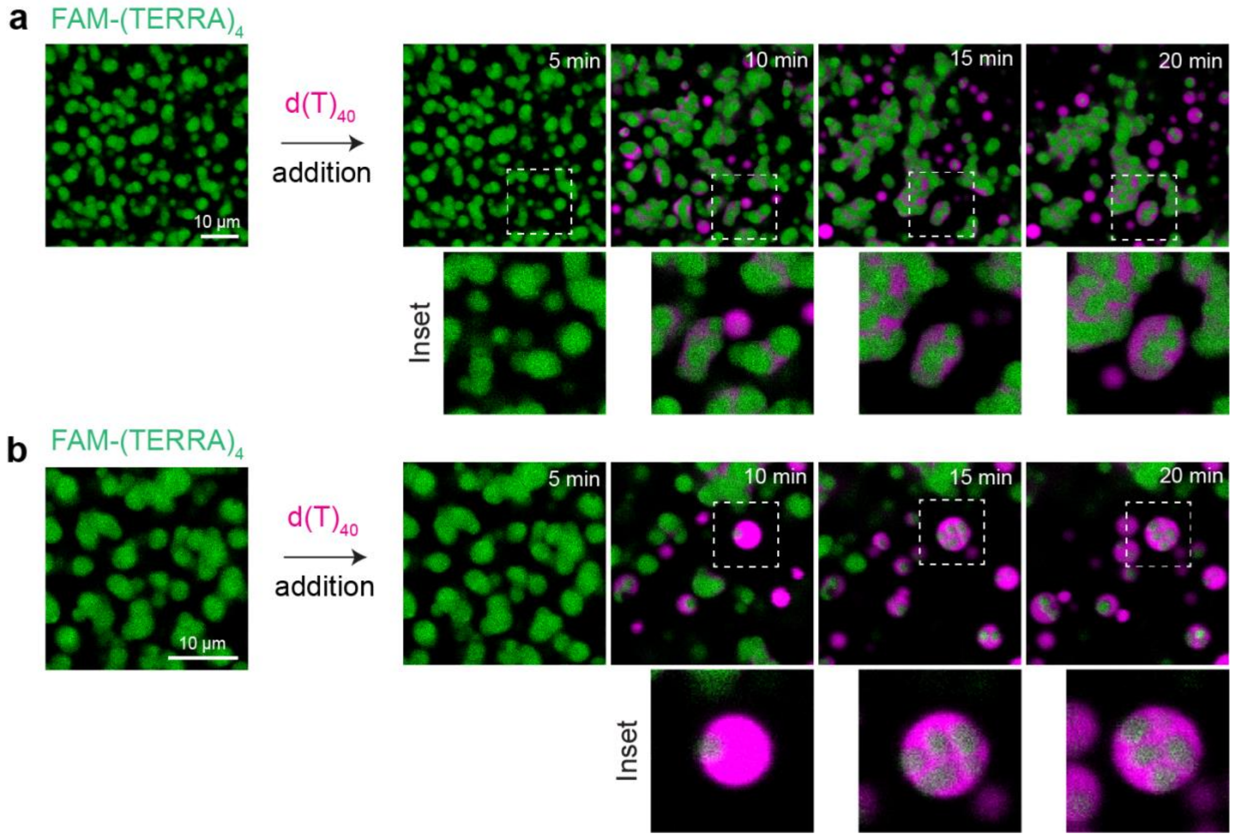


**Supplementary Figure 4.** (a) Lower and (b) higher zoom fluorescence images showing the addition of $d(T)_{40}$, doped with 250 nM Cy5-labeled $d(T)_{40}$ (magenta), to pre-formed binary condensates of $(TERRA)_{10}$ and RGG results in RNA demixing and the formation of RNA clusters visualized using FAM-labeled $(TERRA)_4$ (green). These observations correspond to **Supplementary Video 2.** The composition of the condensate system used here is 0.5 mg/ml $(TERRA)_{10}$ (corresponds to 25.4 µM) and 2.5 mg/ml RGG in a buffer containing 25 mM Tris-HCl (pH 7.5), 25 mM NaCl, and 20 mM DTT. The concentration of $d(T)_{40}$ added is 0.75 mg/ml. The concentration of the labeled components is 250 nM. This experiment was independently repeated three times.

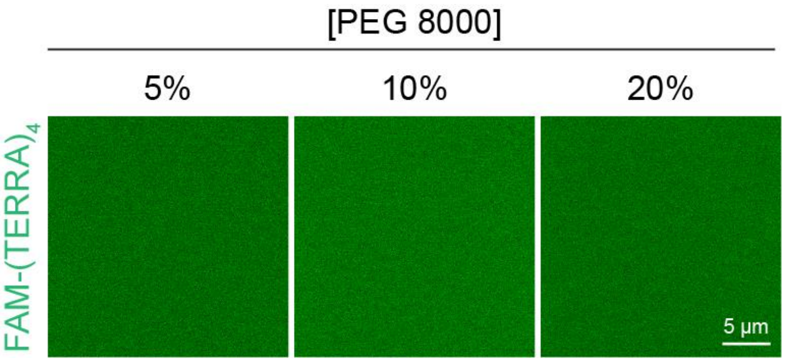


**Supplementary Figure 5.** Effect of titrating a molecular crowder, PEG8000, on $(TERRA)_{10}$ in solution in the absence of condensates. The sample is composed of 1 mg/ml $(TERRA)_{10}$ (corresponds to 50.7 µM) in a buffer containing 25 mM Tris-HCl (pH 7.5) and 25 mM NaCl along with the specified crowder concentration (w/v). The sample is visualized using FAM-labeled $(TERRA)_4$ at a concentration of 250 nM. This experiment was independently repeated three times.

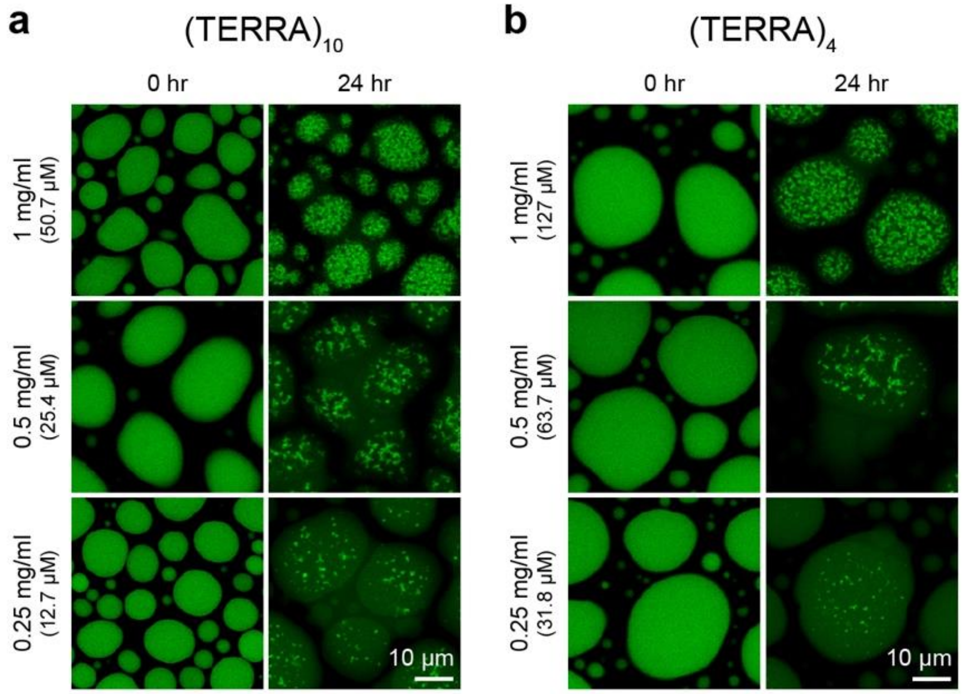


**Supplementary Figure 6.** Effect of titration of bulk RNA concentration in (a) $(TERRA)_{10}$ and (b) $(TERRA)_4$ containing ternary RGG-d$(T)_{40}$ condensate systems on RNA cluster formation upon aging. The condensates and RNA clusters were visualized using FAM-labeled $(TERRA)_4$. The composition of the condensate systems used here is 5 mg/ml RGG, variable d$(T)_{40}$, and variable $(TERRA)_{10}$ concentrations (as indicated in the figure panels), with the total nucleic acid concentration kept constant at 2.5 mg/ml. TERRA concentration (in mg/ml and molarity) used in each sample is indicated in each panel of the figure. The buffer contained 25 mM Tris-HCl (pH 7.5), 25 mM NaCl, and 20 mM DTT. The concentration of FAM labeled $(TERRA)_4$ is 250 nM. Each measurement was independently repeated three times.

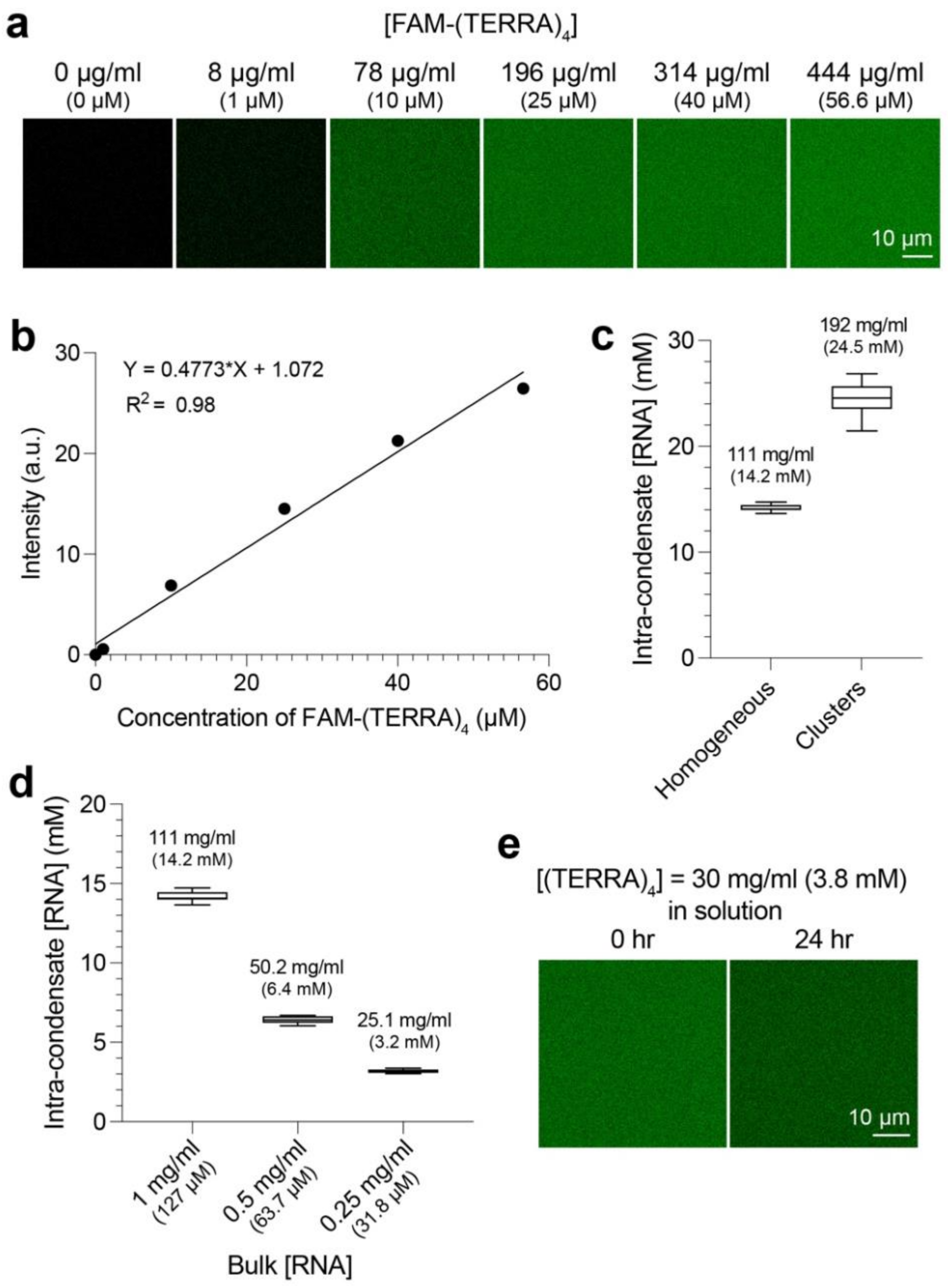


**Supplementary Figure 7.** (a) Fluorescence images of FAM-labeled $(TERRA)_4$ in solution prepared in a buffer containing 25 mM Tris-HCl (pH 7.5), 25 mM NaCl, and 20 mM DTT. (b) Fluorescence intensity versus concentration plot of FAM-labeled $(TERRA)_4$ from images shown in (a). The equation of the fitted line and the $R^2$ value are reported. (c) Estimation of intra-condensate RNA concentration at 15 minutes since sample preparation (homogeneous) and at 24 hours since sample preparation (clusters) based on the calibration curve reported in (b). Additional details of the experiments and analysis are provided 'Materials and Methods' section. The mean intra-condensate RNA concentration is reported in mg/ml and molarity within the plot. The condensate composition is 1 mg/ml $(TERRA)_4$, 5 mg/ml RGG, and 1.5 mg/ml $d(T)_{40}$ in a buffer containing 25 mM Tris-HCl (pH 7.5), 25 mM NaCl, and 20 mM DTT with 180 nM FAM labeled $(TERRA)_4$. (d) Effect of bulk $(TERRA)_4$ concentrations, corresponding to conditions reported in **Supplementary Fig. 6b,** on the intra-condensate RNA concentration. The mean intra-condensate RNA concentration is reported in mg/ml within the plot. (e) 30 mg/ml $(TERRA)_4$ (corresponds to 3.8 mM) in a buffer containing 25 mM Tris-HCl (pH 7.5) and 25 mM NaCl, visualized using 180 nM FAM-labeled $(TERRA)_4$, does not show time-dependent formation of RNA clusters. In (c) and (d), the center line of the box plot represents the median and the whiskers indicate the full range of the data from the minimum to the maximum. Each measurement was independently repeated three times.

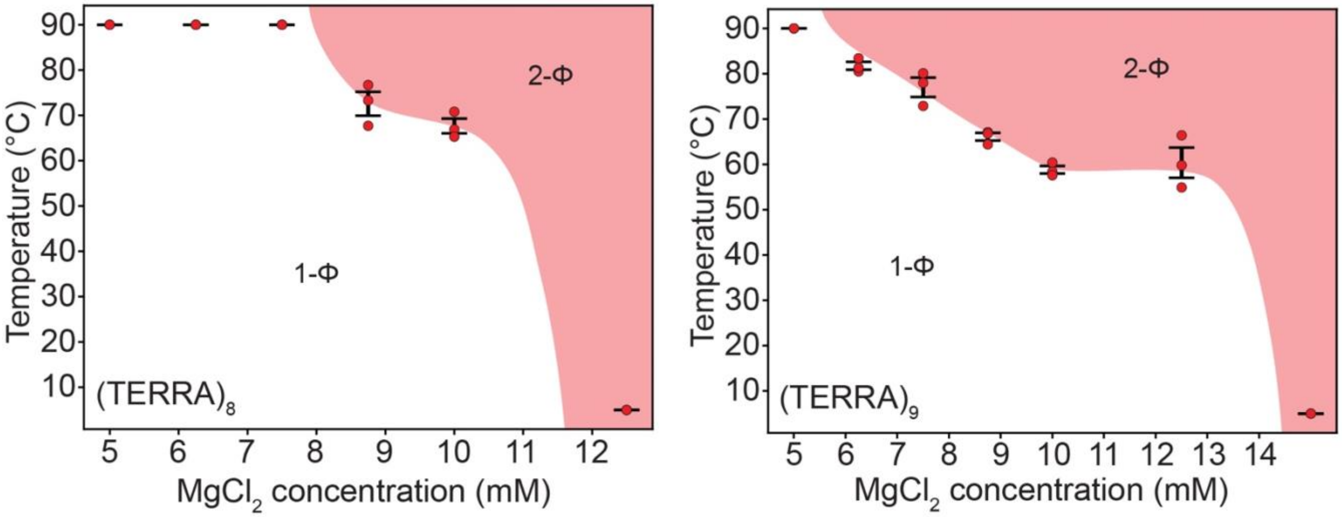


**Supplementary Figure 8.** Temperature-controlled determination of phase diagrams of $(TERRA)_8$ (left) and $(TERRA)_9$ (right) with titrations of $Mg^{2+}$ concentrations. Error bars denote the standard error of the mean (S.E.M.). The concentration of RNA used for temperature-controlled microscopy measurements is 1 mg/ml [$(TERRA)_8$, 63.5 μM; $(TERRA)_9$, 56.4 μM] in 50 mM HEPES (pH 7.5) with the specified $Mg^{2+}$ concentrations. Each measurement was independently repeated three times.

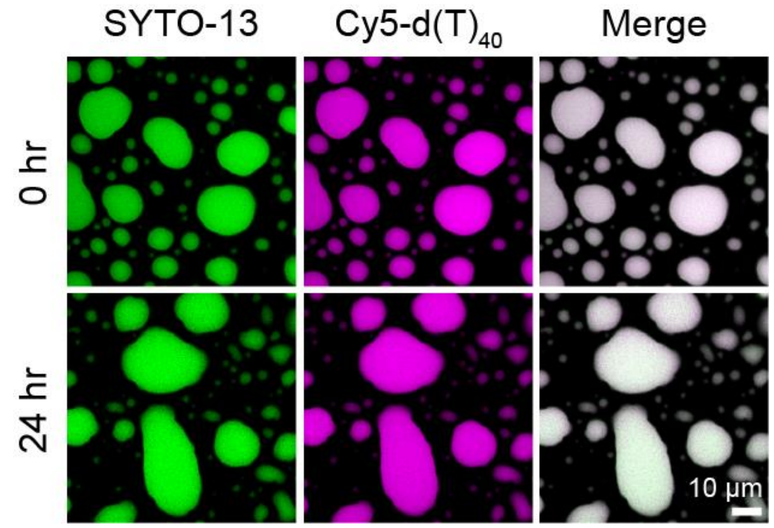


**Supplementary Figure 9.** Fluorescence images of (mut-TERRA)$_{10}$ containing RGG-d(T)$_{40}$ condensates show the absence of RNA clustering over a period of 24 hours. SYTO-13 and Cy5-d(T)$_{40}$ were used for condensate imaging. Also, see **Fig. 2g** and **Supplementary Fig. 10**. The composition of the condensate system used here is 1 mg/ml (mut-TERRA)$_{10}$ (corresponds to 51.8 µM), 5 mg/ml RGG, and 1.5 mg/ml d(T)$_{40}$ in a buffer containing 25 mM Tris-HCl (pH 7.5), 25 mM NaCl, and 20 mM DTT. The concentration range of the labeled components is 250 nM to 500 nM. This experiment was independently repeated three times.

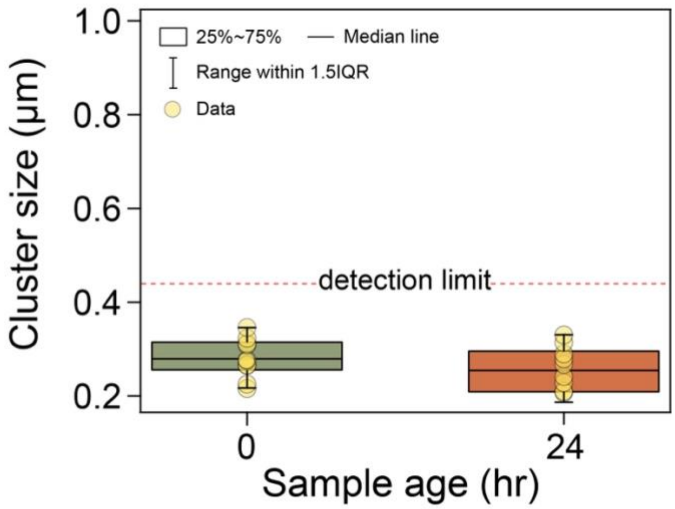


**Supplementary Figure 10.** Cluster size quantification using spatial autocorrelation analysis reveals the absence of time-dependent RNA cluster formation in $(\text{mut-TERRA})_{10}$ containing RGG–$d(T)_{40}$ condensates. This quantification is performed based on experiments independently repeated three times, as reported in **Fig. 2g** and **Supplementary Fig. 9**. The detection limit of SAC is demarcated (see Methods for further details). The box plot elements are defined in the figure.

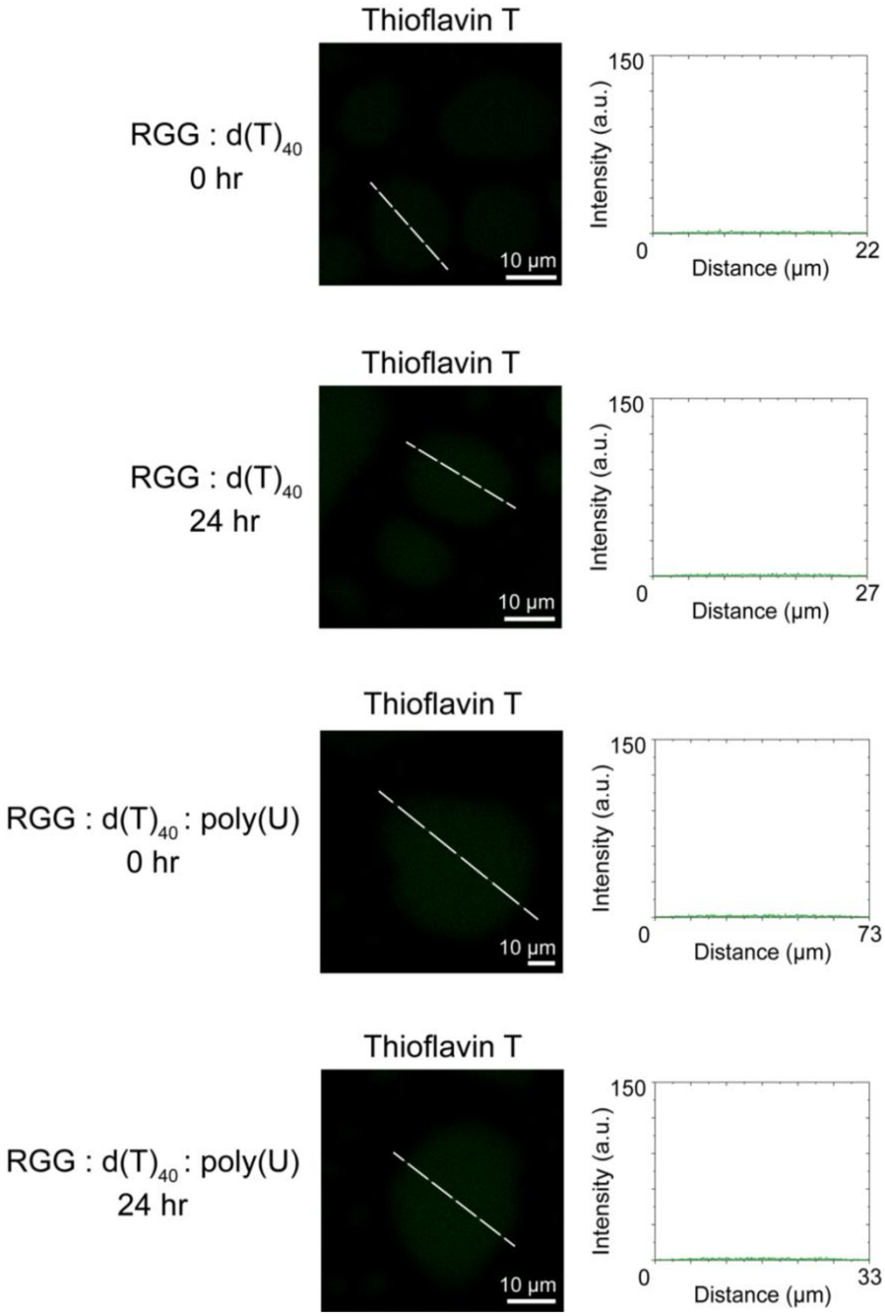


**Supplementary Figure 11.** Thioflavin T (ThT) staining of binary and ternary condensates, as indicated. Images captured at the initial and late time points of RGG-d(T)$_{40}$ and RGG-d(T)$_{40}$-poly(U) condensates do not show any appreciable amount of ThT fluorescence. The composition of the binary condensate system is 5 mg/ml RGG and 2.5 mg/ml d(T)$_{40}$. The composition of the ternary condensate system is 1 mg/ml RNA, 5 mg/ml RGG, and 1.5 mg/ml d(T)$_{40}$. The buffer composition is 25 mM Tris-HCl (pH 7.5), 25 mM NaCl, and 20 mM DTT. The concentration of ThT used is 50 µM. Each measurement was independently repeated three times.

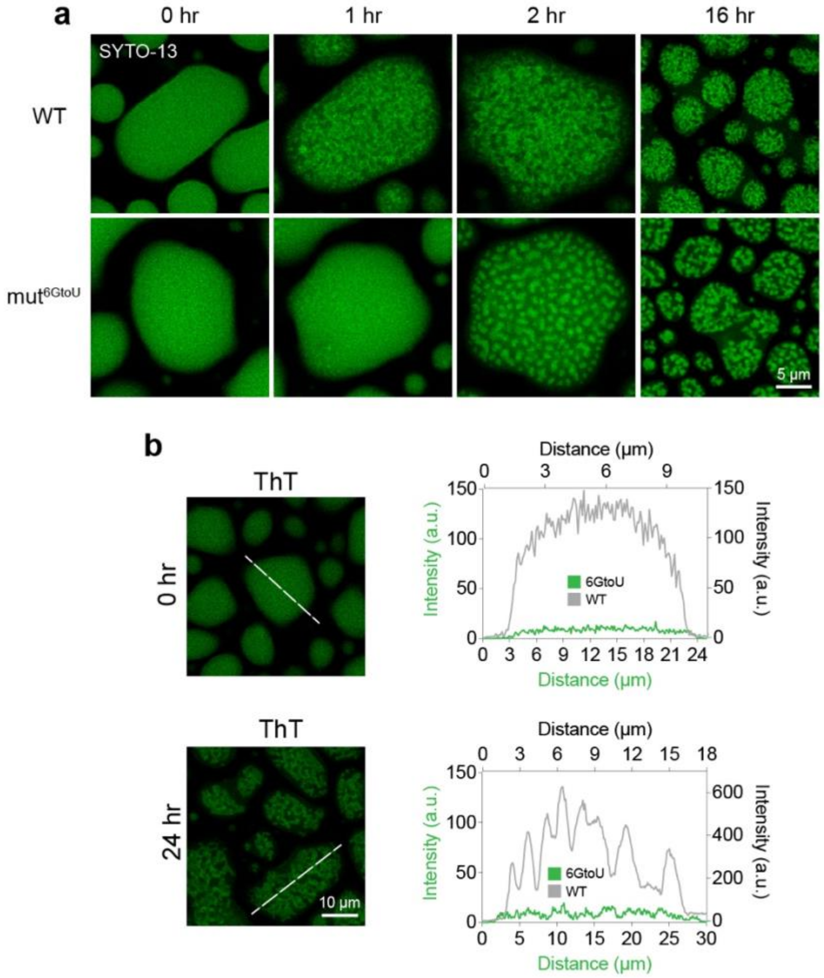


**Supplementary Figure 12.** (a) Fluorescence images of WT $(TERRA)_{10}$ and $(mut^{6GtoU}\text{-}TERRA)_{10}$ containing RGG-d$(T)_{40}$ ternary condensates at different time points after sample preparation. (b) Reports Thioflavin T (ThT) fluorescence images of $(mut^{6GtoU}\text{-}TERRA)_{10}$ containing RGG-d$(T)_{40}$ condensates at the indicated time points along with line profiles (shown in green) and corresponding ThT intensity profiles. The ThT intensity profiles for RGG-$(TERRA)_{10}$-d$(T)_{40}$ ternary condensates are shown (grey; data is taken from **Fig. 2h**) for comparison purposes. The composition of the condensate system used here is 1 mg/ml RNA [$(TERRA)_{10}$, 50.7 μM; $(mut^{6GtoU}\text{-}TERRA)_{10}$, 51.3 μM], 5 mg/ml RGG, and 1.5 mg/ml d$(T)_{40}$ in a buffer containing 25 mM Tris-HCl (pH 7.5), 25 mM NaCl, and 20 mM DTT. RNA clusters (or a lack thereof) were probed using 250 nM to 500 nM SYTO-13 in (a). The concentration of ThT used in (b) is 50 μM. These experiments were independently repeated three times.

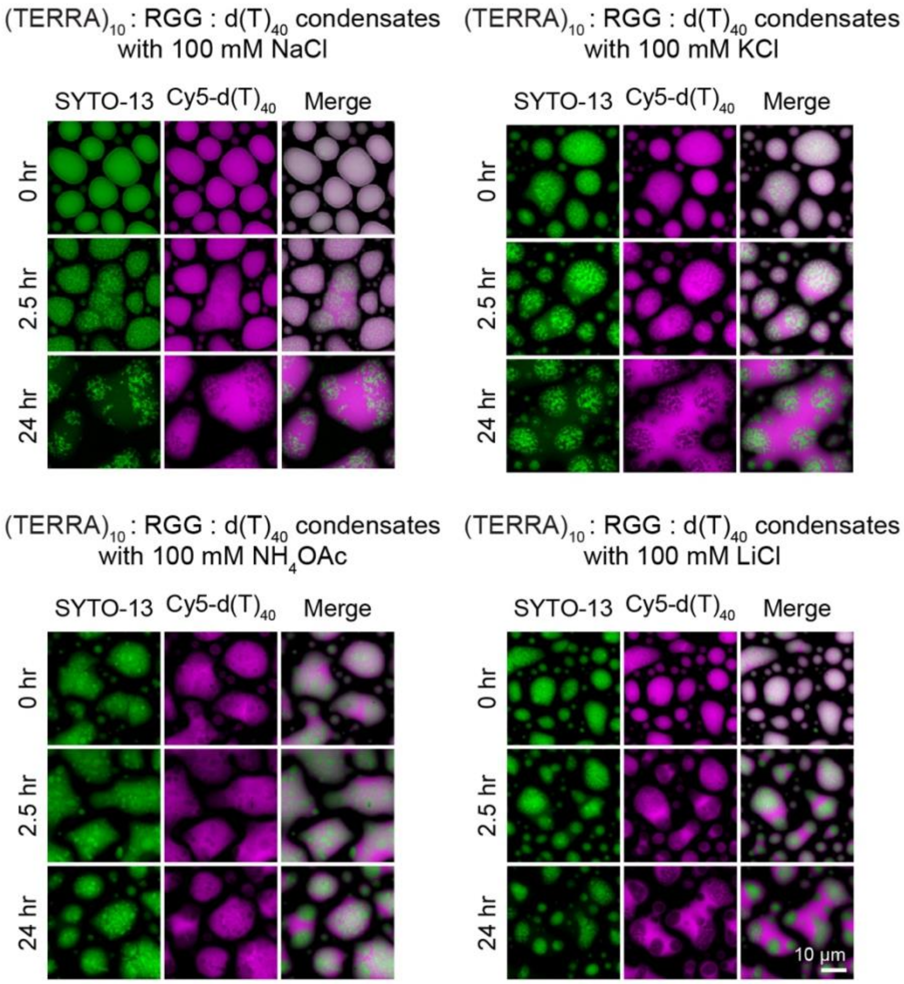


**Supplementary Figure 13.** Effect of various monovalent salts on RNA clustering in $(TERRA)_{10}$-RGG-$d(T)_{40}$ condensates. These salts can be classified as either weakly (such as LiCl) or strongly (such as KCl) stabilizing agents of RNA G-quadruplex structure formation[21]. The composition of the condensates is 1.0 mg/ml $(TERRA)_{10}$ (corresponds to 50.7 μM), 5.0 mg/ml RGG, 1.5 mg/ml $d(T)_{40}$ in a buffer containing 25 mM Tris-HCl (pH 7.5), 25 mM NaCl, 20 mM DTT along with the monovalent salts as indicated. The concentration range of labeled components is 250 nM to 500 nM. Each condition was independently repeated three times.

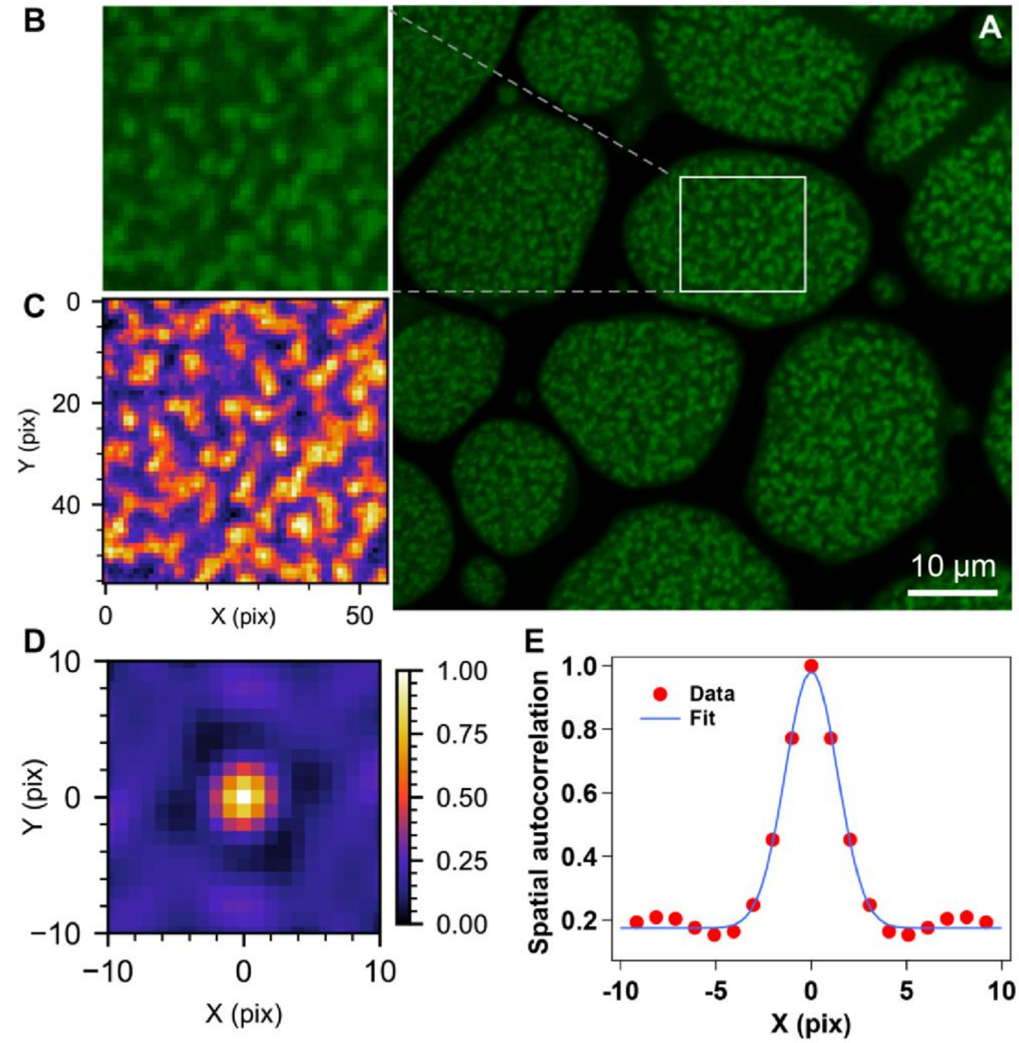


**Supplementary Figure 14.** Quantification of cluster sizes through 2D SAC analysis of fluorescence images. (a) Fluorescence image of $(TERRA)_{10}$ containing RGG-d$(T)_{40}$ condensates at 18 hours of age under the condition as reported in **Fig. 3a**. (b) Cropped image of a region inside a condensate chosen as input for SAC analysis. (c) Color-mapped depiction of the cropped region shown in (b). (d) 2D spatial autocorrelation functions plotted in *xy* plane. (e) Estimation of spatial autocorrelation along the *x*-axis with a Gaussian fit to estimate the cluster size, which is estimated to be 852 nm.

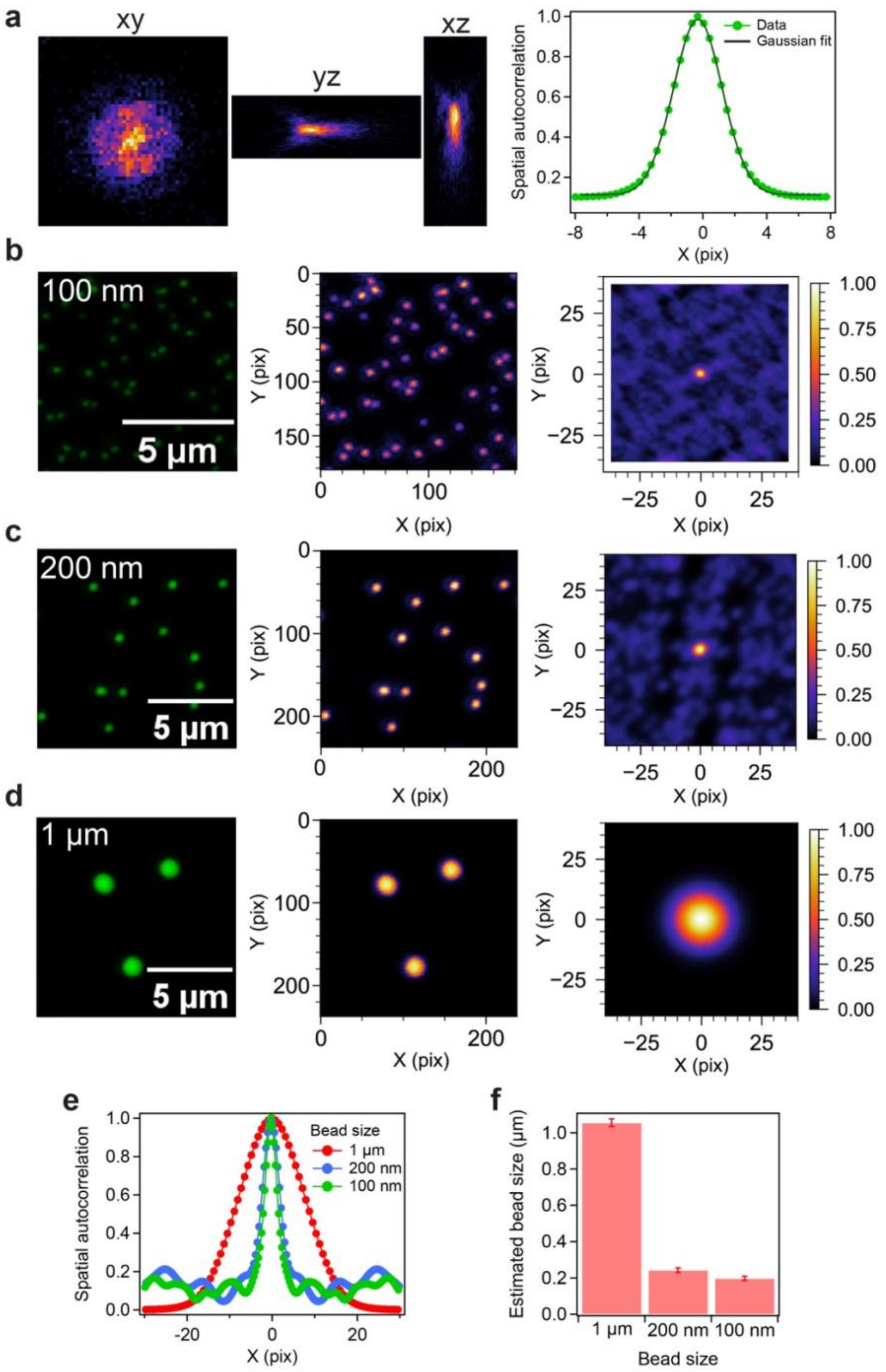


**Supplementary Figure 15.** Estimation of image resolution and probe particle sizes using SAC analysis. (a) Estimation of point spread function using 100 nm sized fluorescent probe particles. Fluorescence images (left) of (b) 100 nm, (c) 200 nm, and (d) 1 μm particles and corresponding color map images (middle) along with x-y spatial autocorrelation functions (right) from SAC. (e) SAC line profiles of the three different probe particles used in our studies. (f) Reports the estimated particle sizes. The mean estimated particle size of the '1 μm' particle is 1.056 μm, the '200 nm' particle is 243.5 nm, and the '100 nm' particle is 197.5 nm. Each measurement was independently repeated three times.

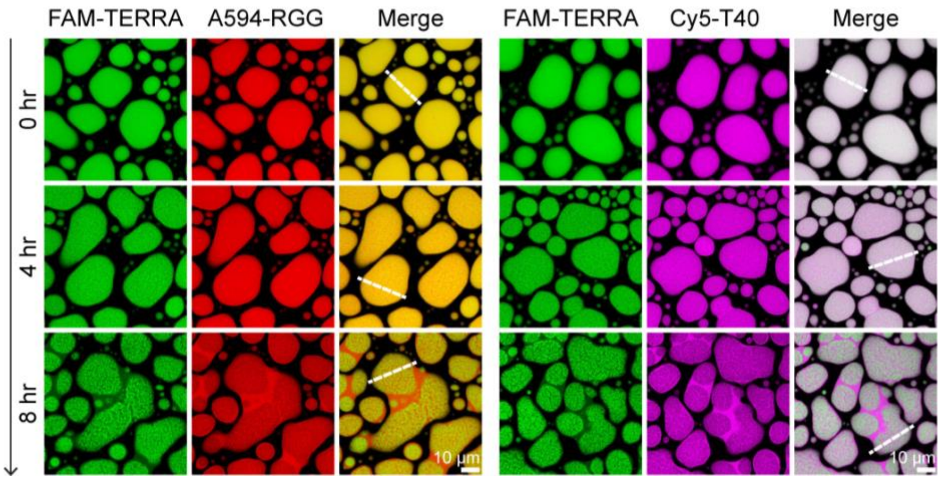


**Supplementary Figure 16.** Time-dependent demixing of $(TERRA)_4$ clusters from RGG and $d(T)_{40}$. The white dashed lines shown here correspond to line profile analyses shown in **Fig. 3e, f**. The composition of the condensate system used here is 1 mg/ml $(TERRA)_4$ (corresponds to 127 µM), 5 mg/ml RGG, and 1.5 mg/ml $d(T)_{40}$ in a buffer containing 25 mM Tris-HCl (pH 7.5), 25 mM NaCl, and 20 mM DTT. The concentration range of labeled components is 250 nM. Each measurement was independently repeated three times.

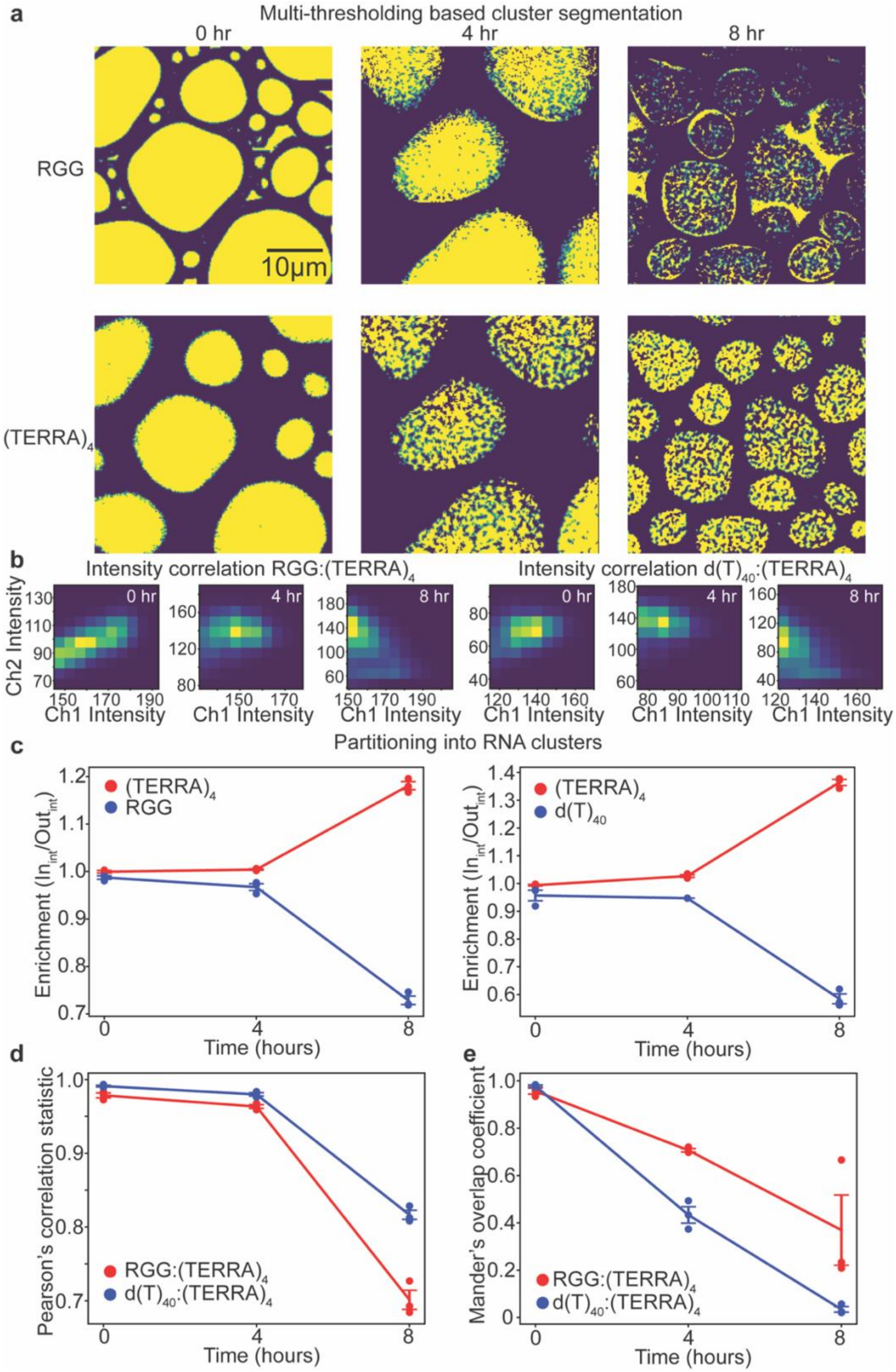


**Supplementary Figure 17.** (a) Multi-thresholding-based segmentation of $(TERRA)_4$ clusters in RGG-$d(T)_{40}$ condensates utilizing fluorescence images of Alexa594-labeled RGG and FAM-labeled $(TERRA)_4$ at three different time points after sample preparation. (b) Intensity correlation plots utilizing fluorescence images of FAM-labeled $(TERRA)_4$ (Ch1 intensity) and Alexa594-labeled RGG or Cy5-labeled $d(T)_{40}$ (Ch2 intensity) at three different time points after sample preparation. (c) (Left) Enrichment analysis of

Alexa594-labeled RGG versus FAM-labeled $(TERRA)_4$ in RNA clusters at three different time points after sample preparation. (Right) Enrichment analysis of Cy5-labeled $d(T)_{40}$ versus FAM-labeled $(TERRA)_4$ in RNA clusters at three different time points after sample preparation. (d) Pearson's correlation analysis of Alexa594-labeled RGG versus FAM-labeled $(TERRA)_4$ and Cy5-labeled $d(T)_{40}$ versus FAM-labeled $(TERRA)_4$. (e) Mander's overlap coefficient analysis of Alexa594-labeled RGG versus FAM-labeled $(TERRA)_4$ and Cy5-labeled $d(T)_{40}$ versus FAM-labeled $(TERRA)_4$. The composition of the condensate system used here is 1 mg/ml $(TERRA)_4$ (corresponds to 127 μM), 5 mg/ml RGG, and 1.5 mg/ml $d(T)_{40}$ in a buffer containing 25 mM Tris-HCl (pH 7.5), 25 mM NaCl, and 20 mM DTT. The concentration of labeled components is 250 nM. Each measurement was independently repeated three times except for the 4-hour time point, which was repeated two times. Error bars denote the standard error of the mean (S.E.M.).

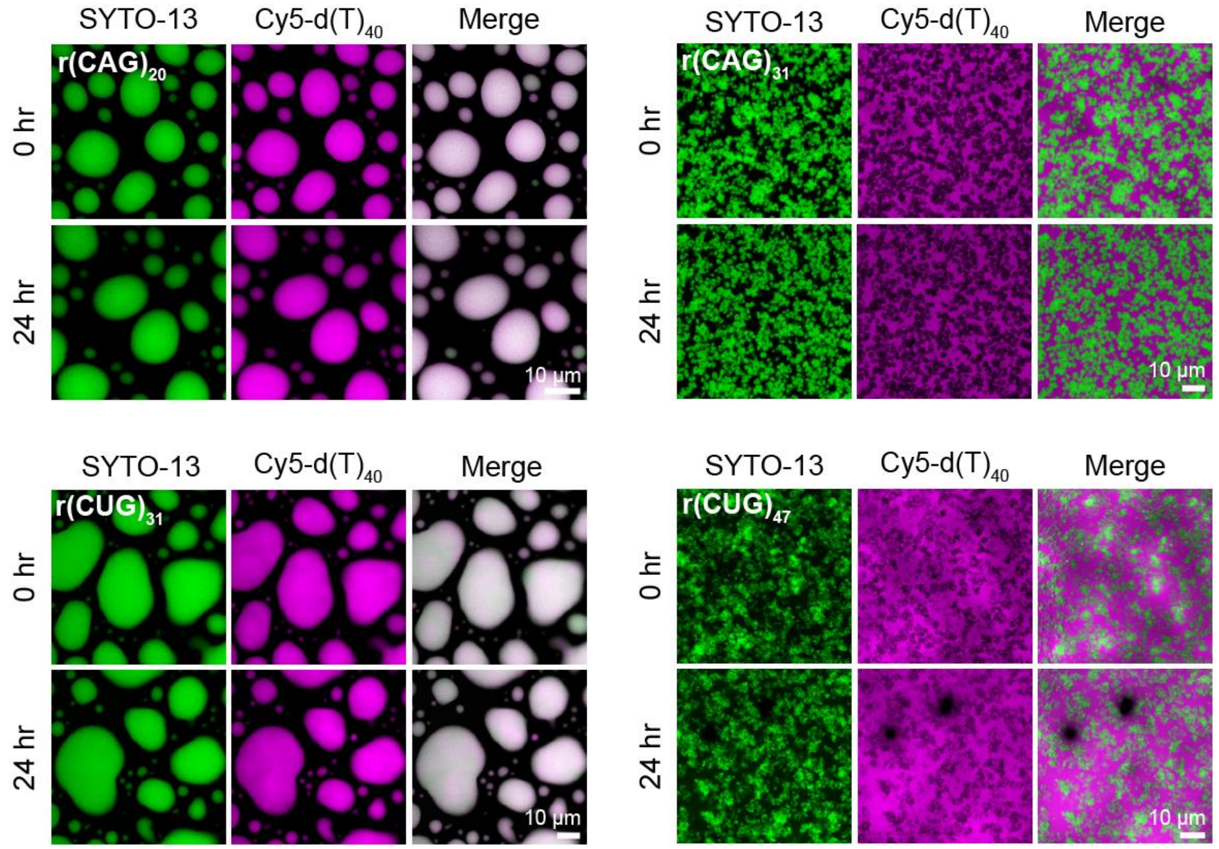


**Supplementary Figure 18.** Fluorescence images utilizing SYTO-13 and Cy5-d(T)$_{40}$ of RGG-d(T)$_{40}$ condensates containing either r(CAG) or r(CUG) repeat RNAs corresponding to data shown in **Fig. 4b-e**. The composition of the condensate system used here is 1 mg/ml RNA [0.45 mg/ml in the case of r(CUG)$_{47}$; r(CAG)$_{20}$, 51.2 µM; r(CAG)$_{31}$, 33 µM; r(CUG)$_{31}$, 33.8 µM; r(CUG)$_{47}$, 10 µM], 5 mg/ml RGG, and 1.5 mg/ml d(T)$_{40}$ in a buffer containing 25 mM Tris-HCl (pH 7.5), 25 mM NaCl, and 20 mM DTT. The concentration range of labeled components is 250 nM to 500 nM. Each measurement was independently repeated three times.

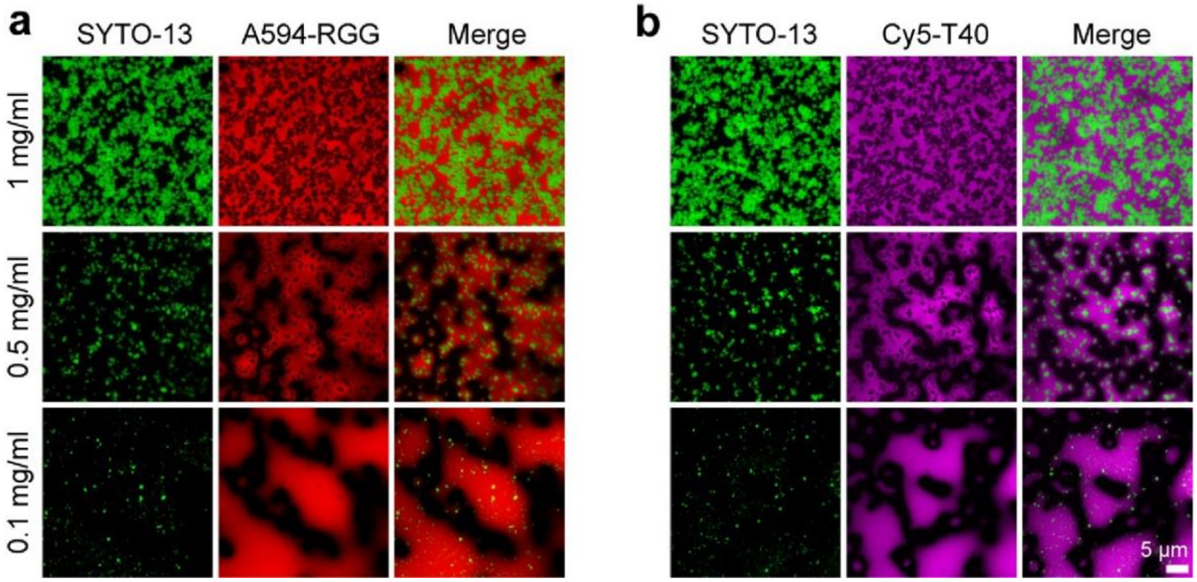


**Supplementary Figure 19.** (a) Fluorescence images utilizing SYTO-13 and A594-RGG of RGG-d(T)$_{40}$ condensates containing r(CAG)$_{31}$ at different concentrations imaged 15 minutes after sample preparation (1 mg/ml data corresponds to data shown in **Fig. 4b**). (b) Fluorescence images utilizing SYTO-13 and Cy5-d(T)$_{40}$ of RGG-d(T)$_{40}$ condensates containing r(CAG)$_{31}$ at different concentrations imaged 15 minutes after sample preparation. The composition of the condensate system used here is 5 mg/ml RGG, variable d(T)$_{40,}$ and r(CAG)$_{31}$ concentration as indicated [1 mg/ml r(CAG)$_{31}$, 33 μM; 0.5 mg/ml r(CAG)$_{31}$,16.5 μM; 0.1 mg/ml r(CAG)$_{31}$, 3.3 μM] while keeping total nucleic acid concentration as 2.5 mg/ml, in a buffer containing 25 mM Tris-HCl (pH 7.5), 25 mM NaCl, and 20 mM DTT. The concentration range of labeled components is 250 nM to 500 nM. Each measurement was independently repeated three times.

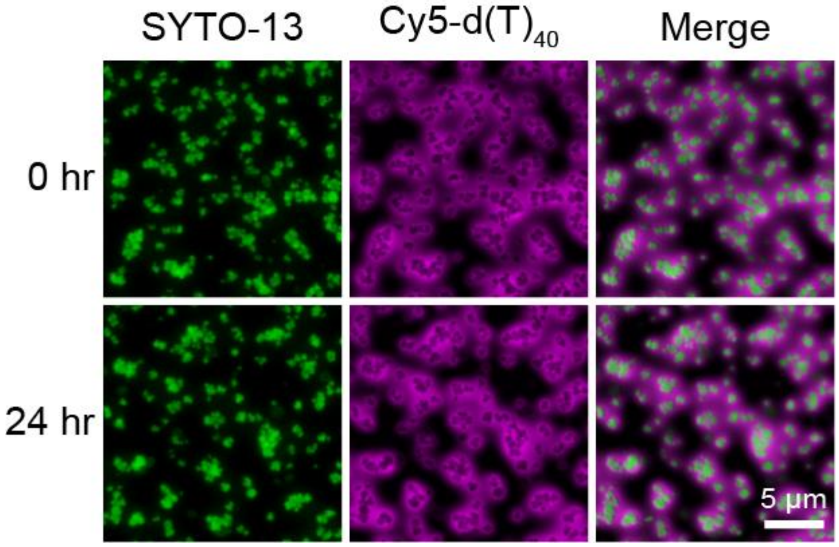


**Supplementary Figure 20.** r(GGGGCC)$_5$ forms clusters in RGG-d(T)$_{40}$ condensates. Imaging was performed 15 minutes after sample preparation. The composition of the condensates used here is 0.2 mg/ml r(GGGGCC)$_5$ (corresponds to 20.2 μM), 1.0 mg/ml RGG, 0.3 mg/ml d(T)$_{40}$ in a buffer containing 25 mM Tris-HCl (pH 7.5), 25 mM NaCl, 20 mM DTT. The concentration range of labeled components is 250 nM to 500 nM. This experiment was independently repeated three times.

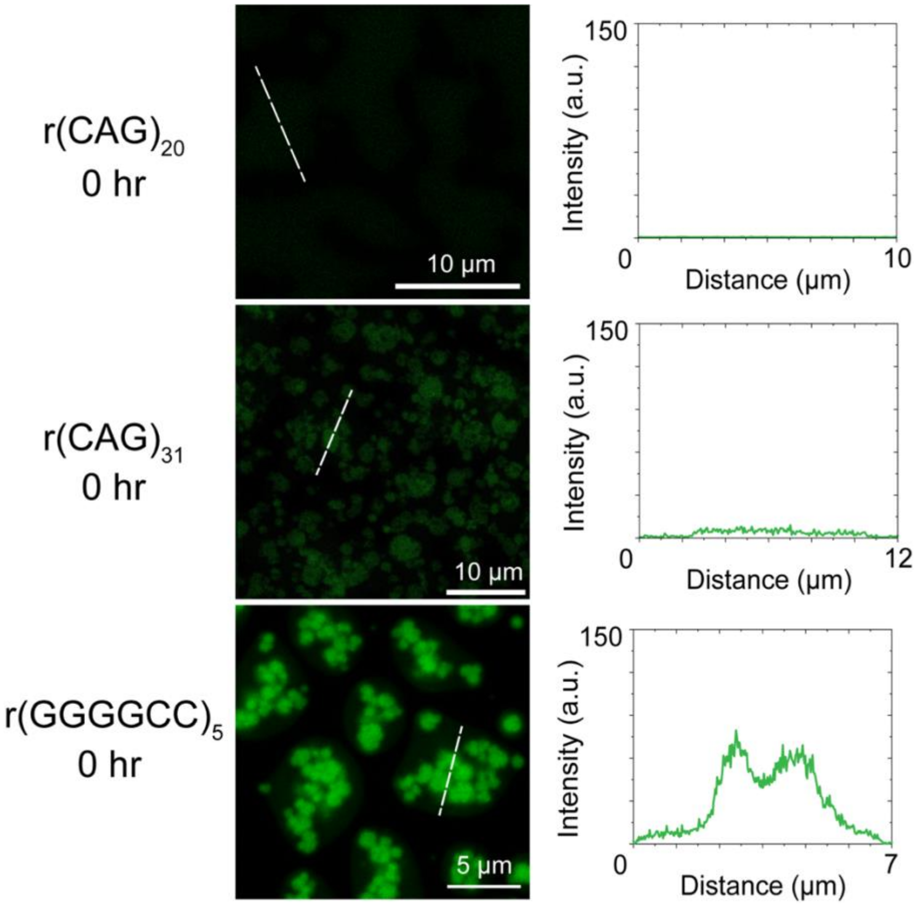


**Supplementary Figure 21.** Thioflavin T (ThT) fluorescence images (left) and profiles (right) of RGG-d(T)$_{40}$ condensates containing r(CAG)$_{20}$, r(CAG)$_{31,}$ or r(GGGGCC)$_5$, as indicated. The composition of the r(CAG)$_{20}$ and r(CAG)$_{31}$ condensate samples is 1 mg/ml RNA [r(CAG)$_{20}$, 51.2 μM; r(CAG)$_{31}$, 33 μM;], 5 mg/ml RGG, and 1.5 mg/ml d(T)$_{40}$ in a buffer containing 25 mM Tris-HCl (pH 7.5), 25 mM NaCl, and 20 mM DTT. The composition of the r(GGGGCC)$_5$ condensate sample is 0.2 mg/ml RNA (corresponds to 20.2 μM), 1 mg/ml RGG, and 0.3 mg/ml d(T)$_{40}$. The concentration of ThT used is 50 μM. Each measurement was independently repeated three times.

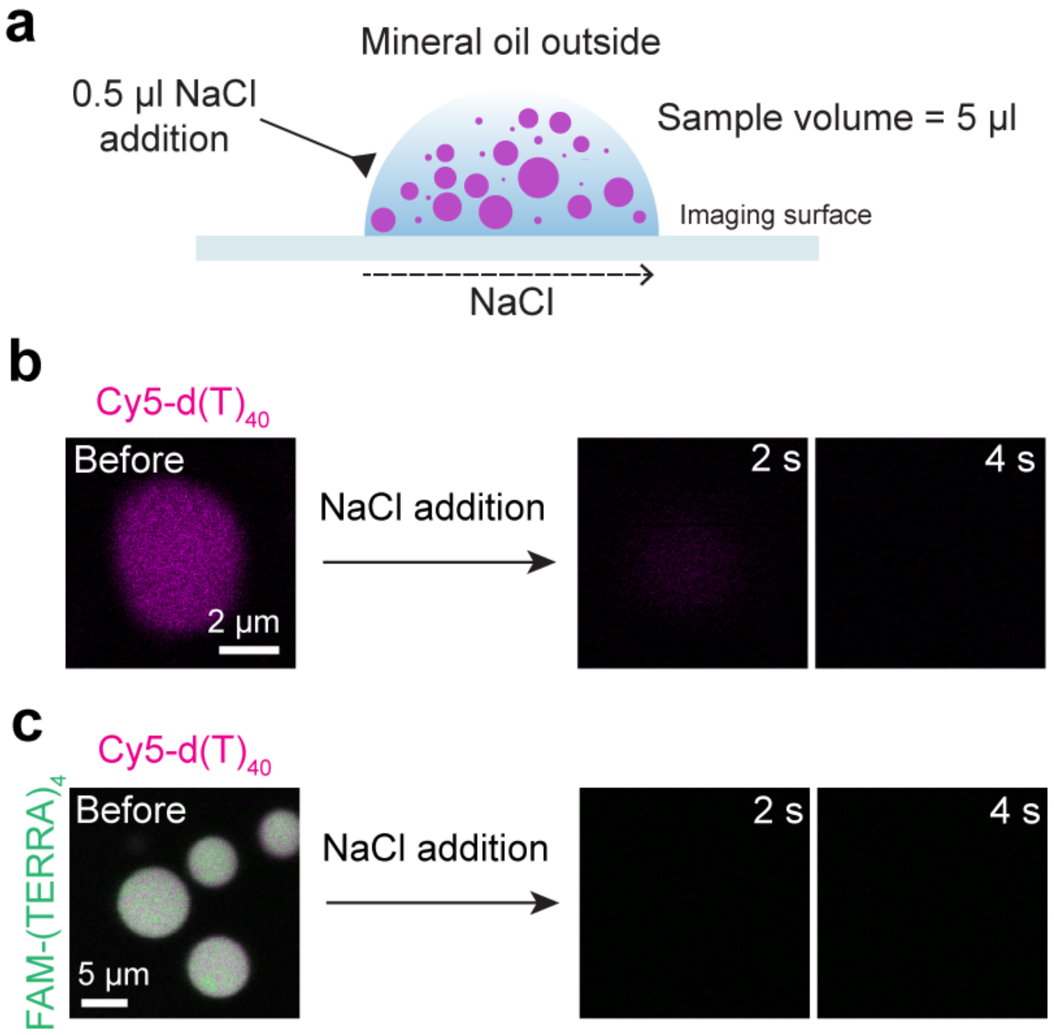


**Supplementary Figure 22.** (a) A schematic depicting condensate dissolution assay using NaCl. (b) Dissolution of freshly prepared binary RGG-d(T)$_{40}$ condensate by addition of 0.5 μl of 5 M NaCl. The sample composition is 1 mg/ml RGG, and 1 mg/ml d(T)$_{40}$ in 25 mM Tris-HCl (pH 7.5), 25 mM NaCl, and 20 mM DTT. The condensate was visualized using 250 nM Cy5-labeled d(T)$_{40}$. (c) Dissolution of freshly prepared (TERRA)$_{10}$ containing RGG-d(T)$_{40}$ condensate by addition of 0.5 μl of 5 M NaCl. The sample composition is 2.5 mg/ml RGG, 0.75 mg/ml d(T)$_{40}$, and 0.5 mg/ml (TERRA)$_{10}$ (corresponds to 25.4 μM) in 25 mM Tris-HCl (pH 7.5), 25 mM NaCl, and 20 mM DTT. The concentration of labeled components is 250 nM. Each experiment was independently repeated three times.

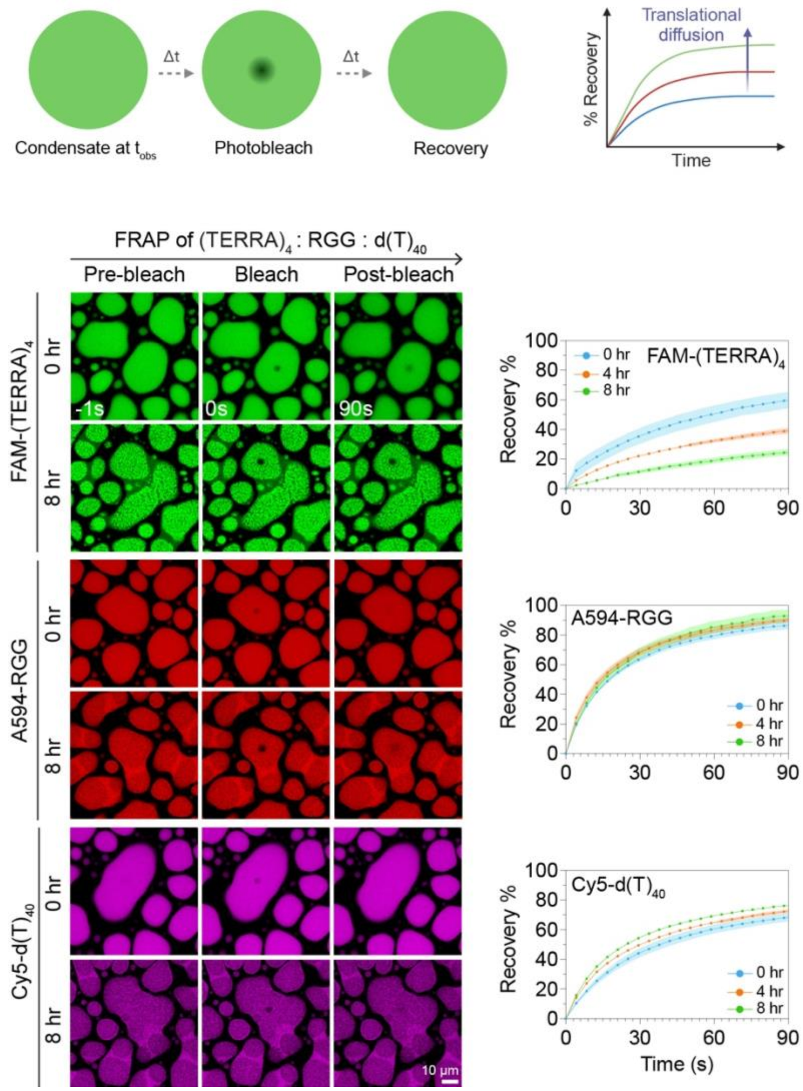


**Supplementary Figure 23.** Fluorescence recovery after photobleaching (FRAP) reveals dynamical arrest of the RNA component, TERRA, but not RGG and $d(T)_{40}$ with condensate aging. (Top) Schematic of FRAP assay in condensate. (Bottom) Fluorescence images of $(TERRA)_4$ containing RGG-$d(T)_{40}$ condensates at pre-bleach, bleach, and post-bleach steps of FRAP experiments corresponding to **Fig. 5c** (identical FRAP profiles are also shown here) and **Supplementary Videos 11-16**. Shaded regions in each plot signify the standard error. The composition of the $(TERRA)_4$ containing RGG-$d(T)_{40}$ condensate system used here is 1 mg/ml $(TERRA)_4$ (corresponds to 127 µM), 5 mg/ml RGG, and 1.5 mg/ml $d(T)_{40}$ in a buffer containing 25 mM Tris-HCl (pH 7.5), 25 mM NaCl, and 20 mM DTT. The concentration of labeled components is 250 nM. Each experiment was independently repeated three times.

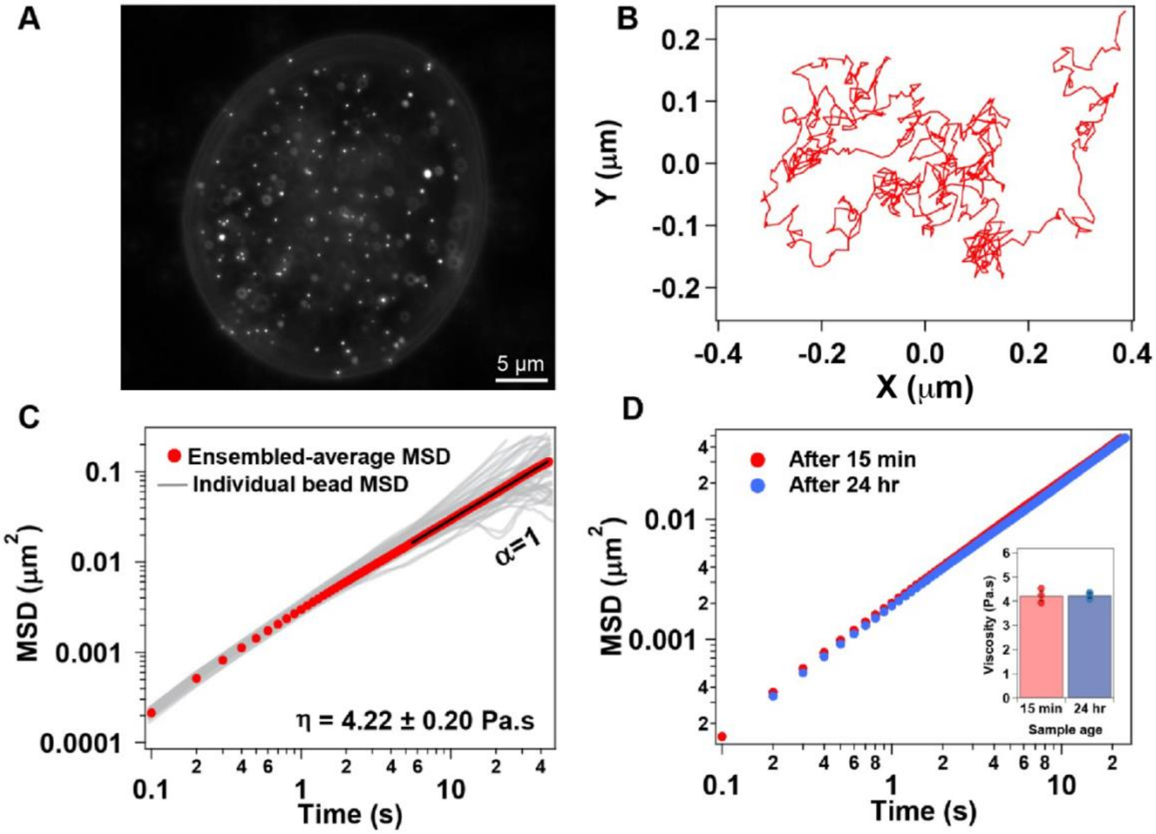


**Supplementary Figure 24.** VPT-based nanorheology of RGG-d(T)$_{40}$ condensates. (a) The initial frame of the video that was used for VPT analysis of 200 nm fluorescent carboxylate-modified polystyrene beads passively embedded inside an RGG-d(T)$_{40}$ condensate. (b) Representative trajectory of a bead inside the condensate. (c) Ensemble-average MSD overlaid over individual trajectories of the beads tracked in (a). Diffusivity coefficient, α=1 at longer lag times indicates normal diffusion of the beads inside RGG-d(T)$_{40}$ condensates, allowing estimation of terminal viscosity of η = 4.22 ± 0.20 Pa.s. (d) Overlaid ensemble-average MSDs of the beads at 15 minutes and 24 hours after sample preparation. There is no substantial change in the ensemble-average MSDs and corresponding viscosities (inset) of a sample at 15 minutes and after 24 hours of aging, suggesting that RGG-d(T)$_{40}$ condensates do not show any changes in material properties during the 24-hour observation period. Error bars denote the standard deviation. The composition of the condensate system is 5 mg/ml RGG and 5 mg/ml d(T)$_{40}$ in a buffer containing 25 mM MOPS, 25 mM NaCl, and 20 mM DTT. Each measurement was independently repeated three times.

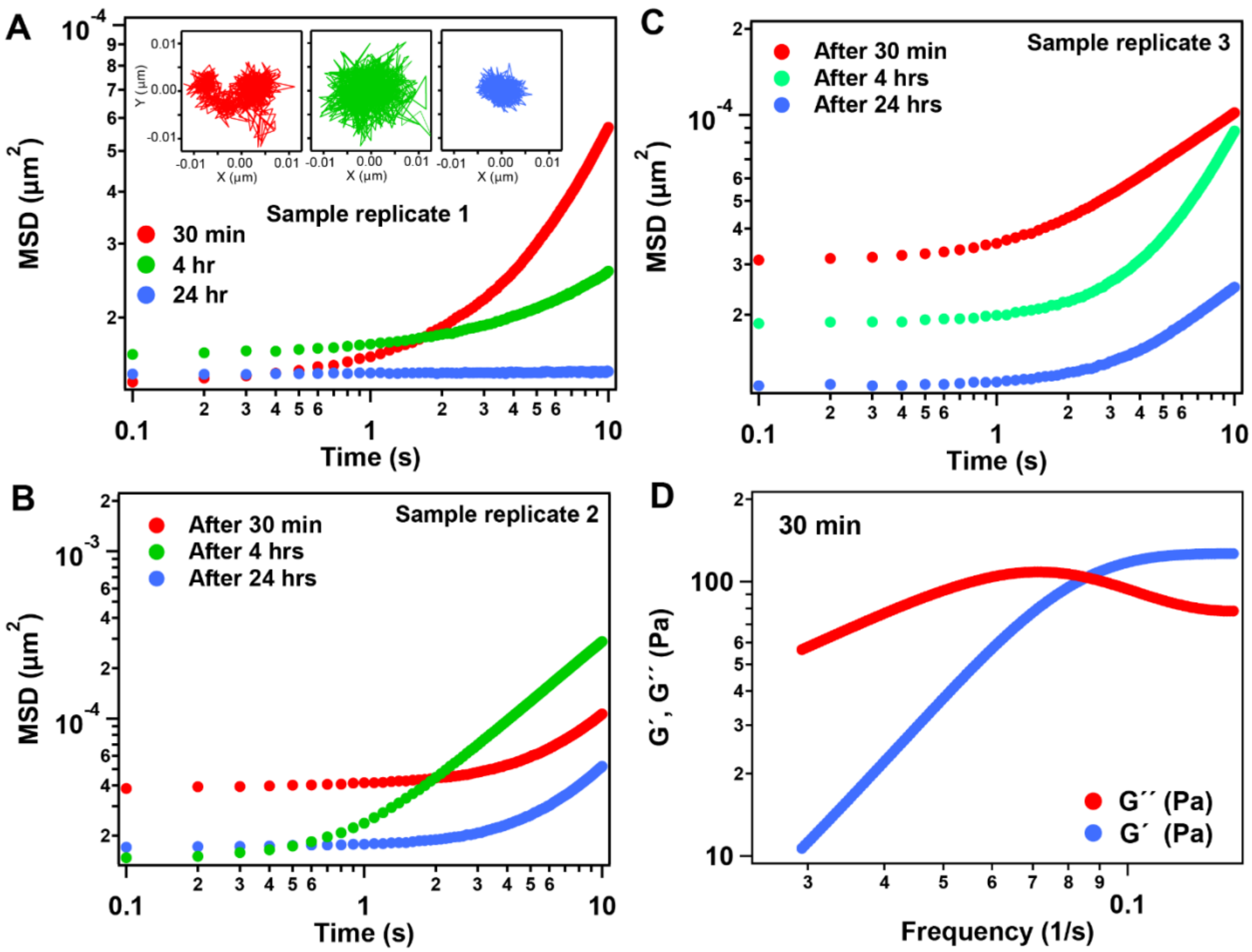


**Supplementary Figure 25.** (a-c) MSD plots for three replicates of RGG-$(TERRA)_{10}$ condensates. The MSD plots were derived from VPT analysis of 200 nm carboxylate-modified fluorescent polystyrene beads passively embedded inside RGG-$(TERRA)_{10}$ condensates. Insets shown in (a) represent the trajectories of the probe particles at the indicated time points. (d) Estimated dynamical moduli of RGG-$(TERRA)_{10}$ condensates at the indicated time point. The composition of the condensate system is 5 mg/ml $(TERRA)_{10}$ (corresponds to 254 μM) and 10 mg/ml RGG. Buffer composition is 25 mM Tris-HCl (pH 7.5), 25 mM NaCl, and 20 mM DTT.

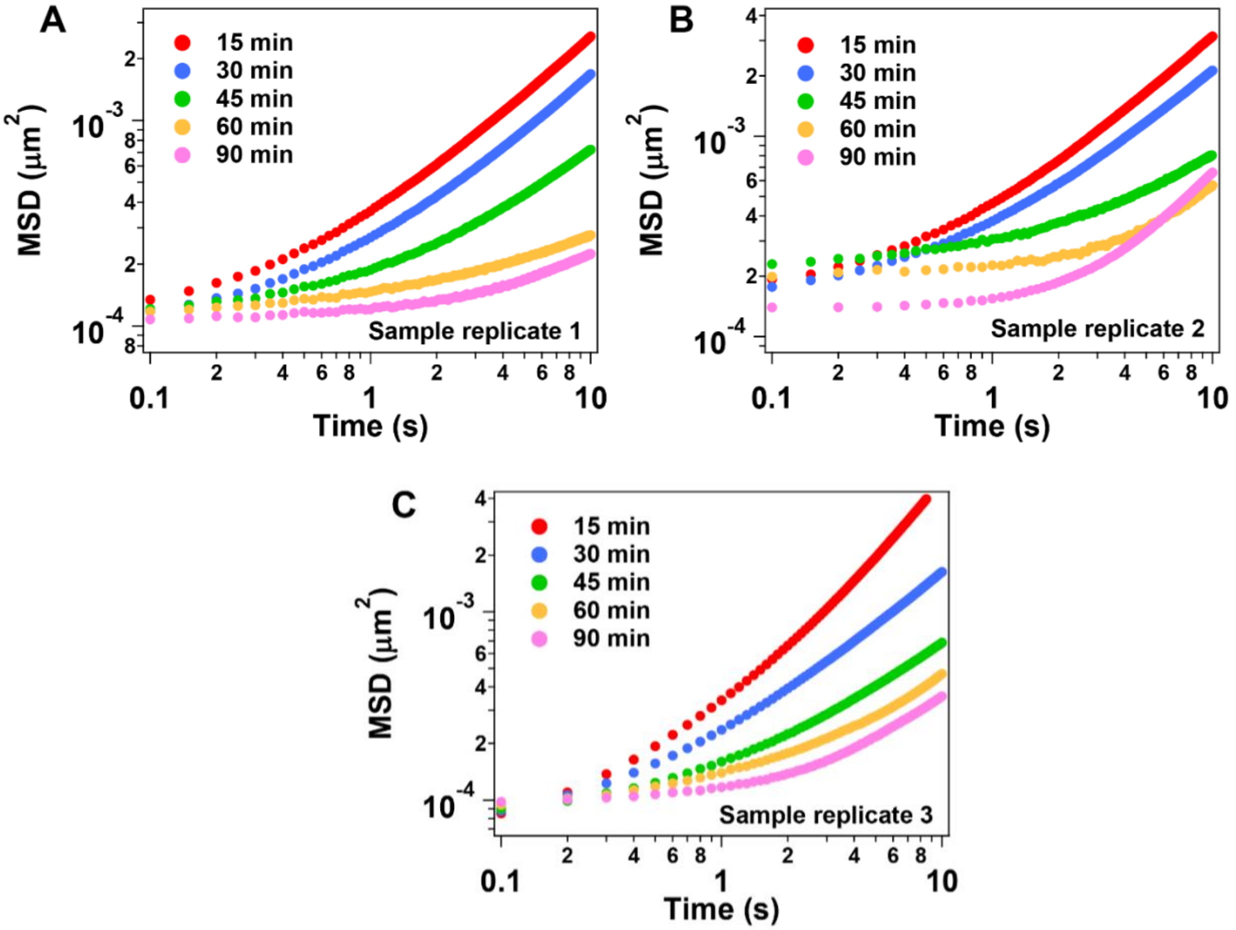


**Supplementary Figure 26.** (a-c) MSD plots for three replicates of (TERRA)$_{10}$ containing RGG-d(T)$_{40}$ condensates. These measurements were used to estimate the viscosities reported in **Fig. 5h**.

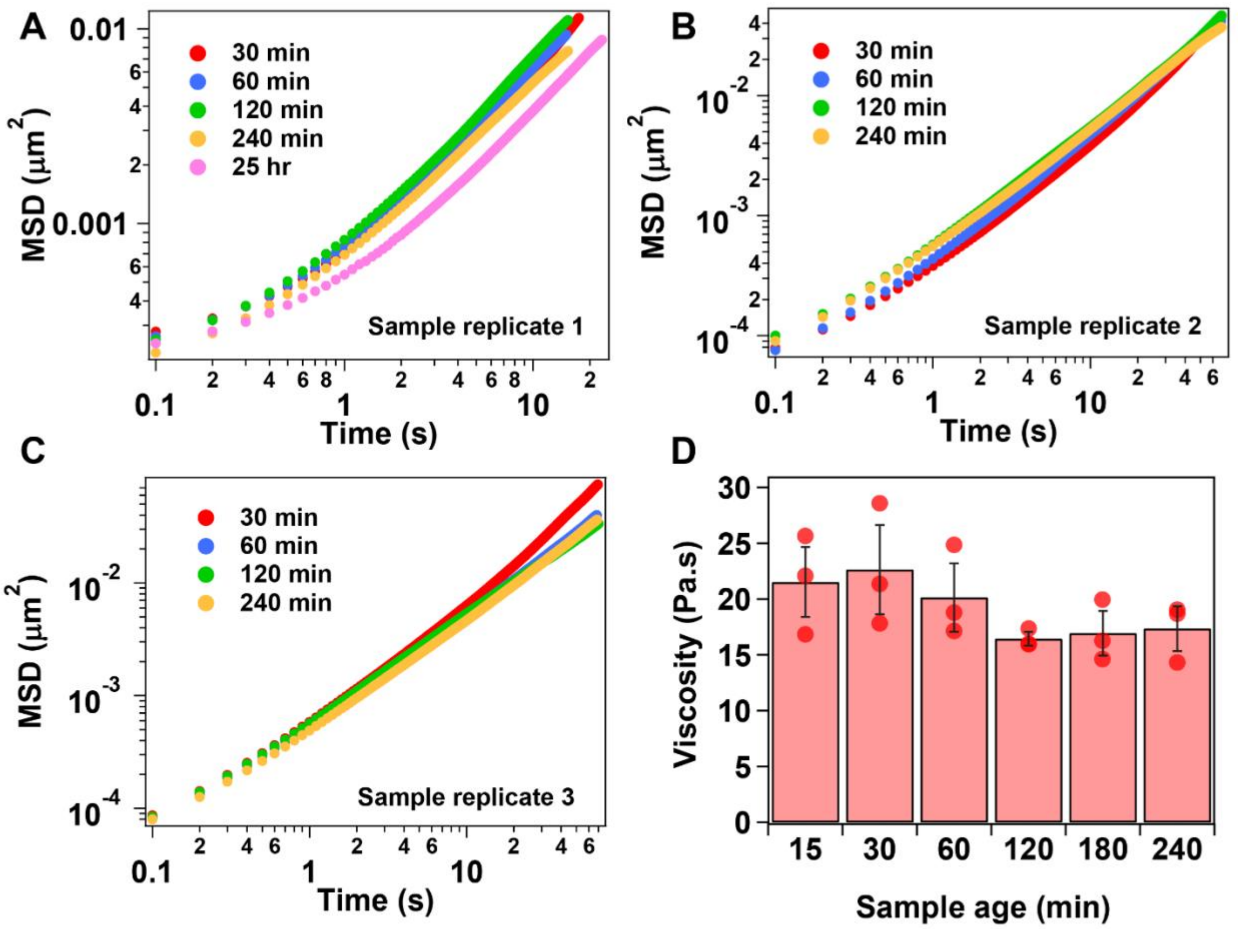


**Supplementary Figure 27.** (a-c) MSD plots for three replicates of $(\text{mut-TERRA})_{10}$ containing RGG-$d(T)_{40}$ condensates. (d) Estimated viscosities from the MSDs shown in (a-c), which is also reported in **Fig. 5j**. Error bars denote the standard deviation.

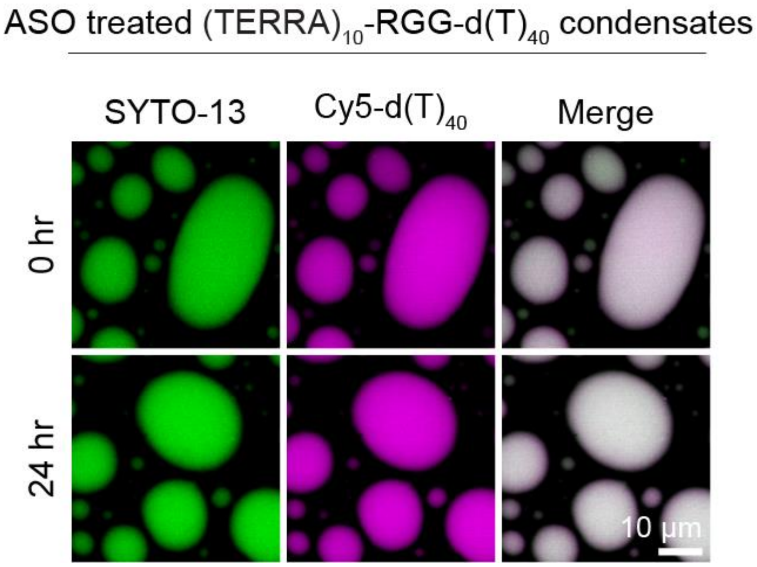


**Supplementary Figure 28.** Fluorescence images utilizing SYTO-13 and Cy5-d(T)$_{40}$ of (TERRA)$_{10}$ containing RGG-d(T)$_{40}$ condensates treated with TERRA antisense oligonucleotide [ASO; sequence: r(CCCUAA)]. The sample conditions are the same as the data reported in **Fig. 6b, c**. The composition of the condensates is 1.0 mg/ml (TERRA)$_{10}$ (corresponds to 50.7 μM), 5.0 mg/ml RGG, 1.5 mg/ml d(T)$_{40}$ in a buffer containing 25 mM Tris-HCl (pH 7.5), 25 mM NaCl, 20 mM DTT. 1.0 mg/ml ASO was added to the buffer prior to condensate formation. The concentration range of labeled components is 250 nM to 500 nM. This experiment was independently repeated three times.

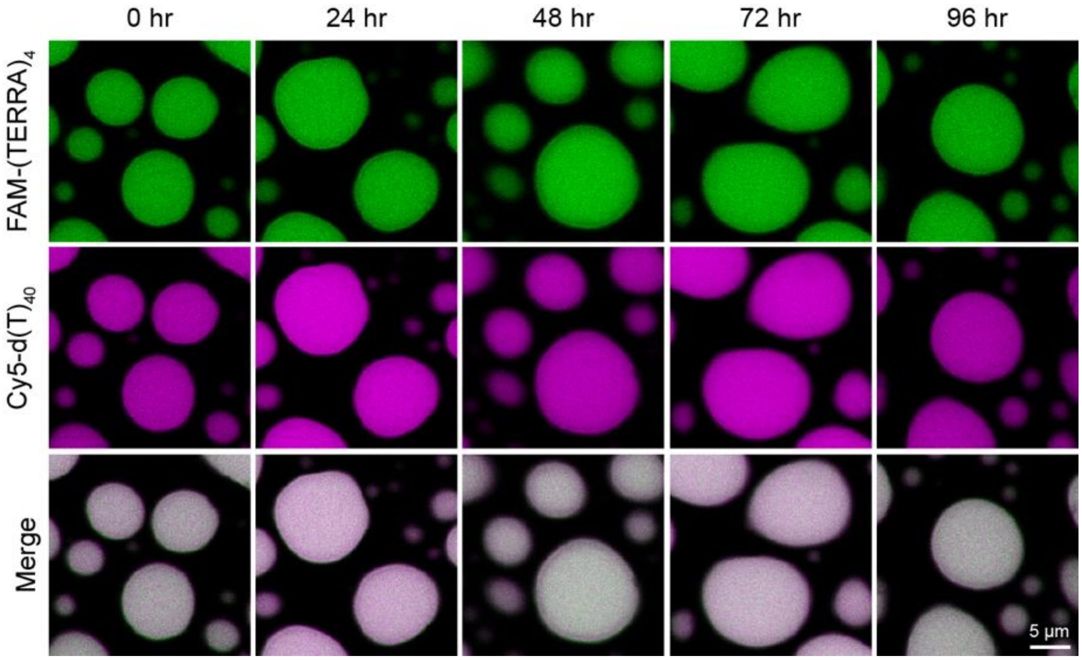


**Supplementary Figure 29.** Fluorescence images utilizing FAM-$(TERRA)_4$ and Cy5-$d(T)_{40}$ of $(TERRA)_{10}$ containing RGG-$d(T)_{40}$ condensates treated with TERRA antisense oligonucleotide [ASO; sequence: r(CCCUAA)]. The sample conditions are the same as the data reported in **Fig. 6b, c**. The composition of the condensates is 1.0 mg/ml $(TERRA)_{10}$ (corresponds to 50.7 µM), 5.0 mg/ml RGG, 1.5 mg/ml $d(T)_{40}$ in a buffer containing 25 mM Tris-HCl (pH 7.5), 25 mM NaCl, 20 mM DTT. 1.0 mg/ml ASO was added to the buffer prior to condensate formation. The concentration range of labeled components is 250 nM to 500 nM. This experiment was independently repeated three times.

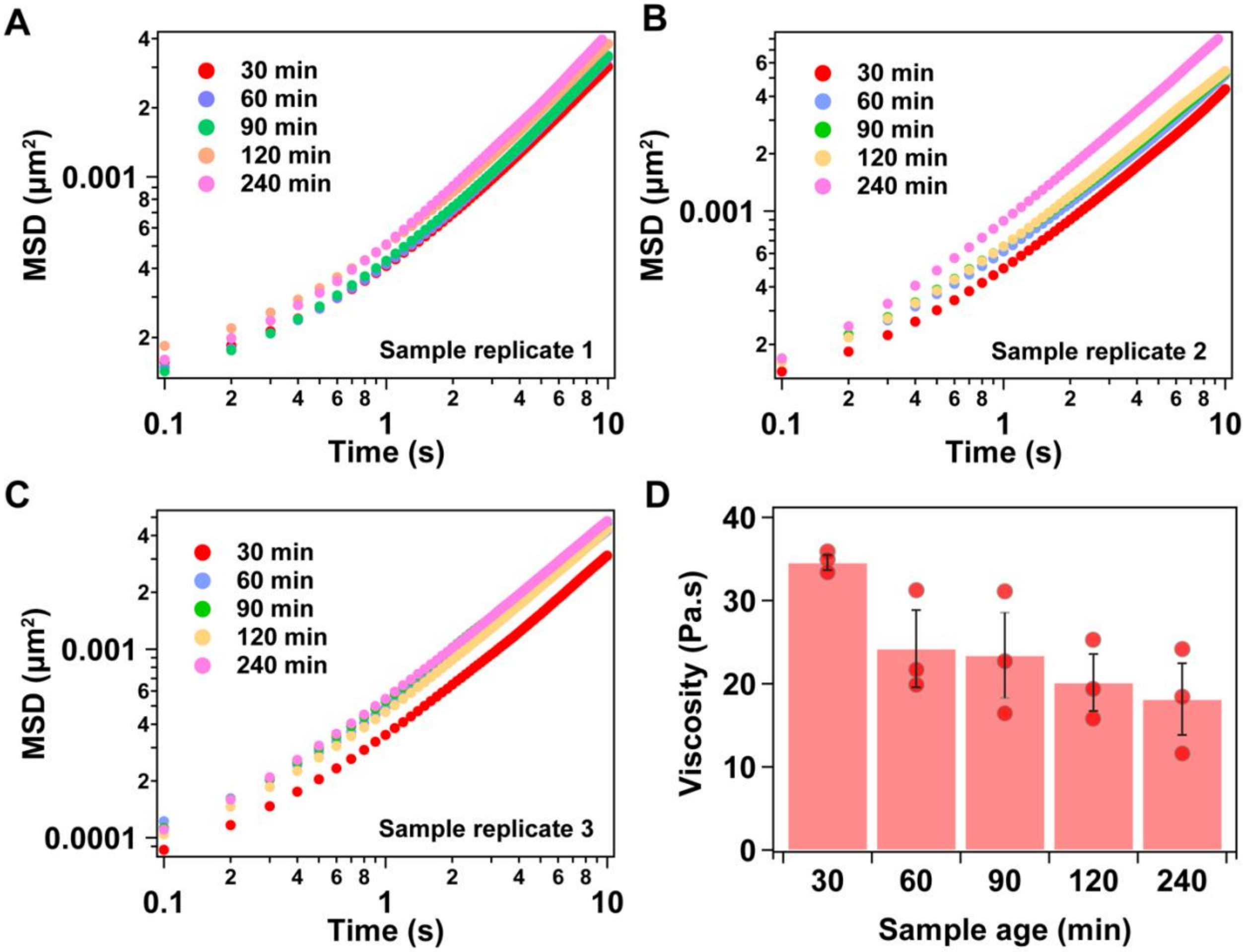


**Supplementary Figure 30.** (a-c) MSD plots for three replicates of $(TERRA)_{10}$ containing RGG-$d(T)_{40}$ condensates treated with ASO. (d) Estimated viscosities from the MSDs shown in (a-c), which is also reported in **Fig. 6j**. Error bars denote the standard deviation.

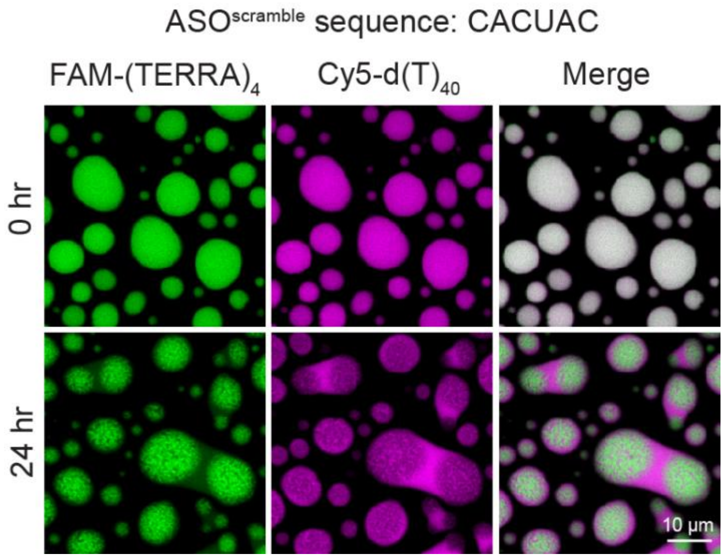


**Supplementary Figure 31.** Fluorescence images utilizing FAM-$(TERRA)_4$ and Cy5-$d(T)_{40}$ of $(TERRA)_{10}$ containing RGG-$d(T)_{40}$ condensates treated with a scrambled version of TERRA antisense oligonucleotide (ASO): CACUAC. The composition of the condensates is 1.0 mg/ml $(TERRA)_{10}$ (corresponds to 50.7 µM), 5.0 mg/ml RGG, 1.5 mg/ml $d(T)_{40}$ in a buffer containing 25 mM Tris-HCl (pH 7.5), 25 mM NaCl, 20 mM DTT. 1.0 mg/ml ASO was added to the buffer prior to condensate formation. The concentration range of labeled components is 250 nM to 500 nM. This experiment was independently repeated three times.

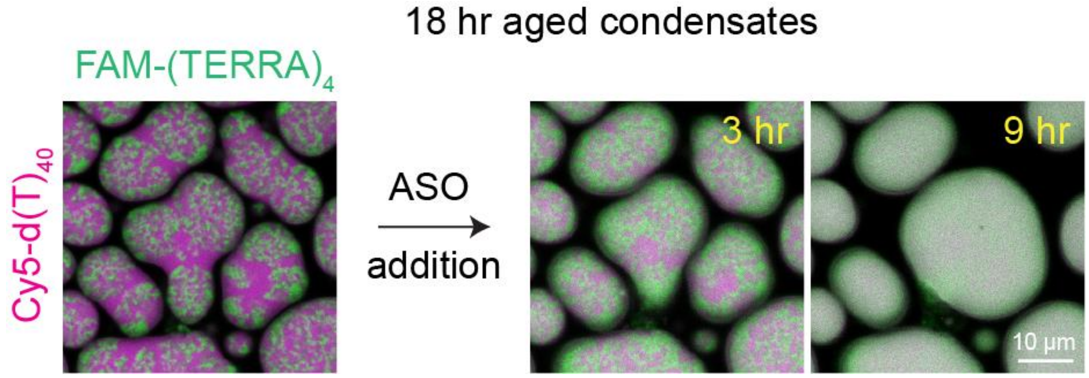


**Supplementary Figure 32.** Addition of ASO [sequence: r(CCCUAA)] to $(TERRA)_{10}$ containing RGG-$d(T)_{40}$ condensates after the formation of RNA clusters at 18 hours since sample preparation. These observations correspond to **Supplementary Video 21**. The composition of the ternary condensate system is 0.5 mg/ml RNA (corresponds to 25.4 μM), 2.5 mg/ml RGG, and 0.75 mg/ml $d(T)_{40}$. The buffer composition is 25 mM Tris-HCl (pH 7.5), 25 mM NaCl, and 20 mM DTT. The concentration of ASO used is 0.5 mg/ml. The concentration of labeled components is 250 nM. This measurement was independently repeated three times.

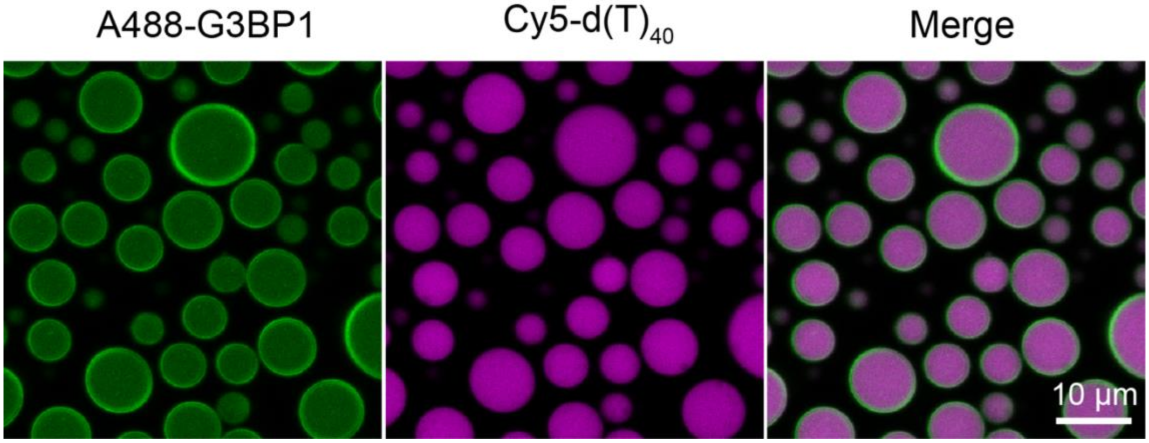


**Supplementary Figure 33.** Representative fluorescence images of RGG-(TERRA)$_{10}$-d(T)$_{40}$-G3BP1 condensates utilizing Alexa488-labeled G3BP1 (A488-G3BP1) and Cy5-d(T)$_{40}$. Images taken 15 minutes after sample preparation. The composition of the condensate system used here is 10 μM G3BP1, 1 mg/ml RNA (corresponds to 50.7 μM), 5 mg/ml RGG, and 1.5 mg/ml d(T)$_{40}$ in a buffer containing 25 mM Tris-HCl (pH 7.5), 25 mM NaCl, and 20 mM DTT. The concentration range of labeled components is 250 nM to 500 nM. This experiment was repeated three times.

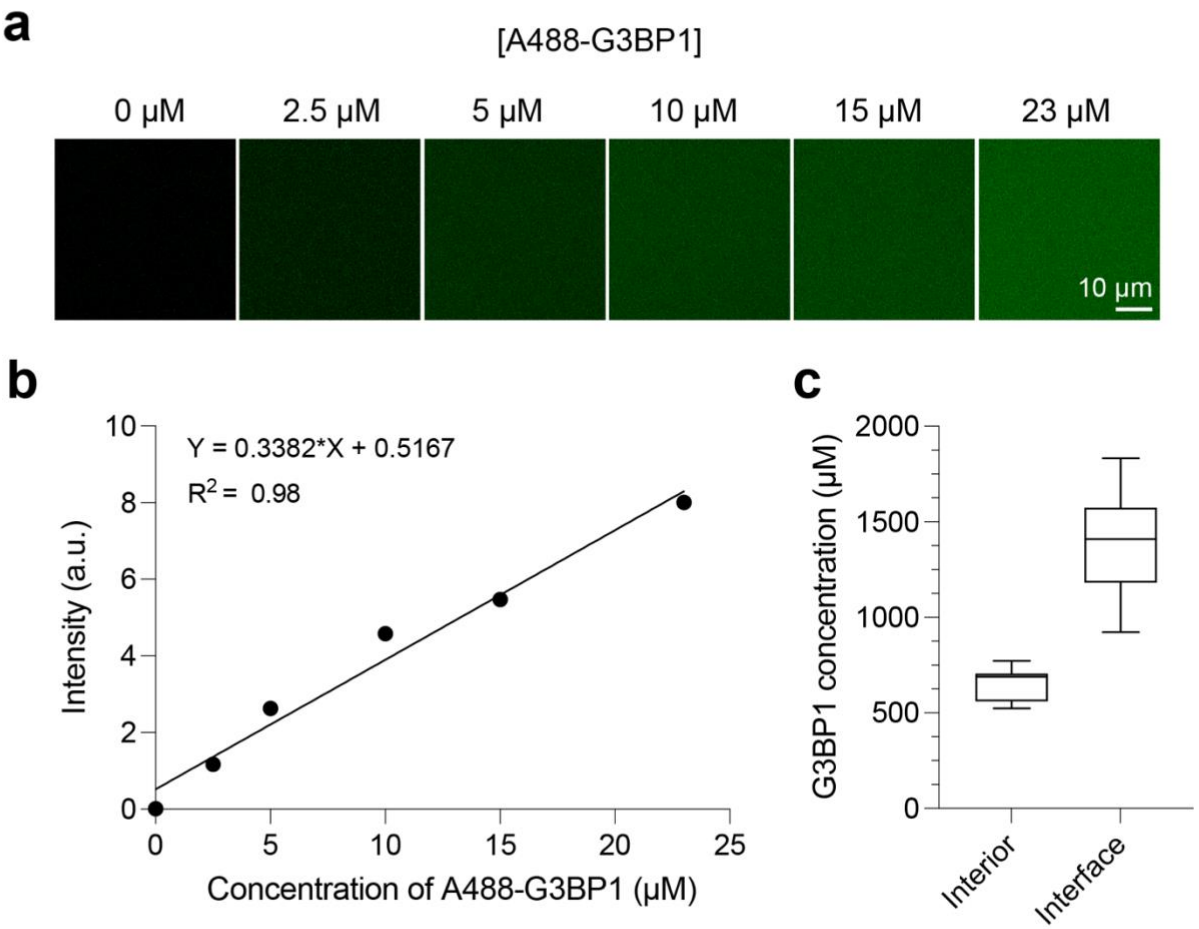


**Supplementary Figure 34.** (a) Fluorescence images of Alexa488-labeled G3BP1 (A488-G3BP1) in a buffer composed of 25 mM Tris-HCl (pH 7.5), 25 mM NaCl, and 20 mM DTT. These images were used for generating a calibration curve, which is reported in (b). The equation of the fitted line and the $R^2$ value are reported. (c) The concentration of G3BP1 in condensates is estimated through interpolation with the calibration curve reported in (b), using fluorescence readings of A488-G3BP1 in condensates (interior and interface) composed of 10 µM G3BP1, 1 mg/ml $(TERRA)_{10}$ (corresponds to 50.7 µM), 5 mg/ml RGG, and 1.5 mg/ml $d(T)_{40}$ in a buffer containing 25 mM Tris-HCl (pH 7.5), 25 mM NaCl, 20 mM DTT, and 160 nM A488-G3BP1. The center line of the box plot represents the median and the whiskers indicate the full range of the data from the minimum to the maximum. Each measurement was independently repeated three times.

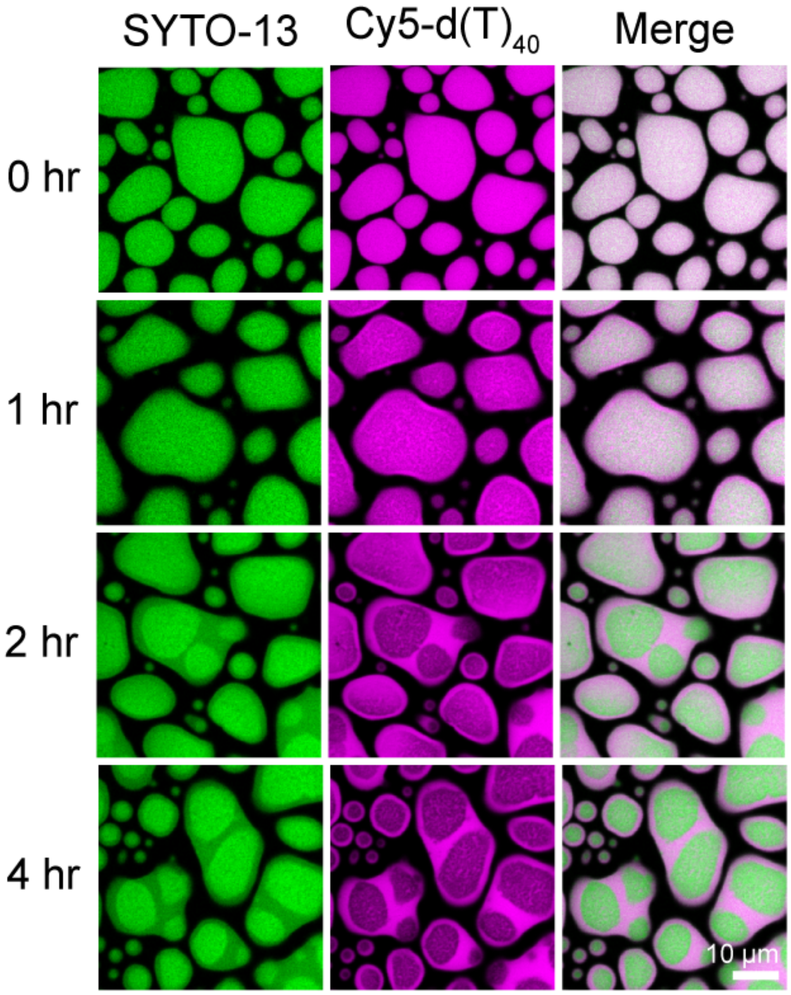


**Supplementary Figure 35.** Fluorescence images utilizing SYTO-13 and Cy5-d(T)$_{40}$ of (TERRA)$_{10}$ containing RGG-d(T)$_{40}$ condensates along with 17-fold diluted G3BP1 buffer [50 mM HEPES (pH 7.5) and 150 mM NaCl] without G3BP1 protein. The composition of the condensates is 1.0 mg/ml (TERRA)$_{10}$ (corresponds to 50.7 μM), 5.0 mg/ml RGG, 1.5 mg/ml d(T)$_{40}$ in a buffer containing 25 mM Tris-HCl (pH 7.5), 25 mM NaCl, 20 mM DTT. The concentration range of labeled components is 250 nM to 500 nM. This experiment was independently repeated two times.

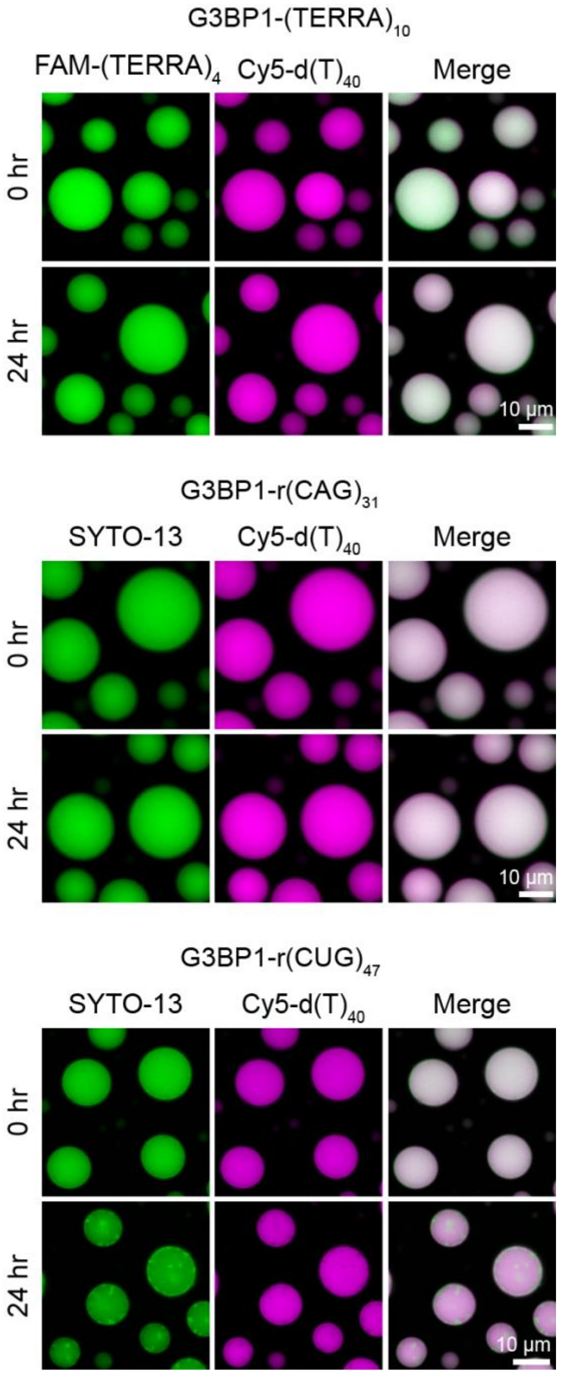


**Supplementary Figure 36.** Fluorescence images utilizing FAM-(TERRA)$_4$ or SYTO-13 and Cy5-d(T)$_{40}$ of RGG-d(T)$_{40}$ condensates containing (TERRA)$_{10}$, r(CAG)$_{31}$, or r(CUG)$_{47}$ repeat RNAs along with G3BP1. These observations correspond to data shown in **Fig. 6**. The composition of the condensate system used here is 10 μM G3BP1, 1 mg/ml RNA [0.45 mg/ml in the case of r(CUG)$_{47}$; (TERRA)$_{10}$, 101 μM, r(CAG)$_{31}$, 33 μM; r(CUG)$_{47}$, 10 μM], 5 mg/ml RGG, and 1.5 mg/ml d(T)$_{40}$ in a buffer containing 25 mM Tris-HCl (pH 7.5), 25 mM NaCl, and 20 mM DTT. The concentration range of labeled components is 250 nM to 500 nM. Each measurement was independently repeated three times.

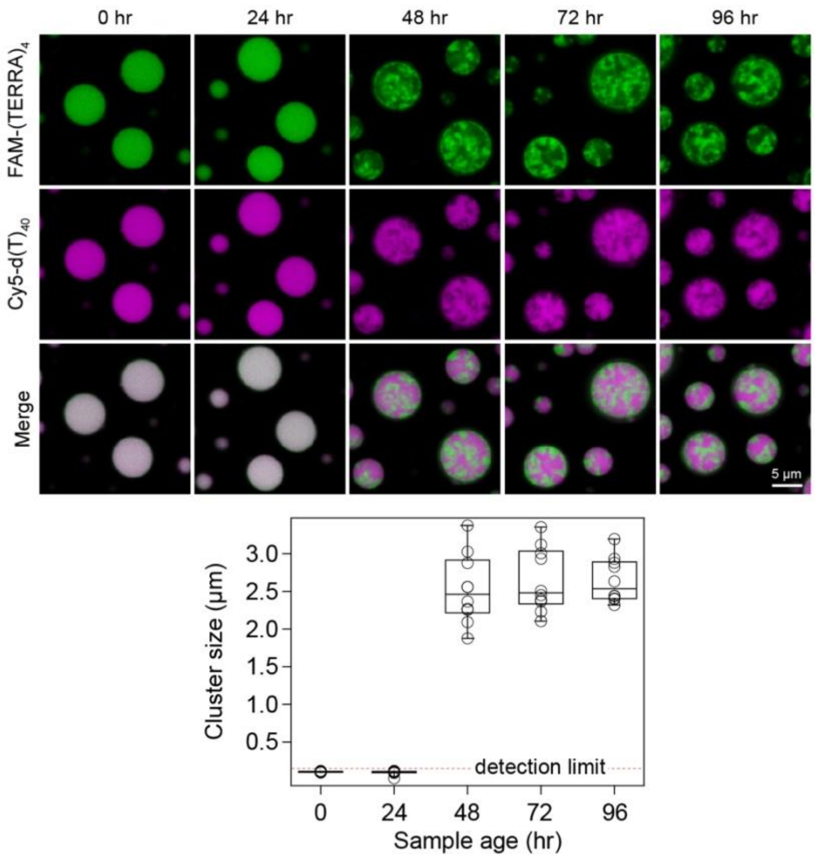


**Supplementary Figure 37.** (top) Fluorescence images utilizing FAM-$(TERRA)_4$ and Cy5-$d(T)_{40}$ of RGG-$d(T)_{40}$ condensates containing $(TERRA)_{10}$ along with G3BP1. (bottom) Cluster sizes derived from SAC are reported. The detection limit of SAC is demarcated. These observations correspond to data shown in **Fig. 6**. The composition of the condensate system used here is 10 μM G3BP1, 1 mg/ml RNA (corresponds to 50.7 μM), 5 mg/ml RGG, and 1.5 mg/ml $d(T)_{40}$ in a buffer containing 25 mM Tris-HCl (pH 7.5), 25 mM NaCl, and 20 mM DTT. The concentration of labeled components is 250 nM. Each measurement was independently repeated three times.

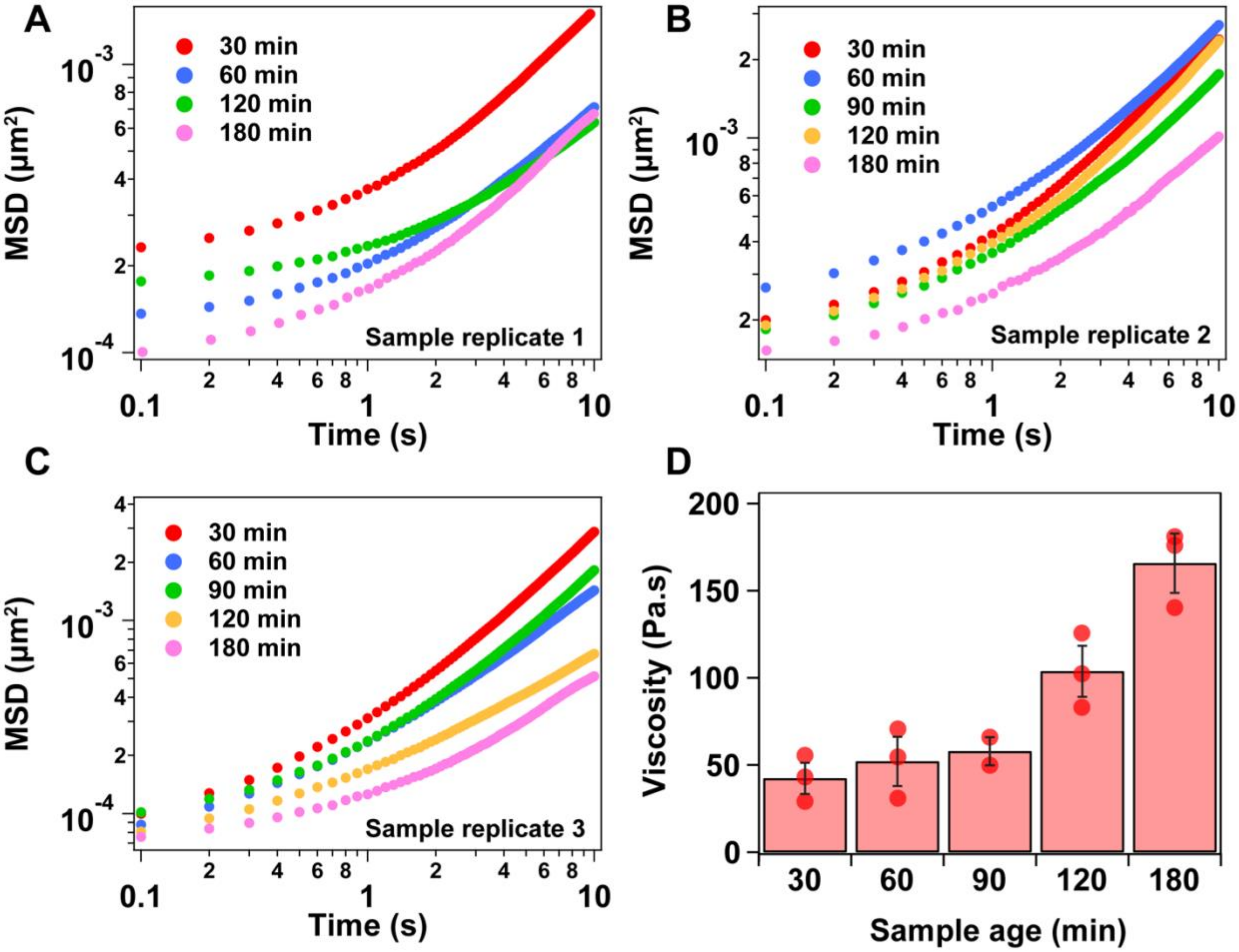


**Supplementary Figure 38.** (a-c) MSD plots for three replicates of $(TERRA)_{10}$ containing RGG-d$(T)_{40}$ condensates with G3BP1. (d) Estimated viscosities from the MSDs shown in (a-c), which is also reported in **Fig. 6j**. Error bars denote the standard deviation.

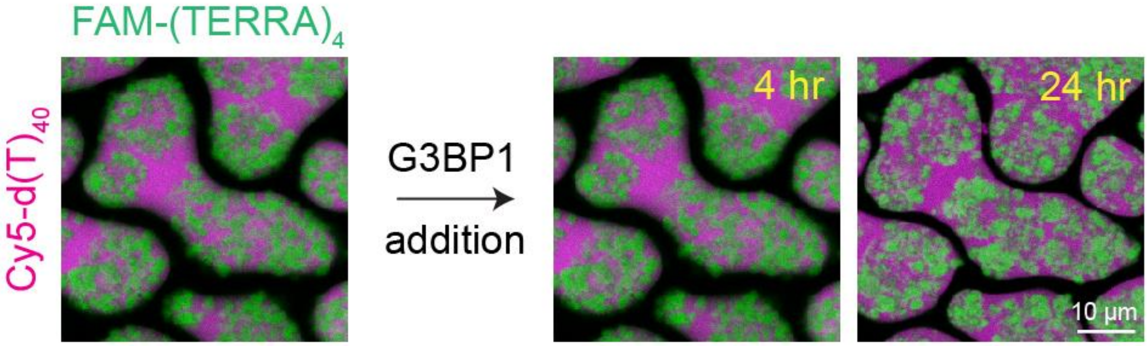


**Supplementary Figure 39.** Addition of G3BP1 to $(TERRA)_{10}$ containing RGG-d$(T)_{40}$ condensates after the formation of RNA clusters at 5 hours since sample preparation. The composition of the ternary condensate system is 0.5 mg/ml RNA (corresponds to 25.4 μM), 2.5 mg/ml RGG, and 0.75 mg/ml d$(T)_{40}$. The buffer composition is 25 mM Tris-HCl (pH 7.5), 25 mM NaCl, and 20 mM DTT. The concentration of G3BP1 used is 10 μM. The concentration of labeled components is 250 nM. Each measurement was independently repeated three times.

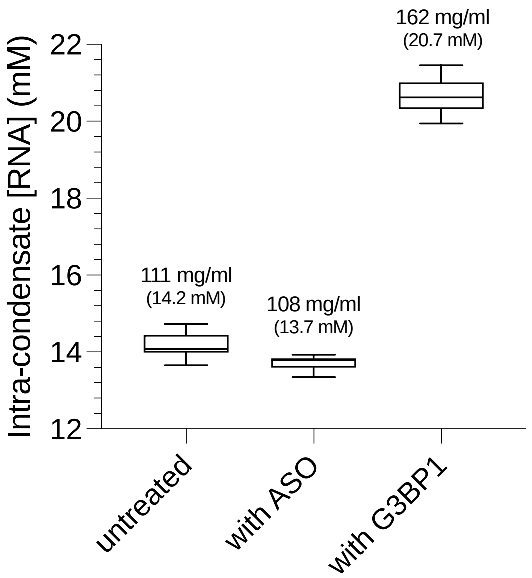


**Supplementary Figure 40.** Estimation of intra-condensate RNA concentration in RGG-$(TERRA)_4$-$d(T)_{40}$ condensates in the absence or presence of ASO or G3BP1. The 'untreated' sample (similar data is shown in Supplementary Fig. 8) is composed of 1 mg/ml $(TERRA)_4$ (corresponds to 127 µM), 5 mg/ml RGG, and 1.5 mg/ml $d(T)_{40}$ in a buffer containing 25 mM Tris-HCl (pH 7.5), 25 mM NaCl, and 20 mM DTT with 180 nM FAM labeled $(TERRA)_4$. In 'with ASO' and 'with G3BP1' samples, 1 mg/ml ASO [r(CCCUAA)] and 10 µM G3BP1 were included, respectively, while all other components were kept unchanged with the 'untreated' sample conditions. The mean intra-condensate RNA concentration is reported in mg/ml and molarity within the plot. The center line of the box plot represents the median, and the whiskers indicate the full range of the data from the minimum to the maximum. Each measurement was independently repeated three times.

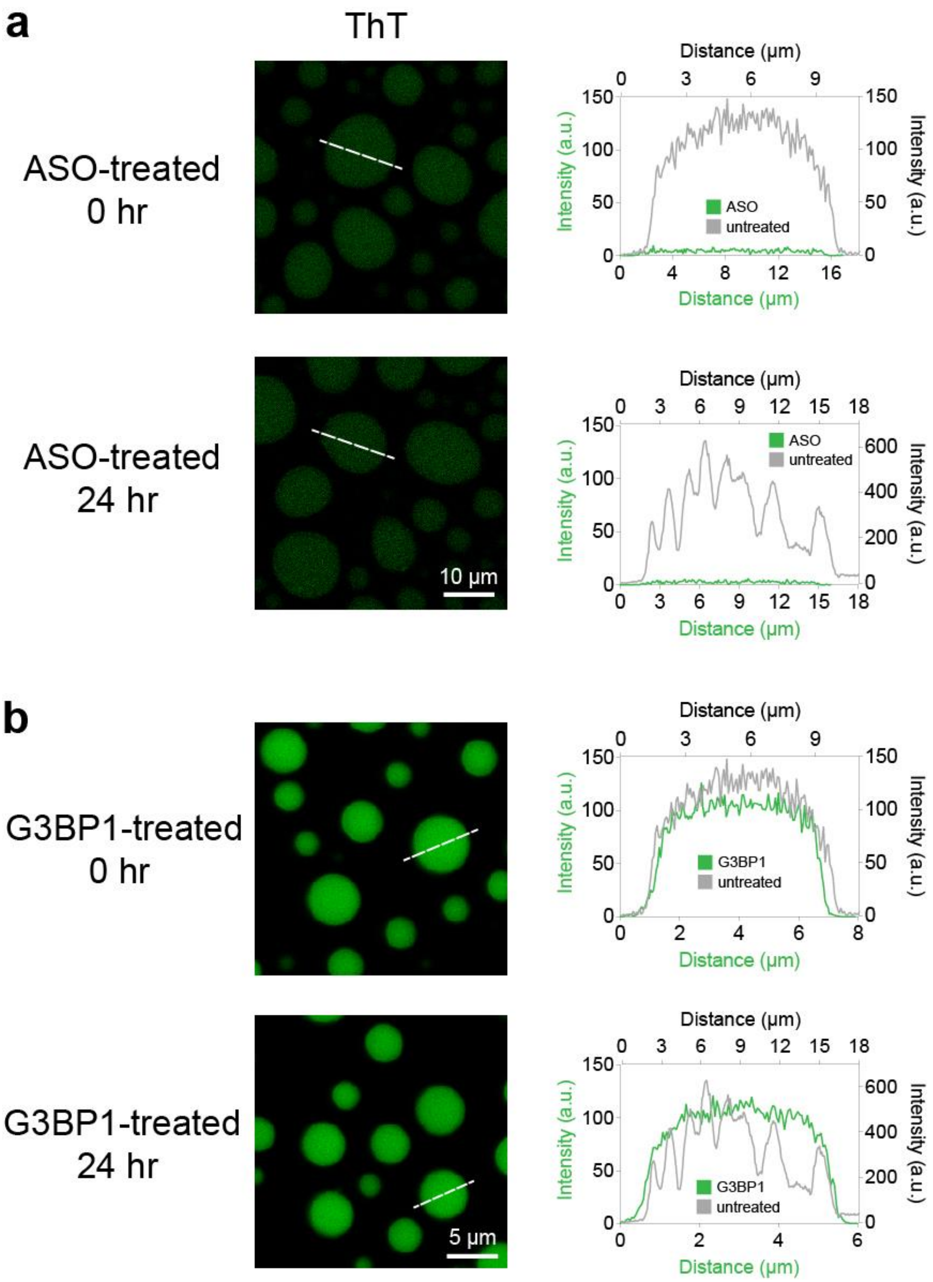


**Supplementary Figure 41.** Thioflavin T (ThT) staining of $(TERRA)_{10}$ containing RGG-d$(T)_{40}$ treated either with (a) ASO [sequence: r(CCCUAA)] or (b) G3BP1 at initial (15 minutes after sample preparation) and late (24 hours after sample preparation) time points (shown in green) along with ThT intensity profiles of untreated ternary condensates (shown in grey; data is taken from **Fig. 2h**). The composition of the quaternary condensate system is 1 mg/ml RNA (corresponds to 50.7 μM), 5 mg/ml RGG, and 1.5 mg/ml d$(T)_{40}$ with either 1 mg/ml ASO or 10 μM G3BP1. The buffer composition is 25 mM Tris-HCl (pH 7.5), 25 mM NaCl, and 20 mM DTT. The concentration of ThT used is 50 μM. Each measurement was independently repeated three times.

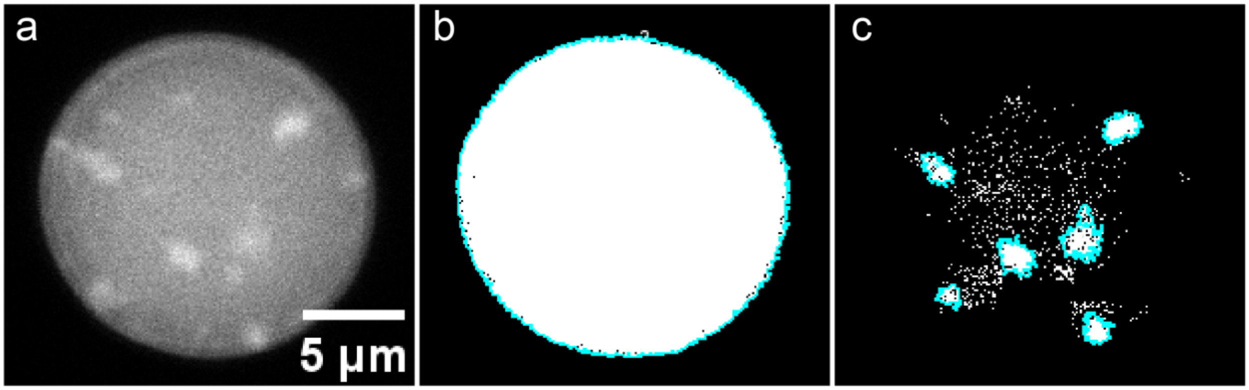


**Supplementary Figure 42.** (a) Greyscale image of SYTO-13 stained r(CUG)$_{47}$ clusters in RGG-d(T)$_{40}$ condensates containing G3BP1 at 24 hours after sample preparation. The image shown here highlights two different phases of r(CUG)$_{47}$ inside RGG-d(T)$_{40}$ condensates: cluster and homogeneous. (b) Binary image generated by thresholding (a), highlighting entire condensate. (c) Binary image after thresholding (a) showing r(CUG)$_{47}$ clusters. The blue outline in (b) and (c) depicts the boundaries of the detected condensate and clusters, respectively. These observations correspond to data shown in **Fig. 6m, n**. The composition of the condensate system used here is 10 µM G3BP1, 0.45 mg/ml r(CUG)$_{47}$ (corresponds to 10 µM), 5 mg/ml RGG, and 1.5 mg/ml d(T)$_{40}$ in a buffer containing 25 mM Tris-HCl (pH 7.5), 25 mM NaCl, and 20 mM DTT. The concentration of the labeled component is 500 nM. The analysis reported here was performed using 6 condensates from 2 independently prepared samples.

**Supplementary Table 1**. Sequence details of the RNAs used in the study.

| S. no. | Name | Sequence (5' to 3') | Source |
|---|---|---|---|
| 1. | $(TERRA)_1$ | UUAGGG | IDT |
| 2. | $(TERRA)_4$ | UUAGGGUUAGGGUUAGGGUUAGGG | IDT |
| 3. | $(TERRA)_6$ | UUAGGGUUAGGGUUAGGGUUAGGGUUAGGGUUAG GG | IDT |
| 4. | $(TERRA)_8$ | UUAGGGUUAGGGUUAGGGUUAGGGUUAGGGUUAG GGUUAGGGUUAGGG | IDT |
| 5. | $(TERRA)_9$ | UUAGGGUUAGGGUUAGGGUUAGGGUUAGGGUUAG GGUUAGGGUUAGGGUUAGGG | IDT |
| 6. | $(TERRA)_{10}$ | UUAGGGUUAGGGUUAGGGUUAGGGUUAGGGUUAG GGUUAGGGUUAGGGUUAGGGUUAGGG | IDT |
| 7. | $(mut\text{-}TERRA)_{10}$ | UUAGUGUUAGUGUUAGUGUUAGUGUUAGUGUUAGU GUUAGUGUUAGUGUUAGUGUUAGUG | IDT |
| 8. | $(mut^{6GtoU}\text{-}TERRA)_{10}$ | UUAGUGUUAGGGUUAGUGUUAGUGUUAGGGUUAG GGUUAGUGUUAGUGUUAGGGUUAGUG | IDT |
| 9. | $r(CAG)_{31}$ | CAGCAGCAGCAGCAGCAGCAGCAGCAGCAGCAGCA GCAGCAGCAGCAGCAGCAGCAGCAGCAGCAGCAGC AGCAGCAGCAGCAGCAGCAGCAG | IDT |
| 10. | $r(CUG)_{31}$ | CUGCUGCUGCUGCUGCUGCUGCUGCUGCUGCUGC UGCUGCUGCUGCUGCUGCUGCUGCUGCUGCUGCU GCUGCUGCUGCUGCUGCUGCUGCUG | IDT |
| 11. | $r(CAG)_{20}$ | CAGCAGCAGCAGCAGCAGCAGCAGCAGCAGCAGCA GCAGCAGCAGCAGCAGCAGCAGCAG | IDT |
| 12. | $r(CUG)_{47}$ | CUGCUGCUGCUGCUGCUGCUGCUGCUGCUGCUGC UGCUGCUGCUGCUGCUGCUGCUGCUGCUGCUGCU GCUGCUGCUGCUGCUGCUGCUGCUGCUGCUGCUG CUGCUGCUGCUGCUGCUGCUGCUGCUGCUGCUGC UGCUG | IVT |
| 13. | $r(GGGGCC)_5$ | GGGGCCGGGGCCGGGGCCGGGGCCGGGGCC | IDT |

**Supplementary Table 2**. Sequence details of the antisense RNA oligonucleotides (ASOs) used in the study.

| S. no. | Name | Sequence (5' to 3') | Source |
|---|---|---|---|
| 1. | Antisense TERRA | CCCUAA | IDT |
| 2. | Scrambled ASO | CACUAC | IDT |

### Supplementary Video Legends

**Supplementary Video 1**. Time-lapse fluorescence imaging showing cluster formation in RGG-d(T)$_{40}$ condensates containing (TERRA)$_{10}$, as visualized by SYTO-13 fluorescence. The composition of the condensate system used here is 1 mg/ml (TERRA)$_{10}$, 5 mg/ml RGG, and 1.5 mg/ml d(T)$_{40}$ in a buffer containing 25 mM Tris-HCl (pH 7.5), 25 mM NaCl, and 20 mM DTT. The concentration of the labeled component is 500 nM.

**Supplementary Video 2.** Addition of 1.5 mg/ml d(T)$_{40}$, doped with Cy5 labeled d(T)$_{40}$, to RGG-(TERRA)$_{10}$ condensates visualized with FAM-(TERRA)$_4$ fluorescence. The composition of the binary condensate system used here is 1 mg/ml (TERRA)$_{10}$ and 5 mg/ml RGG in a buffer containing 25 mM Tris-HCl (pH 7.5), 25 mM NaCl, and 20 mM DTT. The concentration of the labeled components is 250 nM.

**Supplementary Video 3.** Temperature-controlled microscopy showing (TERRA)$_{10}$ phase separation coupled to percolation. 1 mg/ml (TERRA)$_{10}$ in a buffer composed of 50 mM HEPES (pH 7.5), 6.25 mM $Mg^{2+}$.

**Supplementary Video 4.** Temperature-controlled microscopy showing shape relaxation and persistence of percolated (TERRA)$_{10}$ condensates; 1 mg/ml (TERRA)$_{10}$ in buffer composed of 50 mM HEPES (pH 7.5), 10 mM $Mg^{2+}$.

**Supplementary Video 5.** Temperature-controlled microscopy showing reversible phase separation of (mut-TERRA)$_{10}$; 1 mg/ml (mut-TERRA)$_{10}$ in buffer composed of 50 mM HEPES pH 7.5, 25 mM $Mg^{2+}$.

**Supplementary Video 6.** Active fusion of (TERRA)$_{10}$ containing RGG-d(T)$_{40}$ condensates at 20 minutes of age visualized with FAM-(TERRA)$_4$ and Cy5-d(T)$_{40}$ fluorescence. The composition of the condensate system used here is 1 mg/ml (TERRA)$_{10}$, 5 mg/ml RGG, and 1.5 mg/ml d(T)$_{40}$ in a buffer containing 25 mM Tris-HCl (pH 7.5), 25 mM NaCl, and 20 mM DTT. The concentration of the labeled components is 250 nM.

**Supplementary Video 7.** Active fusion of (TERRA)$_{10}$ containing RGG-d(T)$_{40}$ condensates at 150 minutes of age visualized with FAM-(TERRA)$_4$ and Cy5-d(T)$_{40}$ fluorescence. The composition of the condensate system used here is 1 mg/ml (TERRA)$_{10}$, 5 mg/ml RGG, and 1.5 mg/ml d(T)$_{40}$ in a buffer containing 25 mM Tris-HCl (pH 7.5), 25 mM NaCl, and 20 mM DTT. The concentration of the labeled components is 250 nM.

**Supplementary Video 8.** Dissolution of the condensate shell but not the inner RNA clusters using 455 mM NaCl in a 6-hour aged RGG-d(T)$_{40}$ condensate containing (TERRA)$_{10}$, visualized with FAM-(TERRA)$_4$ and Cy5-d(T)$_{40}$ fluorescence. The composition of the condensate system used here is 0.5 mg/ml (TERRA)$_{10}$, 2.5 mg/ml RGG, and 0.75 mg/ml d(T)$_{40}$ in a buffer containing 25 mM Tris-HCl (pH 7.5), 25 mM NaCl, and 20 mM DTT. The concentration of the labeled components is 250 nM.

**Supplementary Video 9.** Dissolution of the condensate shell but not the inner RNA clusters using 455 mM NaCl in a 24-hour aged RGG-d(T)$_{40}$ condensate containing (TERRA)$_{10}$, visualized by FAM-(TERRA)$_4$ and Cy5-d(T)$_{40}$ fluorescence. The composition of the condensate system used here is 1 mg/ml (TERRA)$_{10}$, 5 mg/ml RGG, and 1.5 mg/ml d(T)$_{40}$ in a buffer containing 25 mM Tris-HCl (pH 7.5), 25 mM NaCl, and 20 mM DTT. The concentration of the labeled components is 250 nM.

**Supplementary Video 10.** Dissolution of freshly prepared (15 minutes of age) RGG-d(T)$_{40}$ condensates containing (TERRA)$_{10}$ using 455 mM NaCl, visualized with FAM-(TERRA)$_4$ and Cy5-d(T)$_{40}$ fluorescence. The composition of the condensate system used here is 0.5 mg/ml (TERRA)$_{10}$, 2.5 mg/ml RGG, and 0.75 mg/ml d(T)$_{40}$ in a buffer containing 25 mM Tris-HCl (pH 7.5), 25 mM NaCl, and 20 mM DTT. The concentration of the labeled components is 250 nM.

**Supplementary Video 11.** FRAP of FAM-$(TERRA)_4$ at the 0-hour time point in $(TERRA)_4$ containing RGG-$d(T)_{40}$ condensates. The scale bar is 10 µm. The composition of the condensate system used here is 1 mg/ml $(TERRA)_4$, 5 mg/ml RGG, and 1.5 mg/ml $d(T)_{40}$ in a buffer containing 25 mM Tris-HCl (pH 7.5), 25 mM NaCl, and 20 mM DTT. The concentration of the labeled component is 250 nM.

**Supplementary Video 12.** FRAP of A594-RGG at the 0-hour time point in $(TERRA)_4$ containing RGG-$d(T)_{40}$ condensates. The scale bar is 10 µm. The composition of the condensate system used here is 1 mg/ml $(TERRA)_4$, 5 mg/ml RGG, and 1.5 mg/ml $d(T)_{40}$ in a buffer containing 25 mM Tris-HCl (pH 7.5), 25 mM NaCl, and 20 mM DTT. The concentration of the labeled component is 250 nM.

**Supplementary Video 13.** FRAP of Cy5-$d(T)_{40}$ at the 0-hour time point in $(TERRA)_4$ containing RGG-$d(T)_{40}$ condensates. The scale bar is 10 µm. The composition of the condensate system used here is 1 mg/ml $(TERRA)_4$, 5 mg/ml RGG, and 1.5 mg/ml $d(T)_{40}$ in a buffer containing 25 mM Tris-HCl (pH 7.5), 25 mM NaCl, and 20 mM DTT. The concentration of the labeled component is 250 nM.

**Supplementary Video 14.** FRAP of FAM-$(TERRA)_4$ at the 8-hour time point in $(TERRA)_4$ containing RGG-$d(T)_{40}$ condensates. The scale bar is 10 µm. The composition of the condensate system used here is 1 mg/ml $(TERRA)_4$, 5 mg/ml RGG, and 1.5 mg/ml $d(T)_{40}$ in a buffer containing 25 mM Tris-HCl (pH 7.5), 25 mM NaCl, and 20 mM DTT. The concentration of the labeled component is 250 nM.

**Supplementary Video 15.** FRAP of A594-RGG at the 8-hour time point in $(TERRA)_4$ containing RGG-$d(T)_{40}$ condensates. The scale bar is 10 µm. The composition of the condensate system used here is 1 mg/ml $(TERRA)_4$, 5 mg/ml RGG, and 1.5 mg/ml $d(T)_{40}$ in a buffer containing 25 mM Tris-HCl (pH 7.5), 25 mM NaCl, and 20 mM DTT. The concentration of the labeled component is 250 nM.

**Supplementary Video 16.** FRAP of Cy5-$d(T)_{40}$ at the 8-hour time point in $(TERRA)_4$ containing RGG-$d(T)_{40}$ condensates. The scale bar is 10 µm. The composition of the condensate system used here is 1 mg/ml $(TERRA)_4$, 5 mg/ml RGG, and 1.5 mg/ml $d(T)_{40}$ in a buffer containing 25 mM Tris-HCl (pH 7.5), 25 mM NaCl, and 20 mM DTT. The concentration of the labeled component is 250 nM.

**Supplementary Video 17.** VPT of 200 nm beads inside $(TERRA)_{10}$ containing RGG-$d(T)_{40}$ condensates at 15-minute time point after sample preparation. The composition of the condensate system used here is 2 mg/ml $(TERRA)_{10}$, 10 mg/ml RGG, and 3 mg/ml $d(T)_{40}$ in a buffer containing 25 mM Tris-HCl (pH 7.5), 25 mM NaCl, and 20 mM DTT.

**Supplementary Video 18.** VPT of 200 nm beads inside $(TERRA)_{10}$ containing RGG-$d(T)_{40}$ condensates at 45-minute time point after sample preparation. The composition of the condensate system used here is 2 mg/ml $(TERRA)_{10}$, 10 mg/ml RGG, and 3 mg/ml $d(T)_{40}$ in a buffer containing 25 mM Tris-HCl (pH 7.5), 25 mM NaCl, and 20 mM DTT.

**Supplementary Video 19.** VPT of 200 nm beads inside $(TERRA)_{10}$ containing RGG-$d(T)_{40}$ condensates at 150-minute time point after sample preparation. The composition of the condensate system used here is 2 mg/ml $(TERRA)_{10}$, 10 mg/ml RGG, and 3 mg/ml $d(T)_{40}$ in a buffer containing 25 mM Tris-HCl (pH 7.5), 25 mM NaCl, and 20 mM DTT.

**Supplementary Video 20.** Addition of 1 mg/ml ASO [r(CCCUAA)] to 5-hour aged RGG-$d(T)_{40}$ condensates with RNA clusters of $(TERRA)_{10}$, visualized with FAM-$(TERRA)_4$ and Cy5-$d(T)_{40}$ fluorescence. The composition of the condensate system used here is 0.5 mg/ml $(TERRA)_{10}$, 2.5 mg/ml RGG, and 0.75 mg/ml $d(T)_{40}$ in a buffer containing 25 mM Tris-HCl (pH 7.5), 25 mM NaCl, and 20 mM DTT. The concentration of the labeled components is 250 nM.

**Supplementary Video 21.** Addition of 1 mg/ml ASO [r(CCCUAA)] to 18-hour aged RGG-$d(T)_{40}$ condensates with RNA clusters of $(TERRA)_{10}$, visualized with FAM-$(TERRA)_4$ and Cy5-$d(T)_{40}$

fluorescence. The composition of the condensate system used here is 0.5 mg/ml $(TERRA)_{10}$, 2.5 mg/ml RGG, and 0.75 mg/ml $d(T)_{40}$ in a buffer containing 25 mM Tris-HCl (pH 7.5), 25 mM NaCl, and 20 mM DTT. The concentration of the labeled components is 250 nM.
